# Light-directed evolution of dynamic, multi-state, and computational protein functionalities

**DOI:** 10.1101/2024.02.28.582517

**Authors:** Vojislav Gligorovski, Marco Labagnara, Lorenzo Scutteri, Marius Blackholm, Andreas Möglich, Nahal Mansouri, Sahand Jamal Rahi

**Affiliations:** Laboratory of the Physics of Biological Systems, Institute of Physics, EPFL, Switzerland; Laboratory of Protein and Cell Engineering, Institute of Bioengineering, EPFL, Switzerland; Department of Biochemistry, Universität Bayreuth, Germany; Division of Pulmonary Medicine, Department of Medicine, Lausanne University Hospital (CHUV), University of Lausanne (UNIL), Switzerland

## Abstract

Directed evolution is a powerful method in biological engineering. Current approaches draw on time-invariant selection mechanisms, ideal for evolving steady-state properties such as enzymatic activity or fluorescence intensity. A fundamental problem remains how to continuously evolve dynamic, multi-state, or computational functionalities, e.g., on-off kinetics, state-specific activity, stimulus-responsiveness, or switching and logic capabilities. These require selection pressure on all of the states of a protein of interest (POI) and the transitions between them. We realized that optogenetics and cell cycle oscillations could be leveraged for a novel directed evolution paradigm (‘optovolution’) that is germane for this need: We designed a signaling cascade in budding yeast where optogenetic input switches the POI between off (0) and on (1) states. In turn, the POI controls a Cdk1 cyclin, which in the re-engineered cell cycle system is essential for one cell cycle stage but poisonous for another. Thus, the cyclin must oscillate (1-0-1-0…) for cell proliferation. In this system, evolution can act efficiently on the POI’s different states, input-output relations, and dynamics on the timescale of minutes in every cell cycle. Further, controlling the pacemaker, light, directs and tunes selection pressures. Optovolution is *in vivo*, continuous, self-selecting, and efficient. We first evolved two optogenetic systems, which relay 0/1 input to 0/1 output: We obtained 19 new variants of the LOV transcription factor El222 that were stronger, less leaky, or green light responsive *in vivo*. We demonstrate the utility of the latter mutations for orthogonal color-multiplexing with only LOV domains for the first time. Evolving the PhyB-Pif3 optogenetic system, we discovered that loss of *YOR1* makes supplementing the chromophore phycocyanobilin (PCB) unnecessary. Finally, we demonstrate the generality of the method by evolving a destabilized rtTA transcription factor, which performs an AND operation between transcriptional and doxycycline input. Optovolution makes coveted, difficult-to-change protein functionalities continuously evolvable.

## Introduction

Directed evolution has been critical to improving and diversifying many of the molecular tools currently used in research and industry such as enzymes^1–5^, fluorescent proteins^6,7^, and antibodies^8,9^. The method relies on mutations from global or targeted mutagenesis^10–12^ to diversify genetically encoded molecules. These are then screened for a desired function. The mutation-screen cycle is generally repeated to bring the molecules’ functionalities closer to the desired goal.^13^

A key innovation in the field has been to couple a molecule of interest’s functionality to organismal fitness obviating external intervention and screening. Such ‘continuous’ directed evolution methods rely on *in vivo* self-selection, which substantially simplifies and speeds up experiments, facilitating scale-up.^1,14–18^ Current continuous directed evolution methods, however, use time-invariant selection mechanisms based on, for example, the strength of binding or of transcriptional activation, which are steady-state functionalities. These mechanisms can be combined to achieve complex but non-dynamic functionalities by simultaneously selecting for and against different goals^19–27^ (see **Supplementary Note 1** for a partial list of previous continuous multiple-goal directed evolution experiments).

Thus, existing continuous directed evolution paradigms are poorly suited for the many natural and synthetic biological molecules that can be in different states and can dynamically switch between them. In fact, all signal transduction, control, and computation necessarily involve changes between states, usually, on (1) or off (0). For example, many transcription factors and kinases are controlled by upstream signals. In addition to relaying signals, proteins and circuits often integrate multiple inputs for cellular decisions, e.g., the amount of DNA damage and arrest times jointly determine the DNA damage checkpoint override decision.^28^ Such computational capabilities are highly desirable. Building cells with more sophisticated decision-making capabilities needs logic gating, which has been created by rational design.^29–33^ Evolving such proteins requires, in principle, that selection pressure is simultaneously applied on all of the computations, that is, all input-output relationships of the POI. Otherwise, if directed evolution is performed for only one state, the other states or the ability to switch to them can be corrupted.^34^ For example, long-term directed evolution of GPCRs under step-wise constant ligand concentrations reduces ligand responsiveness.^35^

To address this need, researchers have developed non-continuous directed evolution methods, in which time-invariant selection mechanisms are alternated to screen each POI state sequentially^36–70^ (see **Supplementary Note 2** for a partial list of past alternating selection/counterselection experiments). In these methods, two steps are typically repeated: an assay for on (1) activity is performed under inducing conditions, then an assay for off (0) activity under non-inducing conditions. In practice, selection and counterselection are performed with alternating media conditions, e.g., -uracil and +5-fluoroorotic acid^71^, or changing fluorescence-activated cell sorting (FACS) gates. While creative, these approaches require interventions involving labor or specialized instrumentation or both. Furthermore, one cycle of selection and counterselection based on media switches and cell proliferation took 8-64 hrs with bacteria or 2-5 days with yeast. This timescale determines the selection pressure placed specifically on the kinetics of the POI switch cycle, which is typically desired to be much shorter.

The ability to perform directed evolution for dynamic, multi-state, and computational protein functionalities would advance biological engineering. Here, we present the first such paradigm, which we name ‘optovolution’. In contrast to previous approaches, optovolution was designed with a mechanism to evolve all input-output relationships of a POI as well as switches among them continuously. Two conceptual advances underlie optovolution (**Discussion**): By engineering a POI to control a cell cycle regulator that is essential for one step of cell proliferation but toxic to another, the POI must evolve or maintain the ability to switch the output between 0 to 1 and back rapidly, once per cell cycle. The selection mechanism is cell-intrinsic and inherently dynamic. Secondly, by controlling the input to the POI with light and optogenetics, the selection pressure is shaped by an external signal that is changing at the timescale of a cell cycle.

We apply optovolution to three systems, the optogenetic transcriptional system El222 and the optogenetic protein-protein interaction system PhyB-Pif3, which relay light input by switching on (1) or off (0), and an AND gate, in which a degron-tagged rtTA transcription factor is controlled by transcription and doxycycline. The dynamic, multi-state properties of these proteins make them unsuitable for existing continuous directed evolution paradigms:

1. LOV domains make up the core of most non-ion-channel optogenetic systems^72–80,80–86^, presumably because they are small, reliable, and do not need exogenous chromophores. Interest in LOV domains has spurred past diversification work^54,87–89^. While usually controlled with light, LOV domains have also been developed as temperature sensors^90,91^. The LOV transcription factor El222, in particular, is widely used in many biological models from cell-free systems^92^, to bacteria^93^, yeast^94,95^, mammalian cell lines^96,97^, and zebrafish^98^. (See **Supplementary Note 3** for additional information regarding El222.) In comparison to other inducible transcriptional systems, El222 controlling the standard *light-inducible promoter* (*LIP*) has both more off-state leakiness and lower on-state strength than, for example, the *GAL1* promoter, demonstrating potential room for improvement.^88^ Less leaky or more light-sensitive variants are desirable for different applications: Low leakiness is necessary when controlling toxic genes such as endonucleases; on the other hand, higher output under less light allows reducing phototoxicity.^88^ Further, all existing LOV systems are thought to be activated exclusively by blue light, which limits their use for independently controlling multiple optogenetic systems within the same organism. Ideally, one could build optogenetic systems based on LOV domains, change their color responsiveness by mutation, and thus straightforwardly combine different systems. Instead, for independent multiplexing, one LOV domain protein has to be used together with other light-sensitive domains, each representing a challenging engineering endeavor.^99–102^ In fact, substantially changing the color responsiveness of LOV domains was conjectured to be “likely impossible” based on the lack of such known natural or engineered examples and physicochemical considerations.^103^ Previous work to shift the absorption spectrum of LOV domains with targeted mutations achieved a maximal shift of 8 nm further toward UV in the absorption peak *in vitro*^104^. Shifts in the emission spectrum for use as fluorescent proteins have also been attained.^105^ Using optovolution, we evolved 19 new variants of El222, including more light sensitive, less leaky, or *in vivo* green-light responsive mutants, paving the way for new applications. Furthermore, we built a circuit for green-light-only activation of transcription, demonstrating the possibility of independent addressing of LOV proteins with green and blue light.
2. In the widely used PhyB-Pif3 system, PhyB and Pif3 dimerize in response to red light, dissociate rapidly with far-red light, or more slowly in darkness. The system is often used to change the intracellular localization of proteins by fusing them to PhyB or Pif3 and anchoring the other dimerization partner to a specific intracellular compartment^106,107^. (See **Supplementary Note 4** for additional information regarding the PhyB-Pif3 system.) For transcriptional induction, PhyB and Pif3 are fused to the Gal4 transcriptional activation domain (TAD) and the Gal4 DNA binding domain (DBD), respectively.^108–110^ For use in yeast, *Caenorhabditis elegans*, and mammalian cells, the chromophore PCB has to be supplemented.^106,108,111^ The co-factor is expensive as well as light- and temperature-sensitive, complicating experiments. In previous work, a biochemical pathway to convert heme to PCB was introduced in different organisms.^111–114^ This involved substantial genetic manipulation of the host genome: Several transgenes had to be introduced, some of which needed to be targeted to mitochondria, their expression levels needed to be optimized, and, in certain cases, endogenous genes needed to be deleted. It is unclear whether there are pathways in cells that could be modified for PhyB-Pif3 to work without supplemental PCB. Using optovolution, we found that loss of function of *YOR1* makes PCB unnecessary in budding yeast. The use of *yor1Δ* substantially simplifies applications and further directed evolution experiments.
3. While optovolution leverages light for directing evolution, we demonstrate the more general applicability of optovolution by evolving a non-optogenetic logic gate. It is often desirable to condition the activity of molecules on two inputs. This requires AND logic, which is relatively difficult to create in biological systems. To construct such a system, we used the doxycyclin-controlled transcription factor rtTA^115^ and fused it to a PEST degron^116^. (See **Supplementary Note 5** for additional information regarding rtTA.) This double-lock control represents an AND gate function between transcriptional and doxycycline input. Without the degron, rtTA protein would be stable and thus, after translation, only controlled by doxycycline; responsiveness to further transcriptional input could only happen in proliferating cells at the relatively slow timescale of dilution. However, the PEST-rtTA fusion protein initially produced very weak output. With optovolution, we evolved two mutants that conferred substantial output from the *tetO* promoter while maintaining minimal leakiness.

Optovolution is the first continuous directed evolution paradigm for dynamic, multi-state, and computational functionalities. We demonstrate its utility by uncovering unexpected and useful mutations improving three different important molecular tools.

## Results

### Requirements for evolving dynamic, multi-state, and computational properties

We considered how to evolve a POI with one or more inputs and one output, each of which can be in the 0 or 1 state (Fig. 1 A). Specifically, there can be a single input *i* to the POI, for example, transcriptional induction of the POI, or a complex input vector ***i***=(*i_1_,…,i_n_*) such as specific phosphorylation patterns at *n* residues of the POI. For such a set of possible inputs and outputs, the main conceptual challenge was to find a mechanism to perform selection on each input-output relationship of the POI and the kinetics of the transitions. We suppose that the input vectors are labeled ***i_a_***, ***i_b_***, ***i_c_***, ***i_d_***, … and correspond to desired POI outputs 1, 0, 1, 0, … We realized that we could perform continuous directed evolution on this system if we made the POI fully control a cellular regulator that was essential for one step in cell proliferation but poisonous for another (Fig. 1 B, C). In such a system, if the POI switches between 1-0-1-0-… outputs, driven by repeated cycles of inputs ***i_a_***-***i_b_***-***i_c_***-***i_d_***-…, then cells will proliferate maximally (Fig. 1 D). Any error, either in switching speed or state, e.g., POI output sequence 1-1-1-0-…, causes skipped cell cycles, incurring a substantial fitness penalty and giving an advantage to mutants with better fidelity to the desired input-output relationships.

**Fig. 1.**
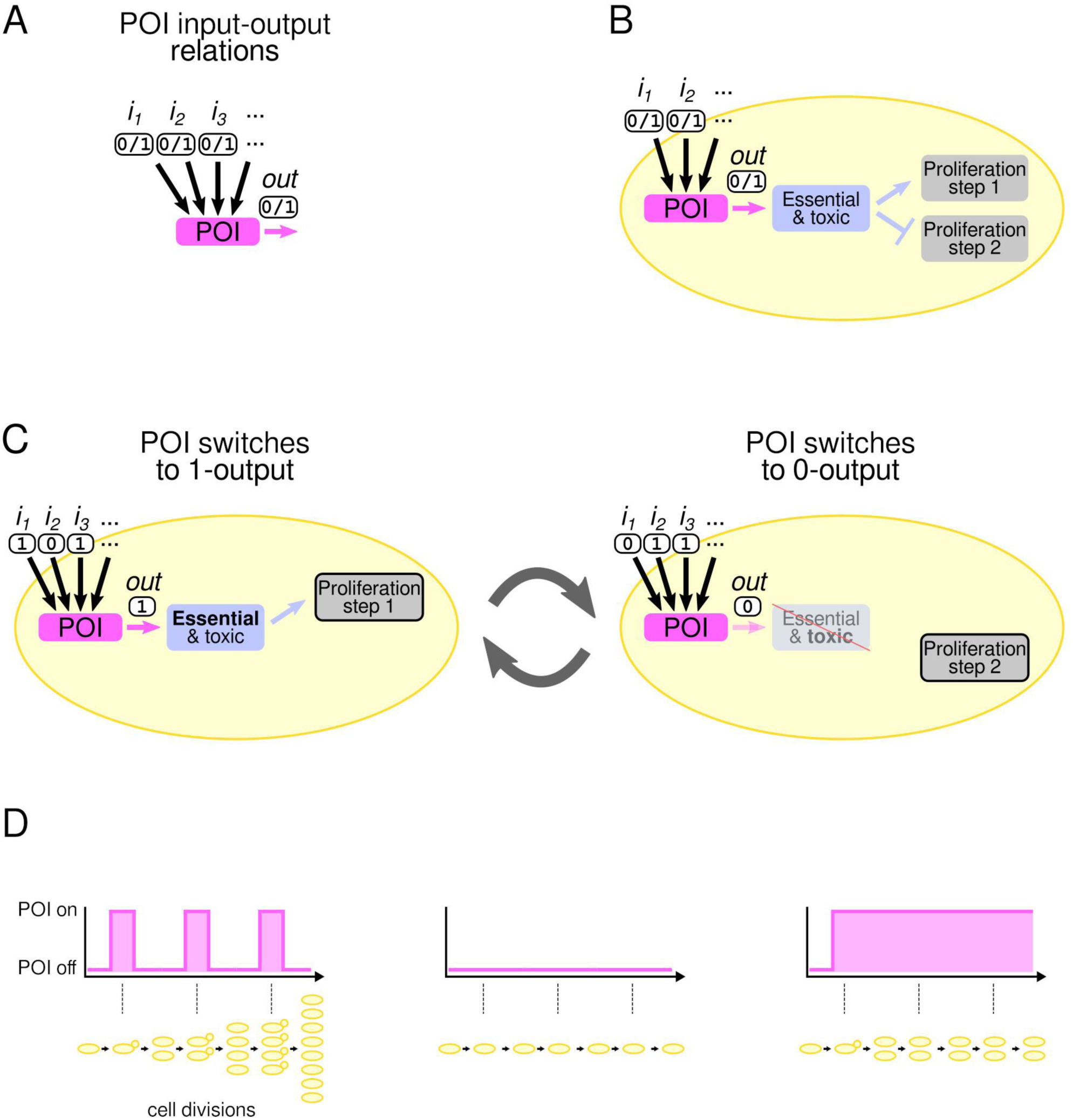
Conceptualization of optovolution. A. The POI has one or multiple inputs and one output. Inputs and output are idealized as having two states, on (1) or off (0). B. In optovolution, the POI controls a cellular regulator that is essential for one step in cellular proliferation but poisonous for another. The inputs to the POI are most conveniently controlled by light, mediated by optogenetic systems, which are not shown. C. By cycling the inputs to the POI, the POI switches output states and drives cycles of cellular proliferation. D. Cells only proliferate with repeated 1-0-1-0-… output pulses of the POI, not continuously on (1) or off (0) output, in a mutationally robust manner. (Cell death and size growth not shown.)

The inputs to the POI must be exogenously controllable. Optogenetic control of the POI inputs is compatible with continuous directed evolution since administering light, even for weeks, is precise, fast, easily tunable, cheap, straightforward, and scalable. For control of non-optogenetic POIs, light input can be indirectly mediated through the rapidly expanding optogenetics toolkit, which includes light-sensitive transcription factors, localization signals, degrons, and kinases, acting on the POI.^117–122^ On the other side, cell proliferation must be fully dependent on the POI’s 1-0-1-0-… output. The key idea was to find a regulator of cell proliferation, which would be required for one step in proliferation but block another (Fig. 1 B-D). By engineering the POI to fully control this regulator, the POI would drive one cell cycle for each 1-0 output oscillation of the POI.

### Implementation in budding yeast

We engineered the system in budding yeast because it has a relatively fast cell cycle (≈ 90 min period in glucose medium). A short generation time is important since it sets the maximum speed with which selection pressure can be exerted on POI state switching.

In search of a gene that is essential for one proliferative step but poisonous for another, we considered cell cycle control regulators that oscillate once per cell cycle. However, it is not obvious which to choose since the cell cycle control system is remarkably robust. For example, constitutive expression of *CLN2* and *CLB2* in the absence of all other Cdk1 cyclins (*cln1-3Δ clb1-6Δ*) suffices for cell cycles^123^. Even in such a highly stripped-down cell cycle control system, the remaining two Cdk1 cyclins do not need to oscillate transcriptionally.

However, past experiments showing that constant transcriptional induction suffices for cell proliferation were often performed with relatively weak promoters such as *GALL*^88^. Thus, we re- examined elements of cell cycle control for being essential for one part of the cell cycle but poisonous for another, at least, when over-expressed. Once found, we could engineer cells to be even more sensitive to both the need for the cell cycle regulator at one cell cycle stage and poisonous for the other.

For our search, we used the relatively strong blue-light-induced LOV transcription factor El222 and the El222-controlled *LIP* (**Supplementary Note 3**). This system allowed us to test rapid switches in target gene expression in glucose medium, where proliferation is relatively fast. We transformed yeast with constructs in which *LIP* controlled the transcription of cell cycle regulators we wished to test (Table 1, column 1). In these cells, we had deleted endogenous cell cycle control genes whose absence makes the corresponding inducible cell cycle gene essential. Further, cells were kept viable before our tests by inducible transcriptional constructs that could be shut off for the test (Table 1, column 2). For more details on cell cycle regulation in budding yeast generally and in our final strain, see **Supplementary Note 6**.

**Table 1:**
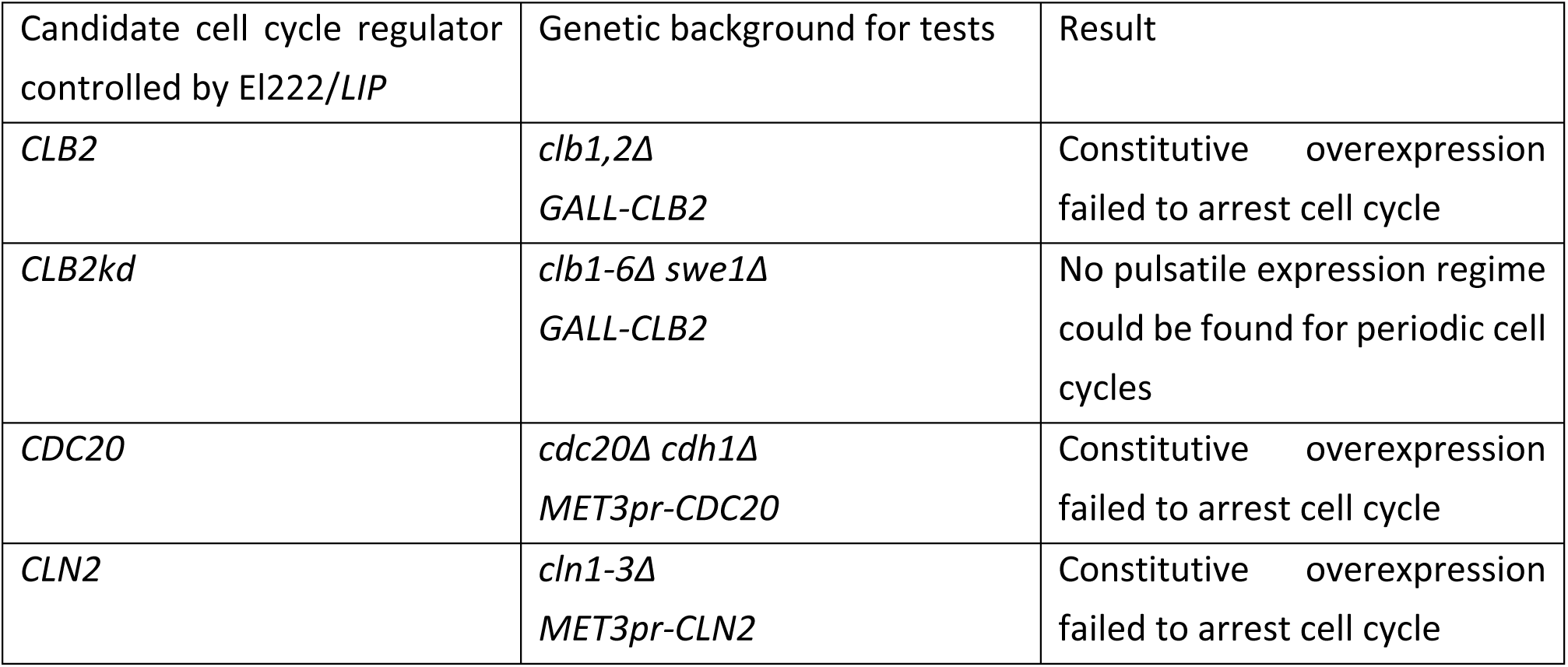
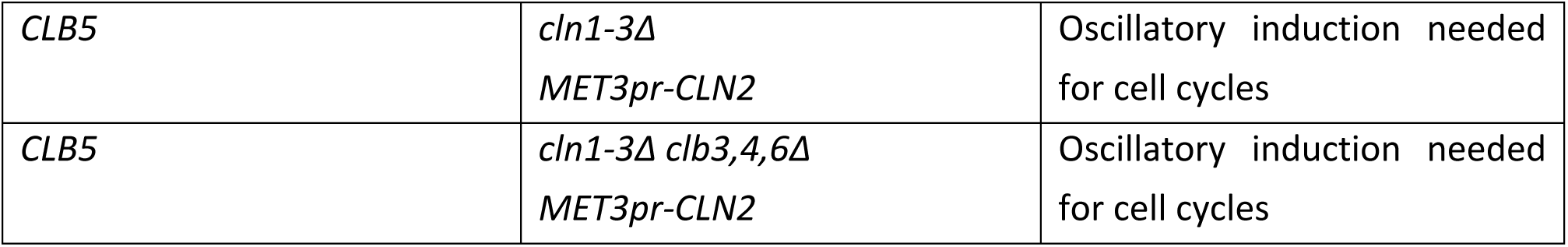
Tests to identify cell cycle regulators which are needed for one part of the cell cycle but toxic for another in specially engineered genetic backgrounds. During the tests, we turned off the nutrient-inducible promoters *MET3pr* and *GALL*.

We tested for three different properties: inviability given continuous off (0) or on (1) expression as well as proliferation under periodic expression. We started with the key components of the core cyclin-Cdk/APC negative feedback loop of the cell cycle (Table 1, Fig. 2 A). First, we tested optogenetic control of *CLB2*, the major M cyclin in budding yeast^124^, in a *clb1,2Δ* background. As expected, in darkness, the lack of *CLB1* and *CLB2* caused cells to arrest before mitosis. However, continuous expression of *LIP-CLB2* did not prevent cells from cycling in glucose medium. Since Clb2 activity has to be reduced for cell cycle exit^125^, we next tested whether Clb2kd, an undegradable variant of Clb2 was more suitable. Clb2kd continued to be needed for entry into mitosis and anaphase in a *clb1,2Δ* background. However, we could not find a regime of pulsing light under which cells performed periodic cell cycles. After a few cell cycles, Clb2kd blocked mitotic exit nearly permanently presumably because Sic1 inhibition and dilution did not reduce Clb2kd activity sufficiently. (Budding yeast cells have been shown to be very sensitive to the duration and period of joint Start cyclin and *GAL1pr-CLB2kd* pulsing over only a few cell cycles; this was interpreted as a signature of an artificial incoherent feedback loop^126^.) Next, testing the B-type cyclin inhibitor *CDC20*, whose deletion leads to metaphase arrest, we found that it was similarly not suitable since *LIP-CDC20* overexpression did not arrest cells.

**Fig. 2.**
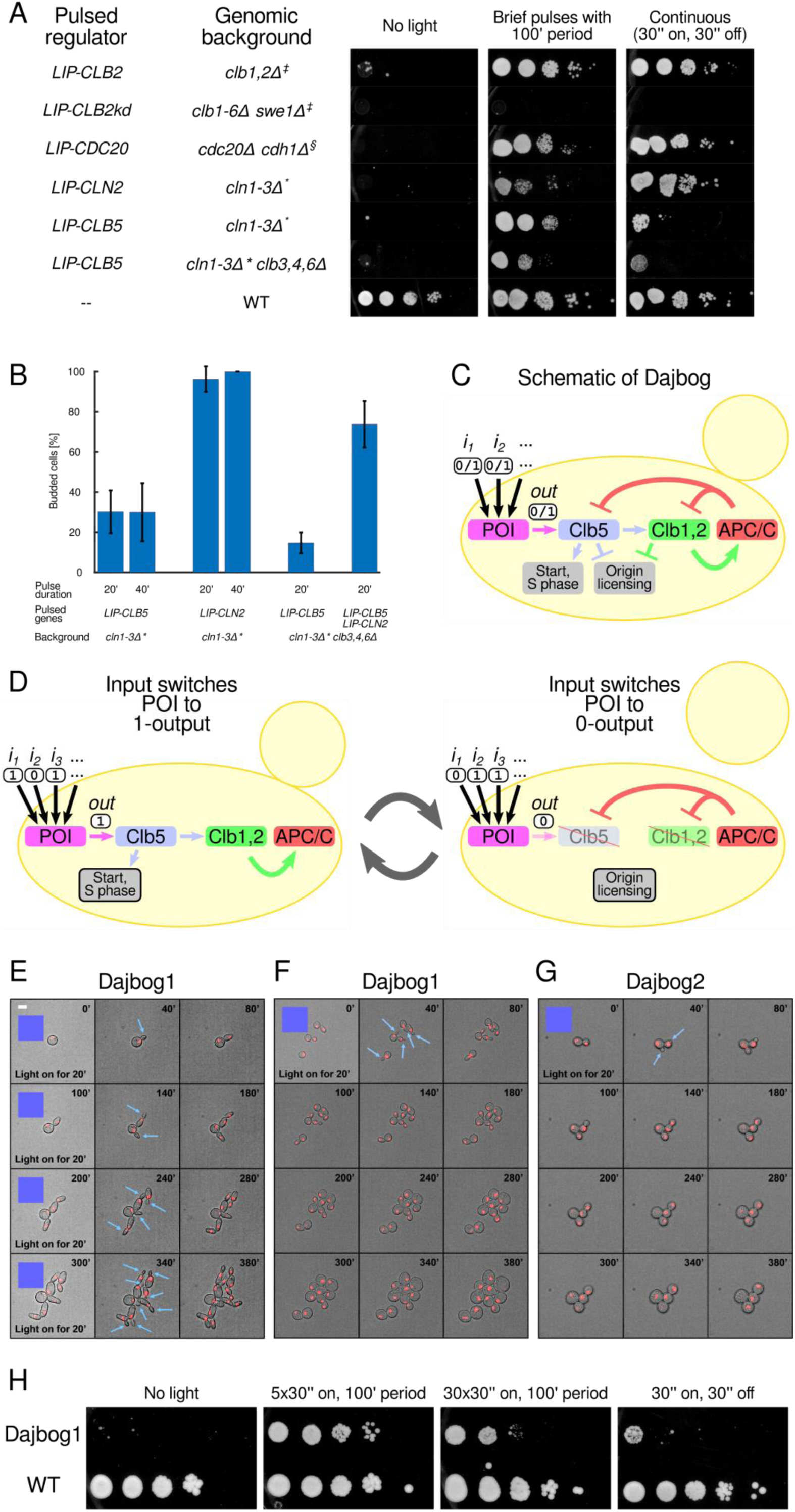
Engineering budding yeast strains for optovolution. A. Evaluation of different cell cycle regulators for three properties: essential for proliferation (cells inviable when regulator is off), sufficient for proliferation when pulsed with cell cycle period (5 light pulses of 30 sec on, 30 sec off administered every 100 min, except for *cln1-3Δ clb3,4,6Δ* when we used 30 such light pulses), and lethal when expressed continuously (30 sec on, 30 sec off). (We used a 50% duty cycle, that is, 30 sec on and 30 sec off, to reduce light toxicity to a negligible level, see wild type in panel A.) All cells expressed *EL222*, which induces *LIP*. Spot tests were performed with ten-fold dilutions. (Note that panel F shows tests for the Dajbog1 (*cln1-3Δclb3-6Δ*) strain, which has been improved with respect to these three properties.) B. Efficiency of cell cycle initiation (budding) when different light-inducible cell cycle regulators are activated, assessed by timelapse microscopy. Error bars show 5-95% confidence intervals. Number of scored cells are given in Supplementary Table 3. C. Schematic of cell cycle regulators in Dajbog strains. A subset of key cell cycle elements is shown only. Cln2 and Sic1 were omitted for clarity. D. Illustration of cell cycle states of the Dajbog strains when the POI output is on (1) or off (0). E. Timelapse recording of Dajbog1 strain, exposed to diascopic microscope light every 100 min for 20 min (Supplementary Figure 1 C). Cells carry the red fluorescent nuclear marker Htb2-mCherry. Blue arrows point to buds. Scale bar, 5 µm. Blue square, time point with optogenetic light induction. F. Timelapse recording with only one light pulse administered for 20 min. G. Timelapse recording of Dajbog2 cells (Dajbog1 with *sic1Δ* deletion) with only one 20 min light pulse. H. Spot test of Dajbog1 strain under different light pulse regimes, revealing 5 times 30 sec on, 30 sec off every 100 min to be optimal among the conditions tested. In the middle two panels, five or thirty 30 sec light pulses, respectively, are separated by 30 sec of no light (off). In the right panel, 30 sec on, 30 sec off light pulses are administered continuously. Spot tests were performed with ten-fold dilutions. To keep cells proliferating before the tests, cells carried nutrient-inducible *MET3pr-CLN2* (*), *GALL-CLB2* (‡), or *MET3pr-CDC20* (§) constructs, which were shut off during the tests by glucose or methionine in the agar plates.

The cell cycle regulators discussed thus far control mitotic transitions, where arrest and release often induce aneuploidy. We next shifted our focus to cyclins which control the G1-to-Start transition, which is less perilous for genomic stability. While pulsing of the G1/S cyclin *CLN2* has been extensively tested,^126^ constitutive over-expression of *CLN2* from El222/*LIP* in the *cln1-3Δ* background, where *CLN2* is essential, did not kill cells. Thus, we turned to the S-phase cyclin *CLB5*. We first tested *LIP-CLB5* in a *cln1-3Δ* strain since *CLB5* expression is known to override *cln1-3Δ* G1 arrest^127^, although with lower efficiency than the G1/S cyclin *CLN2*. In darkness, cells were arrested in G1. Light pulses with a 100 min period led to repeated cell cycles. Fortuitously, we found that constitutive *CLB5* expression by El222/*LIP* severely impeded proliferation and was lethal over multiple cell cycles . (In contrast, *GAL1pr-CLB5* overexpression in galactose medium, as opposed to glucose medium, where cell cycles are substantially slower, is tolerated.^128^) Thus, *CLB5* was a promising candidate for tightly coupling the cell cycle to the POI.

Next, we engineered the system for mutational robustness. To further increase the cells’ dependence on *LIP-CLB5* expression, we deleted the four chromosomal *CLB3,4,5,6* gene copies, which can be functionally redundant with light-induced *CLB5*^129,130^. No mutations are known that can bypass the requirement for Clb5 in a *cln1-3Δ clb3-6Δ* background^131^, where Clb5 triggers Start, S phase, and mitotic entry. We named this genotype Dajbog1 (including appropriate constructs for *CLB5* and *CLN2* control) after the mythological Slavic deity coupling astronomical cycles and light to life^132^.

The reliability of cell cycle initiation in response to a *LIP-CLB5* pulse, however, was low (Fig. 2 B). Only 30% of *cln1-3Δ* cells budded in response to a *LIP-CLB5* pulse. In *cln1-3Δ clb3,4,6Δ* cells, light- induced *LIP-CLB5* led to budding in even fewer (15%) G1-arrested cells. This fraction was much higher (>90%) for *cln1-3Δ* cells with *LIP-CLN2*. Increasing the duration of the light pulse did not change the fraction of *cln1-3Δ LIP-CLB5* cells that committed to the cell cycle. Guided by these results, we sought to increase the reliability of the response to light pulses by adding a *LIP-CLN2* construct. This does not undermine the dependence on *LIP-CLB5*, which continues to be essential for S phase and triggering entry into mitosis in the *cln1-3Δ clb3-6Δ* background. Simultaneous expression of *LIP-CLN2* and *LIP-CLB5* in the *cln1-3Δ clb3,4,6Δ* background led to a more reliable >70% of cells budding in response to a 20 min pulse of light. Altogether, the Dajbog background (Fig. 2 C, D) allowed reliable control by external light pulses, with cell cycle Start only initiated in response to a light pulse (Fig. 2 E-H).

After deleting *CLB3-6* to make the Dajbog1 strain more strongly depend on *LIP-CLB5* induction, we could next also make the strain less tolerant to continuous *LIP-CLB5* expression. This would further increase the need for Clb5 to oscillate on and off for proliferation. We accomplished this by deleting *SIC1*, a B-type cyclin inhibitor. *SIC1* is part of a circuit that can oscillate when cyclin levels are constant and can rescue such cell cycles^123^. When *SIC1* is deleted, the *cln1-3Δ* arrest becomes leaky^133^. However, we found that this depends on at least one of the *CLB3-6* genes being present. *cln1-3Δ clb3-6Δ sic1Δ* cells continued to be tightly blocked in darkness (Fig. 2 G). We named this strain background Dajbog2^132^.

To enable routine propagation of these cells prior to directed evolution campaigns, we created a ‘training-wheel plasmid’. This plasmid allowed cells to grow without exogenously induced Clb5 oscillations, and the directed evolution campaign begins with counter-selection against the plasmid. This increases the method’s tolerance for POIs that initially cannot control *CLB5* sufficiently well. We used a *URA3*-marked CEN/ARS plasmid, which can be selected against with 5-FOA, containing a *CLB5* copy and *MET3pr-CLN2*. Finally, to increase the mutation rate for directed evolution, we introduced the *pol3-L523D*^134–136^ mutation that is reported to increase the global mutation rate about one-hundred fold.

In summary, we achieved tight coupling of the cell cycle machinery to externally applied periodic on-off light signals using Clb5 (aided by Cln2) in a *cln1-3Δ clb3-6Δ SIC1/sic1Δ* background (Fig. 2 C- H). In principle, any POI controlling *CLB5* (and *CLN2* for higher efficiency) can be inserted for directed evolution.

### Overview of directed evolution experiments

We performed six directed evolution campaigns with the Dajbog strains, in which we inserted different POI genes to control cell-cycle progression (four campaigns with El222, one with PhyB-Pif3, and one with PEST-rtTA). All campaigns followed the same procedure (Fig. 3): The Dajbog strains were transformed with genetic constructs controlling both *CLN2* and *CLB5* simultaneously in *trans*, and were selected for loss of the training wheel plasmid under a permissive light pulse regime (preparatory strain manipulation not shown in illustration). Next, the cells were spread on agar growth medium plates (Fig. 3 A, G) and allowed to proliferate until the plate was covered by a lawn of cells (Fig. 3 B, H). The plate was then briefly exposed to a 30 W, 254 nm UV lamp. Immediately afterwards, cells were either spread in a thin layer on agar plates (Fig. 3 C, I) or suspended at low optical density (OD_660_ = 0.1) in liquid growth medium (Fig. 3 E, K, L), depending on the campaign. (Since we wished to explore the diversity of mutational solutions, we opted for the agar plate format in five out of our six campaigns.) Then, the desired selection pressure was imposed by changing to a restrictive light pulse pattern under which the initial cells could not proliferate. We chose both initial and final light pulse patterns without fine-tuning: Starting with the light pulse regime where cells clearly proliferated well during strain construction, we chose a very different regime in which the desired POI property was necessary for proliferation and the vast majority of Dajbog cells were obviously arrested.

**Fig. 3.**
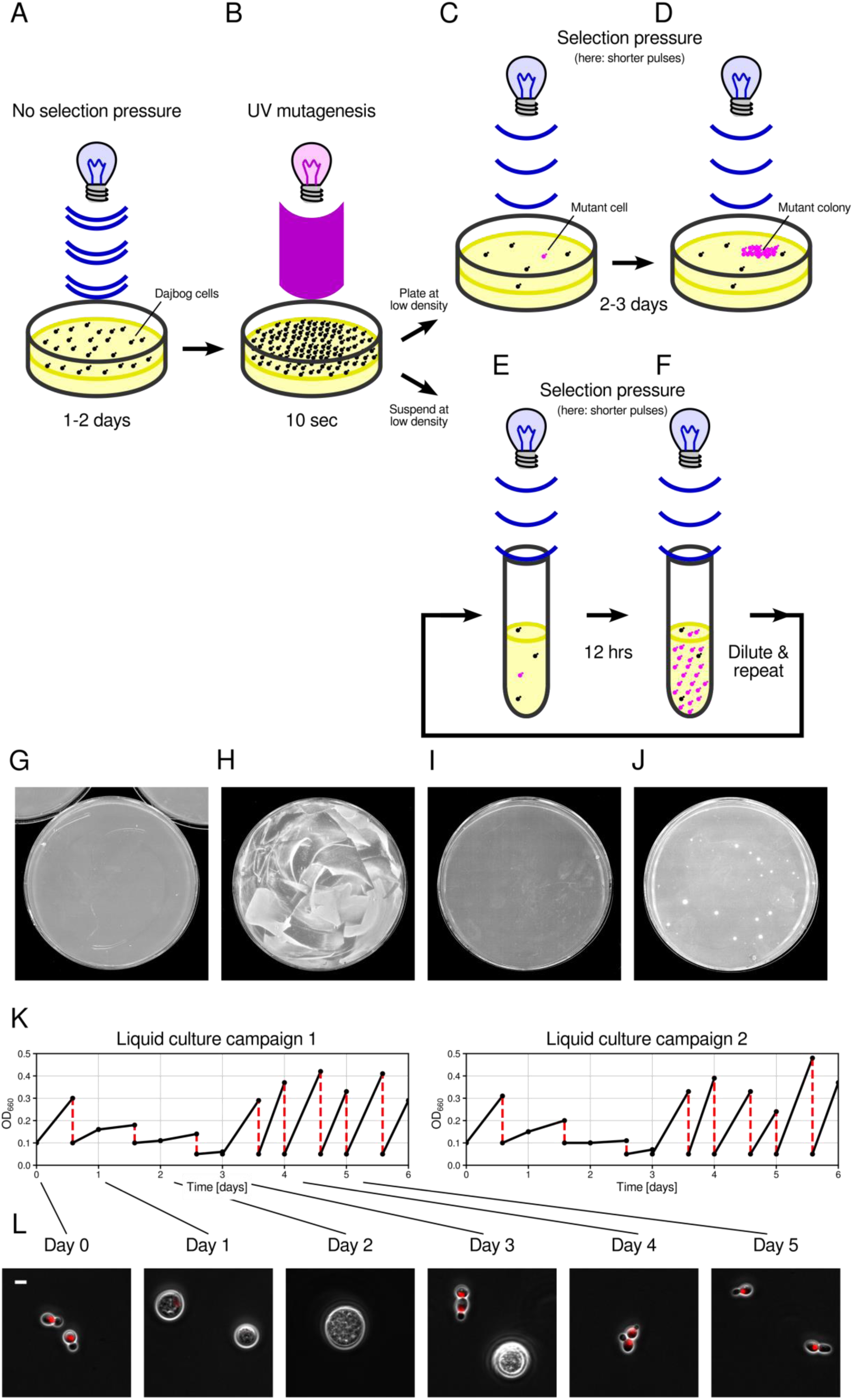
Procedure of optovolution experiments. A-F. Illustration of optovolution experiments on agar plates or in liquid cell culture. G-J. Representative pictures of plates of Dajbog cells: G, at the beginning of the experiment; H, after 2 days of proliferation without selection pressure; I, after spreading on fresh agar plates; and J, after three days subjected to selection pressure. K. OD_660_ measurements during two independent liquid culture experiments. Black circles indicate measurement points. Lines solely serve to guide through the sequence of measurements. Dilutions are marked by dashed red lines. L. Representative images of cells at different days of optovolution in liquid culture. Red, Htb2-mCherry. Scale bar, 5 µm.

On agar plates, mutant colonies emerged after 2 days and were then analyzed (Fig. 3 D, J); the death of all other cells on the plate was evidence for the stringency of our selection pressure. Because optovolution does not require intervention and media changes, cells can remain on the same agar plate. While we stopped evolution here to analyze the mutations by Sanger or whole-genome sequencing, colonies could be propagated by standard replica plating. For the liquid culture experiments, the cell culture was diluted twice per day into fresh medium to maintain log phase (Fig. 3 E, F), unless the OD did not change substantially compared to the previous timepoint. After the switch to a restrictive light pulse regime, the OD rose for 1.5 days, presumably due to cell growth without proliferation (Fig. 3 K, L), and then plateaued until about day 3. The lack of change of OD indicated that the restrictive light pulsing conditions and resulting long arrests had killed the majority of cells. After this period, we observed a more typical OD rate of increase, when adaptive mutants took over the cell culture. After passaging cells for another 3 days, we analyzed the bulk cell culture for mutations in the POI gene. The frequency of mutations from standing variation, error-prone DNA replication, and additional exposure to 10 sec of UV sufficed to find desired mutations in every campaign (**Supplementary Table 4**).

### Validation of mutants

We functionally validated the mutations *in vivo*: We cloned all mutations that emerged and introduced them in wild-type budding yeast cells where the POI drove transcription of the bright, yeast-optimized *yEVenus* or *ymScarletI* fluorescent protein genes (**Methods**). We followed the characterization procedure of ref.^88^, using degron-destabilized fluorescent proteins for dynamic time course measurements, fluorescence microscopy for relatively high precision, and unbiased neural network-based image processing^137^. All fluorescence levels are expressed in maxGAL1 units^88^, the maximal transcriptional output from the well-known *GAL1* promoter, which provides a straightforward, intuitive means to gauge the strength of transcriptional induction across different systems and publications.

Crucially, potential mutations elsewhere in the genome of the Dajbog cells, which could have emerged in the directed evolution campaign, could not affect our characterizations of the mutations. We thus established the causal effects of the evolved mutations independently of their ability to support proliferation under the restrictive light pulsing regime in the Dajbog strains.

### Directed evolution of a LOV-transcription factor

We first applied the method to optogenetic systems, i.e., the POI itself was light sensitive. Optogenetic systems relay external on (1) or off (0) light states to on (1) or off (0) cellular activity, i.e., they have two input-output states. Optogenetic systems have been challenging to evolve, requiring repeated, sequential campaigns for the on (1) and off (0) output states since mutations improving one output state may destroy the other.^138^ Given that El222 is itself widely used and that LOV domains are of broad interest, we chose to evolve El222 in three different directions, toward i) greater light sensitivity, ii) less leakiness, or iii) sensitivity to different colors for multiplexing and reduced phototoxicity.

We used the Dajbog strains with one copy of *EL222* directly controlling the transcription of both *CLN2* and *CLB5* in *trans* in the same configuration as for testing and developing our method (Fig. 4 A). We used the experimental procedure illustrated in Fig. 3. Cells initially proliferated under a permissive light pulse regime (5 times 25 sec on, 35 sec off every 100 min). Then, we changed the light pulse regime to apply one of three selection pressures, in which cells with wild-type El222 could not proliferate (Fig. 4 B). (See Supplementary Figure 1 for the excitation and imaging light spectra.)

**Fig. 4.**
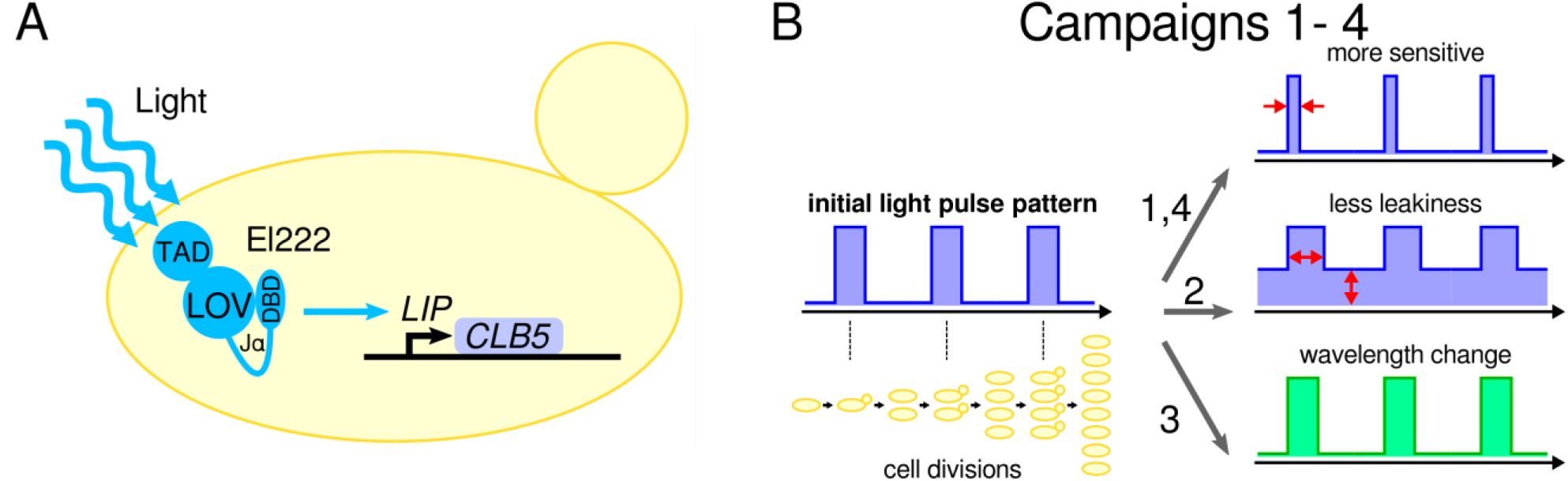
Directed evolution campaigns 1-4 with LOV-transcription factor El222 as POI. A. Schematic of the wiring diagram for El222 evolution. *CLN2*, which iscontrolled by the same copy of the *EL222* gene, and other cell cycle regulators omitted for clarity. TAD, transcriptional activation domain; LOV, LOV domain; Jα, Jα helix; DBD, DNA binding domain. B. Schematic of four directed evolution campaigns. The initial light pulse pattern was switched to very brief pulses (campaigns 1 and 4), long pulses with additional ambient light (campaign 2), or green and longer-wavelength light pulses (campaign 3). Campaign 4 was a long-term campaign in liquid culture. Under each final light pulse regime, wild-type Dajbog cells did not proliferate (see representative spot tests for each campaign, Fig. 5 A, E, Fig. 6 A).

In the first directed evolution campaign, we sought to enhance the 1-output (on) state of El222. This would allow higher output under low light conditions. To evolve more light-sensitive mutants, we exposed cells to very brief light pulses (1 sec on every 100 min, Fig. 4 B), which was insufficient for viability of the wild-type Dajbog cells (Fig. 5 A). In the colonies that grew under these conditions (Fig. 5 A), we found 11 different El222 mutations (Fig. 5 B). (Interestingly, the light-sensitizing mutations did allow extremely slow growth (about 1%) to occur in darkness (Fig. 5 A).) Remarkably, using Dajbog2 (with *sic1Δ*) we observed 96% (n = 247) of colonies had a mutation in *EL222* while Dajbog1 (with *SIC1*) permitted mutations elsewhere in the genome to rescue cells (10% in *EL222*, n = 816). Evaluating the new El222 variants cloned into our reporter strain, we found that they showed improved light sensitivity, with levels of transcription under low-intensity light up to 16 times higher than with wild-type *EL222* (Fig. 5 B). Several variants were comparable or more light sensitive than the most sensitive El222 mutant to date, strongLOV (E84D), which we had engineered previously^88^ and rediscovered here.

**Fig. 5.**
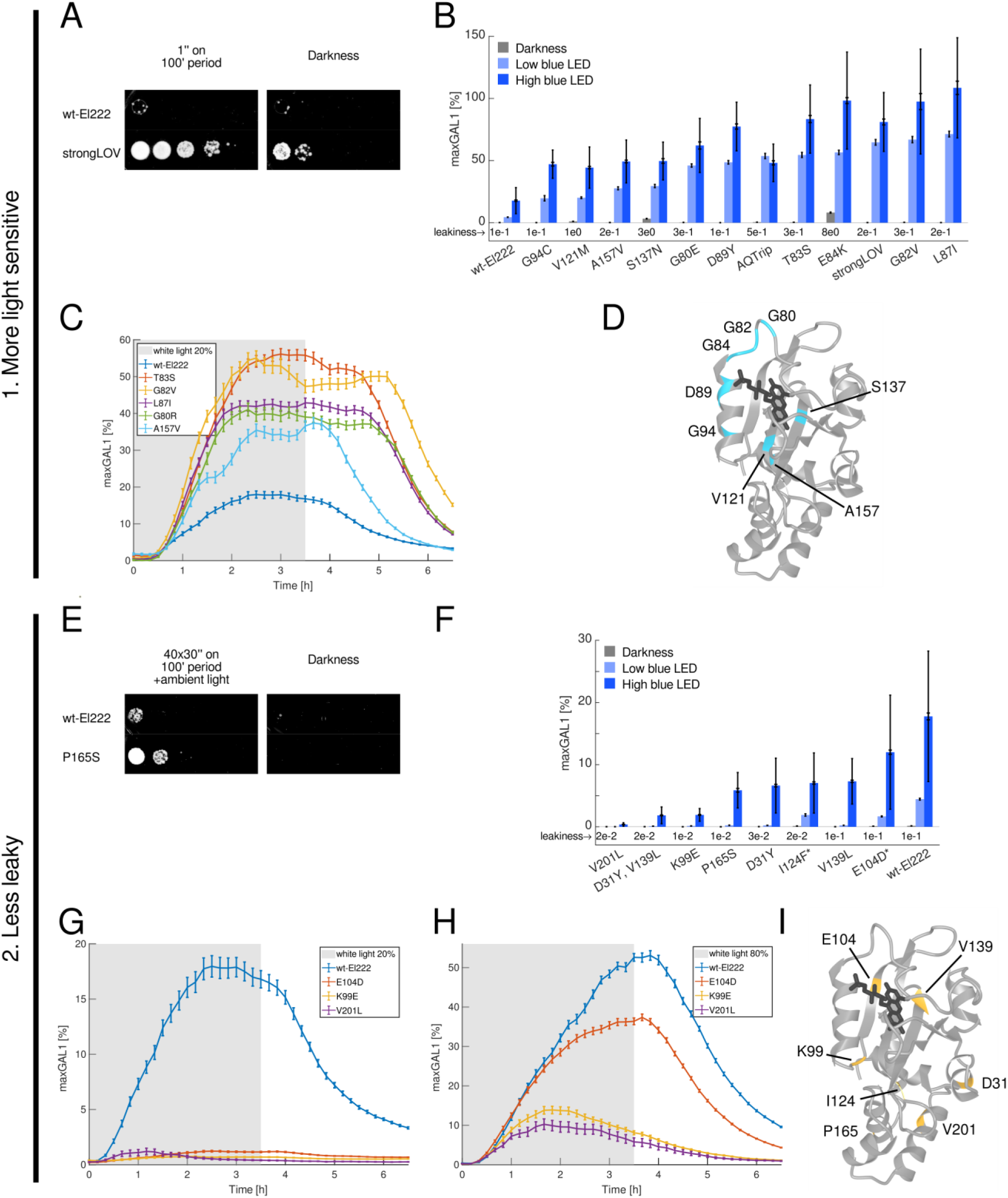
Results of directed evolution campaigns of El222 for higher sensitivity and lower leakiness. A, E. Spot tests for the wild-type Dajbog strain (ancestor) and typical evolved mutants. Spot tests were performed with ten-fold dilutions. B, F. *In vivo* steady-state expression levels of the transcriptional fluorescent reporter *LIP-yEVenus* under different light exposure conditions, as indicated, driven by mutant *EL222*, which was cloned from the Dajbog strains and introduced into the reporter strain. The inner error bars represent the standard error of the mean (SEM); the outer error bars represent the standard deviation (STD). The mean leakiness values in the dark are explicitly stated for each mutant. In panel B, we included AQTrip^140^ for reference. In panel F, * refers to mutants obtained with variations of the low-leakiness campaigns (main text). C, G, H. Time courses of El222 mutants, driving a destabilized fluorescent transcriptional reporter with the PEST degron. The optogenetic systems were induced from 0 h to 3.5 h (grey region) by diascopic light (Supplementary Figure 1)^88^. Error bars, SEM. G and H differ by the intensity of the induction light, as indicated. D, I. Locations of a subset of the mutations (blue or yellow) in the dark-state 3D structure of El222 (PDB ID: 3P7N)^139^. All references to mutant residue numbers in El222 are given with respect to bacterial El222 to match the numbering used previously^88,139,140^. B, C, F, G, H. Numbers of analyzed cells are given in Supplementary Tables 5-7. All values are in %maxGAL1 units^88^. All measured characteristics of the new El222 mutants have been added to the Inducible Transcriptional Systems Database (https://promoter-benchmark.epfl.ch/)^88^.

To characterize the dynamic properties of the evolved El222 variants, we monitored their activity by fluorescence timelapse microscopy where we induced them with diascopic white light for 3.5 h (Fig. 5 C, Supplementary Figure 1 C). All tested variants showed increased activity compared to wild-type El222 under a constant light source of 8.4 mW/cm^2^. For most variants, we observed a trade-off between strength and turn-off dynamics, similar to strongLOV^88^. However, remarkably, the highly light-sensitive variant A157V, containing a mutation in the Jα helix (Supplementary Figure 2), showed turn-off dynamics comparable to wild-type El222. (All references to mutant residue numbers in El222 are conventionally given with respect to bacterial El222 to match the numbering used in previous work^88,139,140^). This mutant is further interesting because it is the only one outside the core of the LOV domain of El222 (Fig. 5 D, Supplementary Figure 2). Within the core of the LOV domain, the Fα helix, which is adjacent to the chromophore, was particularly enriched for light-sensitizing mutations.

In the second campaign, we sought to reduce the leakiness of El222, that is, suppress activity in the 0-output (off) state. This is particularly important for applications where baseline expression needs to be minimized, e.g., when controlling toxic genes. We exposed cells to very long duration pulses of light (40 times 30 sec on, 30 sec off every 100 min, Fig. 4 B) in addition to continuous, dim background light (Fig. 5 E-H). This condition was lethal to Dajbog strains with wild-type El222 (Fig. 5 E). In a preliminary run, we found many mutations in *LIP-CLB5*, which presumably maintained Clb5 function but with reduced activity that blunted the selection pressure. Thus, we introduced additional copies of *LIP-CLB5* to block this evolutionary avenue (**Discussion**). Among the mutant colonies that emerged in this campaign, the frequency of mutations in the *EL222* ORF was 36% (n = 44).

We identified five new mutations whose leakiness was nearly indistinguishable from baseline transcription from *LIP* in the absence of El222 (Fig. 5 F, Supplementary Figure 3). As expected, they were also overall less strongly induced by light. One of the colonies that emerged had fixed two mutations in the LOV domain (D31Y, V139L). To understand their individual contributions to the phenotype, we tested them separately. The mutations contributed independently and about equally under high blue LED light. We also measured the on-off dynamics of low-leakiness mutants by fluorescence microscopy. Using continuous light of 8.4 mW/cm^2^ intensity as in the case for light-sensitized mutants led to little observable activity in the low-leakiness mutants (Fig. 5 G). Under much stronger light (97 mW/cm^2^), the mutants responded noticeably (Fig. 5 H). The low-leakiness mutations were both in the LOV domain as well as the DNA binding domain (Fig. 5 I). To test whether the combination of long induction light pulses and background light had been necessary to discover these mutants, we performed additional campaigns with only the long duration pulses or only background light. We discovered two additional mutations (marked with * in Fig. 5 F) in these campaigns. Their properties were comparable to the mutations found previously.

In our third campaign, we sought to modify the nature of the 1-output state with respect to color responsiveness. After proliferating Dajbog cells under blue light pulses, we evolved them under green, orange, and red light combined (40 times 30 sec on, 30 sec off every 100 min). (Figs. 4 B, 6, Supplementary Figure 1 for details regarding light sources). Among the mutant colonies that emerged, we discovered mutations in *EL222* that allowed cells to proliferate under these LED light pulses (Fig. 6 A). Optovolution was highly efficient: 96% (n=247) of mutations were in the *EL222* ORF when Dajbog2 (with *sic1Δ*) was used and 10% (n = 816) with Dajbog1 (with *SIC1*). As before, we validated the mutations functionally by cloning the evolved *EL222* genes into a reporter strain. To monitor turn-on and turn-off dynamics longitudinally at the single-cell level, we characterized the mutant El222 proteins in the reporter strains by timelapse microscopy (Fig. 6 B-H, J-M). As transcriptional reporter, we chose *LIP-mScarletI-PEST*, which was excited by 562±20 nm light for 100 ms every 10 min for imaging, minimizing unwanted potential optogenetics activation. We covered the relevant part of the light spectrum for inducing the optogenetic system using 10 nm bandpass filters and the microscope’s white light source (Supplementary Figure 1). In the following, we indicate the optogenetic induction light by the central wavelength of the filter and the width of ±5 nm.

**Fig. 6.**
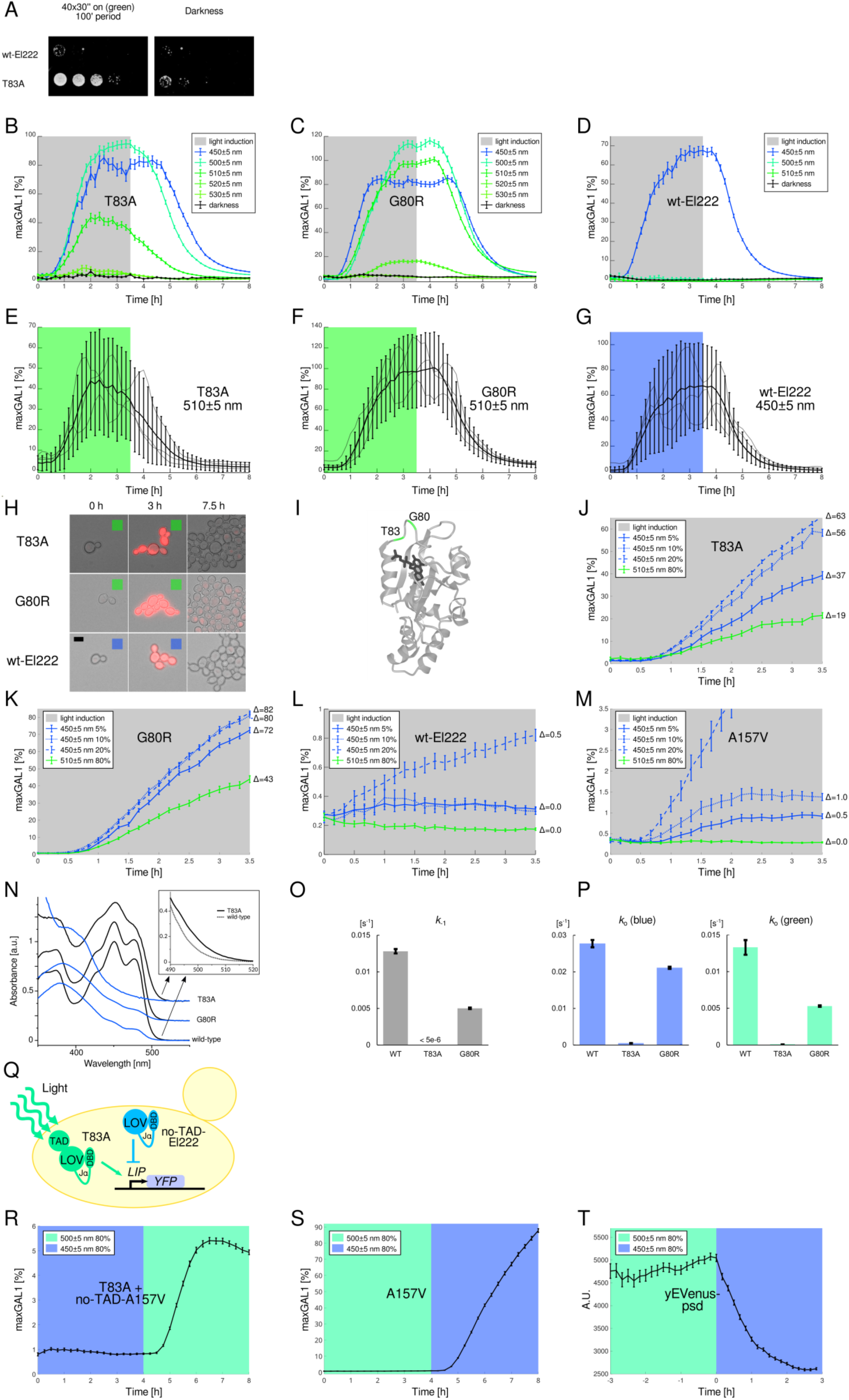
Results of optovolution campaigns for spectrally modified El222. A. Spot tests for the wild-type Dajbog strain (ancestor) and the evolved T83A mutation. Spot tests were performed with ten-fold dilutions. B, C. Time courses of the T83A and G80R El222 mutants cloned from evolved Dajbog strains and introduced into the reporter strain driving *LIP-ymScarletI-PEST*. The optogenetic systems were induced from 0 h to 3.5 h (grey region) by diascopic white light (Supplementary Figure 1) filtered through different 10 nm bandpass filters. Plot colors correspond to the center wavelength of the filters. D. Time course for wild-type El222. Error bars in B-D indicate the standard error of the mean (SEM). E-G. Mean, standard deviation, and three representative single-cell time courses for the T83A and G80R mutants induced by green light and wild-type El222 induced by blue light. The mean time course with SEM error bars is also shown in panels B-D. H. Snapshots of different reporter cell strains (different rows) at three time points in the timelapse recordings (different columns). Colored squares indicate exposure to optogenetic induction light at the respective time points. Scale bar, 5 µm. I. Locations of the mutations in the dark-state 3D structure of El222 (PDB ID: 3P7N)^139^. J-M. Sensitivity of the T83A, G80R, A157V mutants, and wild-type El222 to green light and different intensities of blue light measured using a *LIP-ymScarletI* reporter. Δ values on the right side of the plots indicate the difference between final values at 3.5 h and initial values at 0 h in %maxGAL1 units. Error bars, SEM. N. Normalized dark- and light-state absorption spectra of purified El222 variants, offset along the vertical axis for clarity. Inset presents a superposition of the T83A and wild-type spectra at the long-wavelength tail. O. Dark reversion rate constants of the purified El222 variants. The value for the T83A mutant represents an upper bound. P. Observed photoactivation rate constants *k*_o_ of purified El222 variants under blue and green light. Q. Schematic for green-only transcriptional activation system using El222(T83A) and a no-TAD-El222. R. Time course of T83A-driven *LIP-ymScarletI* expression under simultaneous expression of *GAL1pr-no-TAD-EL222(A157V)* in raffinose + galactose medium and a switch from blue to green light. S. Time course of the A157V mutant under the same conditions as in panel R but without a competitor. T. Time course of yEVenus tagged with *At*LOV2-based photo-sensitive degron (psd)^120^ and expressed under the control of the *PGK1* promoter. Error bars in panels O, P, R-T represent SEM. All references to mutant residue numbers in El222 are given with respect to bacterial El222 to match the numbering used previously^88,139,140^. Numbers of analyzed cells are given in Supplementary Table 6. All values are in %maxGAL1 units^88^. All measured characteristics of the new El222 mutants have been added to the Inducible Transcriptional Systems Database (https://promoter-benchmark.epfl.ch/)^88^.

We discovered two mutants, G80R and T83A, which responded strongly to green light, only markedly reducing in activity at 520±5 nm and above (Fig. 6 B, C). In contrast, wild-type El222 showed no responsiveness at 500±5 nm or longer wavelengths, even at the highest light intensity we could achieve (80% = 0.26 mW/cm^2^) (Fig. 6 D). Variability and noise were similar in T83A and G80R in green light as wild-type El222 in blue (Fig. 6 E-H). Both G80 and T83 residues are in close proximity to the chromophore (Fig. 6 I).

To investigate whether responsiveness to green light was merely a consequence of overall higher sensitivity to light, we sought to measure the ratio of the response to blue and green light (Fig. 6 J-M). We exposed our reporter cells to 510±5 nm light and to different intensities of 450±5 nm light. To maximize measurement sensitivity, we used the *LIP-mScarletI* construct, i.e., without the PEST degron. (We only show induction in Fig. 6 J-M since the long lifetime of the reporter is not informative of the turn-off dynamics of the optogenetic system, which is already shown in panels B-H.) Without the PEST degron, the baseline activity of each mutant was precisely detectable and is represented by the first timepoint in the time courses. For green 510 nm light induction, we applied the highest intensity we could achieve with our light source (0.26 mW/cm^2^). Nevertheless, this green light elicited no detectable response with wild-type El222 (the fluctuations in Fig. 6 L are small, on the scale of ∼0.1% maxGAL1). However, this green light led to robust expression of the fluorescent reporter with T83A and G80R (Fig. 6 J, K). On the other hand, the intensities of blue light that were sufficient to strongly activate and saturate T83A and G80R (10%-20%, Fig. 6 J, K) were just high enough to activate wild-type El222 (only 20% led to detectable activation, Fig. 6 L). The relative insensitivity of El222 to light overall prevented us from making a definitive comparison of relative green-to-blue responsiveness. Thus, we utilized the A157V mutation, which led to greater light sensitivity than wild-type El222 (Fig. 5 B, C) but whose location far from the chromophore (Supplementary Figure 2) was unlikely to differentially affect color sensitivity. Indeed, A157V responded with all blue light intensities tested (5%, 10%, and 20%) but, again, showed no response to green light (Fig. 6 M). Thus, our mutants show differential sensitivity to 510±5 nm light *in vivo* compared to wild-type El222 or A157V, neither of which showed a detectable response to clearly visible, strong green light.

To investigate the biophysical characteristics of the *in vivo* green-responsive mutants, we produced wild-type El222, and the T83A and G80R variants by heterologous expression in *E. coli* and purification. UV/Vis absorption spectroscopy revealed that in their dark-adapted states all three El222 receptors bind flavin-nucleotide chromophores and exhibit characteristic LOV absorbance spectra with peaks around 450 nm and two shoulders at about 430 and 475 nm (Fig. 6 N, Supplementary Figure 5 A). Compared to the other proteins, the spectrum of T83A had a less pronounced fine structure which could be indicative of looser flavin binding. Crucially, the T83A (but not G80R) spectrum revealed a broader tail at long wavelengths that incurred higher relative absorbance in the 500-520 nm range (Fig. 6 N inset).

Once exposed to strong blue light, all El222 proteins adopted their light-adapted state characterized by a shallow absorbance peak around 390 nm. Returned to darkness, wild-type El222 thermally recovered its dark-adapted state in a single-exponential manner with a rate constant *k*_-1_ of 1.28 × 10^-2^ s^-1^ at 25 °C (Fig. 6 O, Supplementary Figure 5 B), corresponding to a time constant *τ* of around 75 s. Owing to a different buffer composition and pH, the kinetics are slower than originally reported^139^. The G80R variant recovered around 2.5-fold more slowly to its dark-adapted state with a rate constant of 5.0 × 10^-3^ s^-1^. By contrast, the dark recovery reaction of the T83A variant was exceedingly slow and incomplete even after extended times (Supplementary Figure 5 B). Based on the observation that after around 16 h only 20% of the initial absorbance at 450 nm was regained, we estimate an upper limit of the recovery rate constant *k_-1_* of around 5 × 10^-6^ s^-1^.

We next addressed to which extent El222 is photoactivated by blue versus green light (Fig. 6 P, Supplementary Figure 5 C, D). To this end, we tracked the gradual decrease of the dark-adapted state upon exposure to light passed through 450-nm and 510-nm filters (10 nm bandwidth in both cases) at powers of 0.75 mW cm^-2^ and 1.5 mW cm^-2^, respectively. The observable rate constant *k*_o_ obtained by evaluating the photoactivation time courses according to single-exponential functions is a sum of the light-driven forward rate constant *k*_1_ and the reverse dark-recovery rate constant *k*_-1_. For instance, under 450-nm light the dark-adapted state of wild-type El222 decayed with an observable rate constant *k*_o_ = 2.77 × 10^-2^ s^-1^ from which a forward rate constant *k*_1_ of 1.5 × 10^-2^ s^-1^ results. Consistent with the similar magnitude of *k*_1_ and *k*_-1_, for these experimental settings a photostationary state resulted with around 51% population of the light-adapted state. Green light also drove the transition to the light-adapted state but with much slower observable kinetics of 1.3 × 10^-2^ s^-1^ and only 5% amplitude, from which a value of around 5 × 10^-4^ s^-1^ results for *k*_1_ of green-light-driven photoactivation. Considering the individual light powers used in the experiments, green light thus proved around 60 times less efficient at activating the wild-type LOV receptor compared to blue light. LOV activation by green light is thus possible, in principle, although too inefficient to lead to sufficient photoactivation as observed *in vivo*. For G80R, blue and green light promoted conversion to the light-adapted state with observable kinetics of 2.1 × 10^-2^ s^-1^ and 5.3 × 10^-3^ s^-1^ and amplitudes of 66% and 14%, respectively. From these data, we determine photoactivation kinetics similar to those of wild-type El222 with *k*_1_ amounting to 1.6 × 10^-2^ s^-1^ and 3 × 10^-4^ s^-1^ under blue and green light, respectively. While the photoactivation therefore did not substantially differ between El222 wild-type and G80R, the much slower dark recovery in G80R gives rise to a 2- to 3-fold higher population of the light-adapted state under constant illumination at the conditions investigated, which at least qualitatively accounts for the higher effective light sensitivity seen above for the *in vivo* data^141^. In the case of the T83A variant, the activation by both blue and green light was remarkably slower than for the other two El222 proteins with observable rate constants *k*_o_ of 5.6 × 10^-4^ s^-1^ and 9.1 × 10^-5^ s^-1^. Given that the dark recovery in T83A proceeds with a rate constant of 5 × 10^-6^ s^-1^ or less, these kinetics are dominated by the forward rate constant *k*_1_. That is, *k*_1_ is essentially equal to the observable rate constants *k*_o_, and T83A is therefore only 12 times less efficiently activated by green than blue light, explaining the observed *in vivo* differential green responsiveness and consistent with the long-wavelength tail of the dark-adapted T83A absorbance spectrum (Fig. 6N inset). We note that modulation of the dark-recovery kinetics is well developed in LOV receptors^89^ but only scarce information exists for the variation of the photoactivation kinetics^142^.

We wished to showcase the utility of the *in vivo* green-responsive mutants for orthogonal color-multiplexing with LOV domains only. The key obstacle to achieving independent control was that the green-responsive mutants remained responsive to blue light. We thus pursued a strategy of combining the T83A mutant with a spectrally wild-type El222 which lacked the transcriptional activation domain (TAD) and should competitively inhibit binding of El222(T83A) to DNA in blue light only (Fig. 6 E). In green light, on the other hand, El222(T83A) could continue to activate transcription. The first inhibitor that we constructed, no-TAD-El222, could not suppress transcription in blue light, even when overexpressed from the *GAL1* promoter (Supplementary Figure 4 A, B). However, using the light-sensitized A157V mutant, we found no-TAD-El222(A157V) to fully suppress transcription in blue 450±5 nm light but permit transcription in 500±5 nm or 510±5 nm light (Fig. 6 R, Supplementary Figure 4 C). Under the same conditions, wild-type El222 and A157V only activate under blue 450±5 nm light (Fig. 6 S, Supplementary Figure 4 B). These results establish the first demonstration of two-color, orthogonal transcriptional control using only LOV domains, providing a modular strategy for optogenetic multiplexing.

An important question about green sensitivity, both for multiplexing and for mechanistic understanding, is whether other LOV proteins are generically green responsive. The long-wavelength tail (500-520 nm) of the wild-type El222 absorption spectrum is clearly below that of the T83A variant – but could still be detected (Fig. 6 N inset). This raises the possibility that El222 is ‘close’ to being green responsive and differences in individual amino acids could make other existing LOV proteins green responsive. On the other hand, if other LOV domains are not generally green responsive, our evolved mutations, which enable the green-only inducible transcriptional circuit (Fig. 6 Q-S), will be particularly valuable for multiplexing, since other LOV domains would be only activated by blue light. Thus, we tested *in vivo* green responsiveness in the widely used wild-type *At*LOV2 domain. We tagged a constitutively expressed yEVenus gene with psd^120^, an *At*LOV2-based degron, and tested its response under blue 450±5 nm and green 500±5 nm light, as used previously (Fig. 6 Q-S). We observed degron activity only in blue light, not in green light (Fig. 6 T). We also verified the absence of a response at 510±5 nm (Supplementary Figure 4 D). The absence of green responsiveness in two widely used LOV systems, wild-type El222 and *At*LOV2 suggests that the T83A and G80R mutations unlock in El222 a highly useful and not commonly shared property.

In total, we discovered 19 new El222 mutations, mostly in the core of the LOV domain, one in the Jα helix, and two in the DNA binding domain (Figs. 5 D, 5 I, 6 I). The analysis of the mutation spectrum in the first three campaigns showed that we observed transitions (A-G, C-T) and transversions (purine-pyrimidine) with about equal probability (**Supplementary Table 4**).

LOV domain proteins such as El222 are ubiquitous across the tree of life and control a variety of cellular processes. Most approaches to altering LOV-optogenetic tools have involved introducing mutations from other known natural LOV domains, which were faster or slower cycling^87,89,140^. To gauge how realistic it would have been to discover our evolved mutants rationally, we quantified the novelty of our mutants by searching a catalog of 7000 known and putative LOV-domains^143^ (Supplementary Figure 6 A) as well as by aligning their sequences with the well-studied LOV-domain proteins FKF1 from *Arabidopsis thaliana*, VVD from *Neurospora crassa*, YtvA from *Bacillus subtilis*, *Ds*LOV from *Dinoroseobacter shibae*, and the LOV2 domain of phot1 from *Avena sativa* (Supplementary Figure 6 B). We discovered that the majority of the mutations (15/19) were rare (≤10%) in putative LOV domains, and 6 are present in LOV domains whose kinetics had previously been characterized. V121M found in VVD is part of the rationally-designed El222 variant AQTrip^140^. Similarly, another mutation (E84D) was used to construct the more light-sensitive strongLOV mutant protein by us recently^88^. Interestingly, the other two mutations, K99E and V139L, would have been expected to lead to more stable light-adapted states based on previously measured off-kinetics^87^. We found them, instead, to lead to lower leakiness and overall weaker responses. None of the LOV proteins with characterized kinetics contained the T83A or the G80R mutation, which lead to green sensitivity in El222. These results further underscore the power of directed evolution to uncover functional mutations with specific desired properties.

### Long-term optovolution of light-sensitized LOV domains

The directed evolution campaigns described thus far were unbiased by design: Any El222 mutant which emerged and which could switch on and off repeatedly for ≈2 days under the respective selection pressure would generate a colony and be detected by us. In our fourth campaign, we wished to test whether optovolution could be used for directed evolution over longer periods of time in liquid culture, where mutations could compound and compete against one another, in principle. The fittest mutant would take over the cell culture.

We implemented the experimental approach presented in Fig. 3 A, B, E, F (additional details in **Methods**). We again sought to boost the 1-output activity of El222, this time over one week. We evolved the Dajbog2 strain under 5 sec of light, administered every 100 min. The OD time course (Fig. 3 K) showed an initial increase likely from arrested cells growing in size (Fig. 3 L). Between days 1.5 to 3, the OD did not change noticeably, reflecting the inability of the majority of cells to grow or proliferate. Subsequently, the growth in OD returned to normal, indicating that mutant Dajbog2 cells had taken over the cell culture. At the end of the week, we stopped the experiment and sequenced *EL222*. We observed three mutations that won in four different long-term experiments, E84D, G82V (twice), and V121M, which we had already discovered using the previous unbiased approach.

### Directed evolution of PhyB-Pif3 for independence from supplemental chromophore

In our fifth directed evolution campaign, we wished to improve the usability of the red-/far-red-light controlled optogenetic switch PhyB-Pif3. For this purpose, we put *CLN2* and *CLB5* under the control of the *GAL1* promoter and introduced single copies of *ADH1pr-PHYB-GAL4BD* and *ADH1pr-GAL4AD-PIF3* constructs, which controlled both cyclin genes in *trans* (Fig. 7 A). We screened for mutations allowing cell proliferation without supplementing PCB (Fig. 7 B). In ten out of ten mutants, which we analyzed by whole-genome sequencing, we found mutations in the *YOR1* (also known as *YCF1*) gene, of which five were nonsense mutations, two were single-nucleotide deletions, and one was a single-nucleotide insertion (Q64*, E421K, E538*, W570*, K676*, G713S, E1318*, A1306>, A1834>, A4690>AA).

**Fig. 7.**
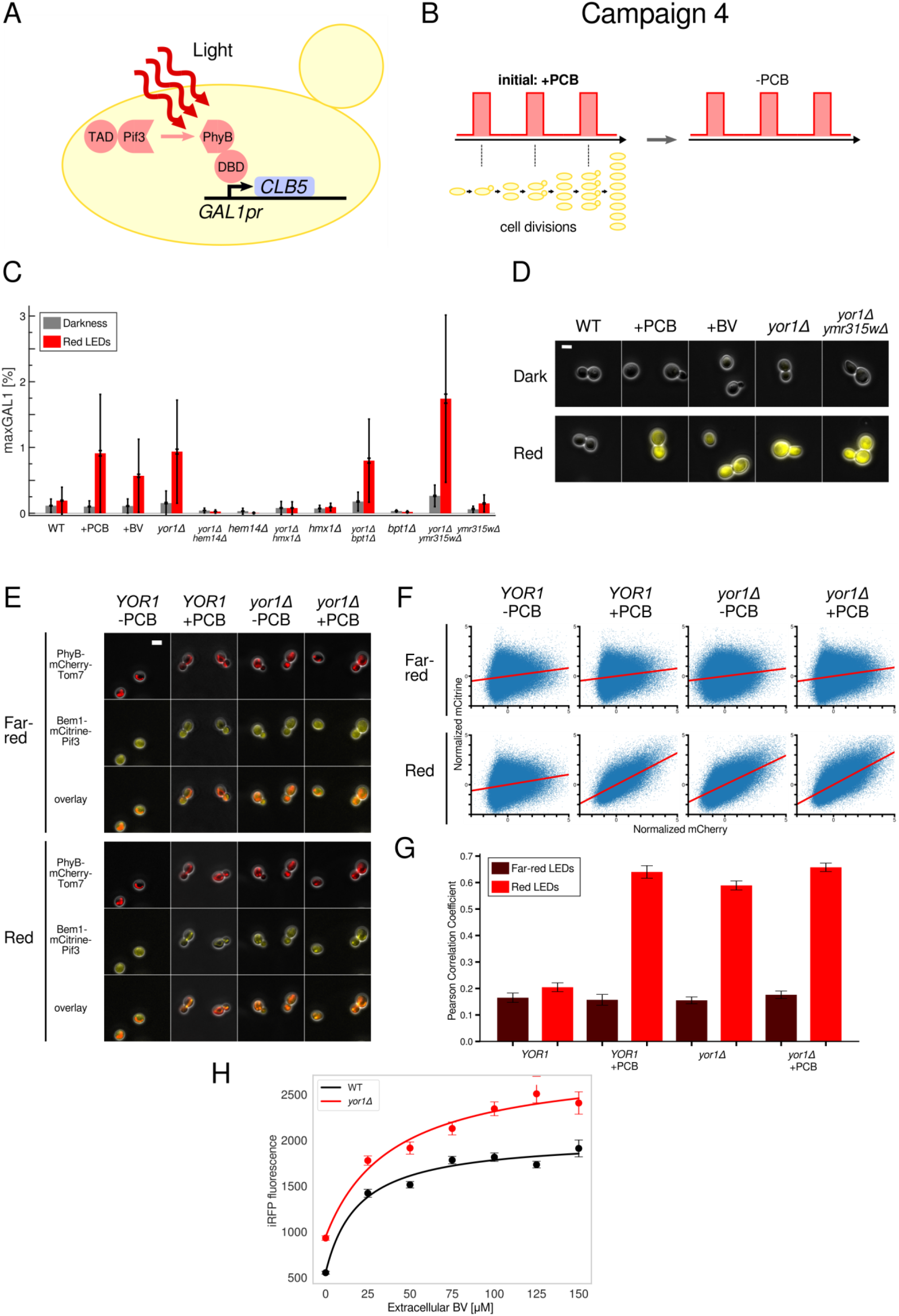
Directed evolution campaign 6 to lose the dependence of PhyB-Pif3 on supplemental PCB. A. Schematic of the wiring diagram relevant for the campaign. (*CLN2* and other cell cycle regulators omitted for clarity.) TAD, transcriptional activation domain; DBD, DNA binding domain. B. Schematic of the campaign. C. *GAL1pr-yEVenus* induction in the presence or absence of different genes involved in biliverdin metabolism and of supplemental biliverdin derivatives. The inner error bars represent the standard error of the mean (SEM); the outer error bars represent the standard deviation (STD). Numbers of analyzed cells are given in Supplementary Tables 8-10. D. Representative images of cells in experiments for panel C. E. Representative images of cells used for testing co-localization of Pif3 to mitochondrially anchored PhyB with or without *YOR1*. Cells were exposed to red LEDs, an image was taken, then cells were exposed to far-red LEDs for 30 sec, after which another image was taken. F. Scatter plot and linear regression fits for mCitrine versus mCherry fluorescence for each intracellular pixel. G. Pearson correlation coefficients for the linear regression fits in panel F. H. iRFP fluorescence in *YOR1* and *yor1Δ* cells at different extracellular biliverdin concentrations. All scale bars, 5 µm. Panels C-G, the supplemental PCB and BV concentrations were 30 µM and 100 µM, respectively. In panels G, H, error bars indicate the standard error of the mean (SEM). For the spectra of the LED lights see Supplementary Figure 1. The measured characteristics of PhyB-Pif3 with and without *YOR1* have been added to the Inducible Transcriptional Systems Database (https://promoter-benchmark.epfl.ch/)^88^.

We validated that a *YOR1* deletion suffices to obviate the need to supplement PCB for PhyB-Pif3 function in two reporter strains from other laboratories. We deleted *YOR1* in a strain from ref.^110^, in which PhyB-Pif3 drives the transcription of a fluorescent reporter (Fig. 7 C, D). In the absence of PCB, the fluorescent reporter responded as strongly in the *yor1Δ* background as with PCB and wild-type *YOR1*. Leakiness was comparable with and without *YOR1*. Since PhyB-Pif3 is frequently used to control the localization of proteins to different cellular compartments, we tested if the *yor1Δ* deletion sufficed to obviate supplemental PCB for this purpose as well. We deleted *YOR1* in a yeast strain from ref.^144^ in which PhyB is anchored to mitochondria, labeled with mCherry, and Pif3 is fused to a yellow fluorescent protein and Bem1 (PhyB-mCherry-Tom7, Bem1-mCitrine-Pif3). Exposing this strain with the *yor1Δ* deletion to red light led to colocalization of PhyB and Pif3 without supplemental PCB, comparable to *YOR1* cells with supplemental PCB in extent and reversibility (Fig. 7 E-G).

Yor1 is an ABC plasma membrane exporter^145^, a homolog of human cystic fibrosis transmembrane receptor (CFTR)^146^. We speculated that budding yeast produces a chromophore that can accumulate sufficiently for PhyB-Pif3 function unless Yor1 exports it out of the cell. Because PCB is a derivative of biliverdin in plants, we hypothesized that the chromophore was a precursor or a derivative of biliverdin, or biliverdin itself. To investigate this idea, we deleted two genes in the biliverdin biosynthesis pathway, namely, *HEM14*^147,148^, a protoporphyrinogen oxidase needed for heme biosynthesis, or *HMX1*^149^, a heme oxygenase that converts heme to biliverdin. Both genes turned out to be needed for *yor1Δ* to enable supplement-free use of PhyB-Pif3 (Fig. 7 C), suggesting that the chromophore is biliverdin or a biliverdin derivative. Further supporting this idea, we found that supplementing biliverdin instead of PCB suffices for PhyB-Pif3 function (Fig. 7 C). The recent discovery that PhyB can covalently incorporate biliverdin *in vitro* and interact with PIF6 provides mechanistic support for our finding^150^. (The ability to substitute lower-cost biliverdin for PCB for use with budding yeast is by itself advantageous.) Additionally, the biliverdin sensor iRFP^151^ showed about 70% higher fluorescence in *yor1Δ* versus wild-type *YOR1* cells (Fig. 7 H). To characterize the efficiency with which Yor1 may export biliverdin, we quantified the difference in iRFP fluorescence with and without *YOR1* at different extracellular biliverdin concentrations. The gap in iRFP fluorescence indicates that Yor1 can maintain a substantial difference between intra- and extracellular biliverdin. Together, our results strongly suggest that biliverdin or a biliverdin derivative are exported out of the cytosol by Yor1 and accumulate sufficiently for PhyB-Pif3 function with *yor1Δ* deletion.

Going further, we searched for other proteins that might reduce the levels of biliverdin in yeast cells, in addition to Yor1. Eliminating such proteins would further boost PhyB-Pif3 function in the absence of a supplemental chromophore. Our first strategy was to find potential biliverdin-binding proteins. We used Foldseek^152^ to find yeast proteins with structural homology to human biliverdin reductase A (BLVRA). This search identified the poorly characterized yeast protein Ymr315w. The ligand identification tool AlphaFill^153^ showed that Ymr315w was likely to bind biliverdin. In line with these computational predictions, deleting *YMR315W* enhanced PhyB-Pif3 transcriptional output in a *yor1Δ* background without supplementing a chromophore (Fig. 7 C). (The *ymr315wΔ* deletion alone had a mild effect, which is why our directed evolution campaign may not have caught it.) In a second approach, we tested whether another ABC transporter could also export the PhyB-Pif3 chromophore. Among 16 characterized ABC transporters in yeast, Bpt1, a vacuolar importer, has the greatest sequence similarity to Yor1^154^. However, deleting *BPT1* did not enable or improve supplement-free use of PhyB-Pif3 (Fig. 7 C). These results further support our finding that specific cellular pathways can be modified to enable supplement-free PhyB-Pif3 in budding yeast. Optovolution, being an *in vivo* directed evolution method, can access such solutions which originate in the biology of the cell. Depending on the goal, these solutions can be desired or, if undesired, these avenues can also be foreclosed (**Discussion**).

### Directed evolution of a light- and doxycycline-inducible AND gate

In our sixth evolutionary campaign, we applied optovolution to a non-optogenetic system. First, we engineered an AND gate based on the transcription factor rtTA, which is controlled by doxycycline. To make rtTA dynamically sensitive to transcriptional input, we destabilized it by fusing its N-terminus with a PEST degron. When *PEST-rtTA* was transcribed under the control of the El222/*LIP* system, transcription from the rtTA-controlled promoter *tetOpr* was very weak.

We wished to evolve variants of the logic gate with stronger on (1) state activity. So, we introduced *tetOpr-CLN2*, *tetOpr-CLB5*, and *PEST-rtTA* into the Dajbog1 strain (Fig. 8 A). We then evolved the system by exposing cells to short pulses of blue light, 2 sec on every 100 min, under which they could not proliferate (Fig. 8 B). Before evolution, light pulsing 30 times 30 sec on, 30 sec off every 100 min and doxycycline allowed cell proliferation, confirming the AND gate functionality prior to directed evolution (Fig. 8 C). However, the *tetO* promoter is very leaky in budding yeast (*tetOpr-yEVenus* expression was at 2% maxGAL1 without *rtTA*) so that even in darkness and without doxycycline, Dajbog1 proliferated, albeit much more slowly. This is why we used the more leakiness-tolerant Dajbog1 strain in this campaign and screened for mutants as soon as small colonies emerged under 2 sec light pulses every 100 min.

**Fig. 8.**
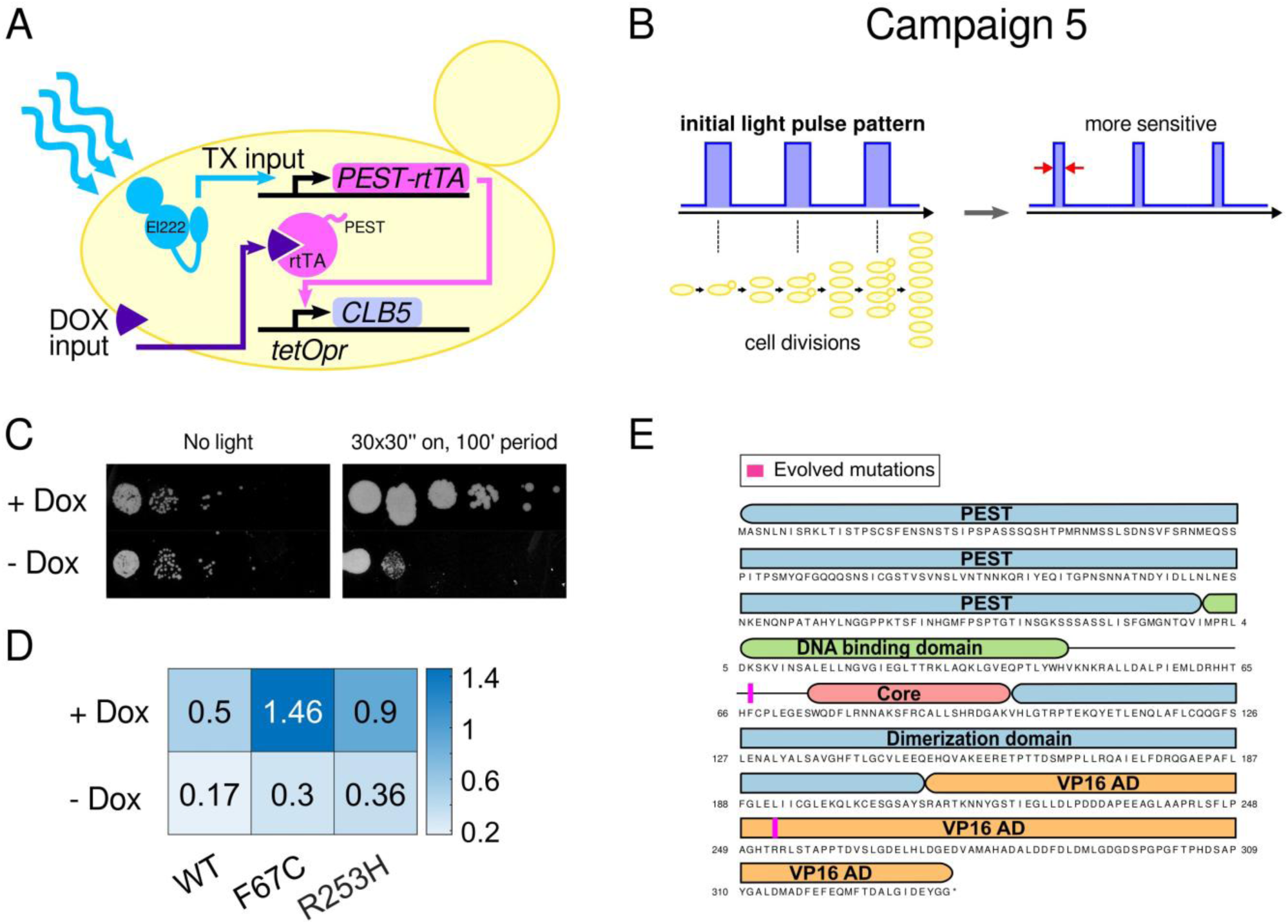
Directed evolution campaign 6 to improve the AND logic gate function of PEST-rtTA. A. Schematic of wiring diagram relevant for campaign. TX, transcription; DOX, doxycycline. B. Schematic of the campaign. C. Spot test for wild-type Dajbog1 strain with PEST-rtTA system with ten-fold dilutions. D. Expression level of *tetOpr-yEVenus-PEST* under the control of the original or evolved PEST-rtTA in maxGAL1 units. For lowering experimental variability, the original and mutant *PEST-rtTA* were expressed from the *PGK1* promoter. Residue numbers are with respect to +1 methionine of rtTA. Numbers of analyzed cells are given in the Supplementary Table 11. E. Positions of mutations in domain diagram of PEST-rtTA.

In principle, mutations in any part of the system could lead to the increased activity. Given that we have already extensively characterized the mutations that boost the activity of the El222/*LIP* optogenetic system, we now focused on screening for mutations in *PEST-rtTA* (**Discussion**). We discovered two mutations, F67C and R253H (numbering is with respect to +1 methionine in rtTA).

To characterize the *PEST-rtTA* mutants, we cloned and introduced them as single-copy constructs driven by *PGK1pr* into a reporter strain with a single copy of *tetOpr-yEVenus-PEST* (Fig. 8 D). Both mutations showed increased activity in the presence of doxycycline. The mutated residues are located between the DNA binding domain and the rtTA core domain or in the VP16 transcriptional activation domain, respectively (Fig. 8 E). Relatedly, in previous directed evolution experiments of rtTA, F67S was identified as increasing the sensitivity of rtTA (without a degron) to doxycycline^155^.

These results demonstrate how the light-driven selection pressure in optovolution can be leveraged for the evolution of non-optogenetic proteins with dynamic, multi-state, and computational properties.

## Discussion

### Two key ideas underlying optovolution

The primary advance of our directed evolution paradigm is the coupling of the POI output to an element of cell cycle control that has to oscillate for cellular replication in the specially engineered Dajbog genetic background. This obviates interventions needed to screen phenotypes or to switch between selection mechanisms after each round of evolution. Instead, both 0-to-1 and 1-to-0 switching of the POI are effectively screened every ≈90 min by the cell cycle control machinery *in vivo*. In addition, the coupling of the desired POI properties to organismal fitness automatically ensures that the evolved variants are functional *in vivo*. Additional *in vitro* screening is not needed. These advantages of our method arise from leveraging cell cycle control, to our knowledge, for the first time for directed evolution.

Secondarily, we use optogenetics to cycle through the inputs of the POI. Fundamentally, light control is powerful for directed evolution because it allows communicating with cells rapidly, conveniently, and for extended periods. Thus, new kinds of selection pressures can be created, which require dynamic interaction with cells. As far as we know, this is the first time that optogenetics has been leveraged for continuous directed evolution.

Light control also allows straightforwardly shaping the fitness landscape of the POI by tuning the selection pressure. Different evolutionary outcomes can be achieved depending on the light exposure protocol. Input-output relationships can be challenged by administering light input that is too weak or too strong. Directed evolution can be expected then to improve the input-output by mutational adaptations. Similarly, engaging and releasing the POI early or late forces adaptations toward improved POI kinetics. We used the same strain, with *EL222* directly controlling both *CLB5* and *CLN2* in *trans*, for four directed evolution campaigns: enhancement of the 1-output activity of El222 (on agar plates or liquid culture), suppression of the 0-output activity, or green responsiveness.

### Tolerance toward poor initial POI functionality

A general challenge is that a POI may be so poor with respect to a desired function that the directed evolution campaign cannot even be initiated. This is mitigated in two key ways in our method: The selection pressure can be easily tuned by changing the light pulse regime, e.g., shorter, longer, dimmer, or brighter light pulses. Furthermore, the training wheel plasmid allows proliferation in the absence of any POI function, so that directed evolution of the POI can be initiated concomitantly with selection for loss of the training wheel plasmid on 5-FOA.

### Efficiency and targeted mutagenesis

Mutations in the POI gene evolved efficiently and bypass mutations were either rarely observed in Dajbog2 (*sic1Δ*) or were reasonable in number to quickly find the mutation of interest with Dajbog1 (wt-*SIC1*). The strains can in principle be further optimized to suppress mutations in the budding yeast genome. After identifying such bypass mutations, one can effectively prevent them by inserting additional copies of the gene. This strategy takes advantage of the tolerance of yeast toward multiple copies of most of its genes^156,157^ and the extremely low likelihood of mutations occurring in different copies of the same gene. We demonstrated this strategy with multiple *LIP-CLB5* copies in our second directed evolution campaign. Should bypass mutations occur in non-essential genes, one can in principle simply prevent them by deleting the gene.

The deep literature on cell cycle control can be used to guide future improvements of optovolution. We leveraged the extensive genetic screens for suppressors of cell cycle arrest performed in the past to create a genetically robust platform. Screens have only revealed *sic1Δ* to bypass *cln1-3Δ* arrest.^133^ This bypass relies on *CLB3-6*, which are absent in the Dajbog strains. (It should be noted that *cln1-3Δsic1Δ* cells proliferate poorly, suggesting that bypass mutations do not necessarily affect directed evolution experiment in practice.) Bypass mutations in Cdk1 can be practically ruled out since at least five mutations are required to run the cell cycle oscillator in a *cln1-3Δ clb5,6Δ* background.^131^ It is unknown whether even these quintuply mutated Cdk1 could run the Dajbog strains, which have additional deletions of *CLB3,4*.

Another potential way to increase the efficiency of our method is to take advantage of the orthogonal advances in targeted mutagenesis methods. These are especially needed when mutations elsewhere in the genome can readily bypass the desired POI functionality. However, Dajbog2 was genetically tight, so it was not necessary to target mutations to the POI gene to achieve our goals. Using *pol3-L523D*, we expected to increase the genome-wide mutation rate to 0.5×10^-7^ per base-pair per generation. This means that in tens of milliliters of log-phase yeast population, about all single mutants of the POI gene should have been present. Straightforward ways to further increase the mutation rates globally involve the use of DNA damaging radiation and chemicals, although not all nucleotide mutations can be attained in this manner. Another common method to achieve very high rates of targeted mutations at the start of the directed evolution campaign is to introduce the POI on mutagenized plasmid libraries, e.g., by error-prone PCR. Our method will also presumably benefit from incorporation of targeted *in vivo* mutagenesis systems such as OrthoRep^10^, MutaT7^12,158^, or Cas9-^159^ or transposon-mediated systems^17^, especially when evolutionary goals require multiple mutations at the same time.

### Ease of use

After extensive engineering and testing, the Dajbog strains can now be used without further modification, in principle, including different variants (*SIC1*/*sic1Δ* and with multiple *CLB5* copies). The steps to setting up optovolution (Fig. 3) resemble those of other *in vivo* directed evolution methods, i.e., *CLB5* and *CLN2* must be placed under the control of the POI, similarly to non-continuous directed evolution methods with fluorescent proteins or *URA3*. The key difference is that cumbersome alternating selection mechanisms are obviated by the cell cycle oscillator. The training wheel plasmid further helps exploring which pulsing regimes exert selection pressure, and can be cleanly removed by counter-selection. With Clb5 (and Cln2) being critical for a key biological process and cell cycle phenotypes extensively characterized and easy to interpret, the use and debugging of optovolution is straightforward. Only standard equipment such as LED panels and incubators as well as knowledge of standard techniques in budding yeast maintenance and transformation are needed.

### Limitations of optovolution

Limitations on types of POI input: Our directed evolution of PEST-rtTA demonstrated that the POI itself need not be light sensitive. To take advantage of intervention-less optovolution, a non- optogenetic POI has to be wired in a multi-step signaling cascades (optogenetics → POI → Clb5,Cln2). Options for light control of the POI are diverse and rapidly expanding, including light- controlled transcriptional input (used for PEST-rtTA), light-controlled protein-protein interaction (used for PhyB-Pif3), light-controlled degrons, kinases, and mRNA targeting systems. POIs that can only be controlled with chemicals are compatible with our platform but would require intervention, albeit only to cycle through the chemical POI inputs but not to change the selection mechanism.

Limitations on types of POI output: We demonstrated optovolution by evolving POIs that control the transcription of *CLB5* (and simultaneously *CLN2*): two transcription factors, El222 and PEST-rtTA, and an inducible protein-protein interaction system, PhyB-Pif3, which was engineered to control transcription using the yeast two-hybrid system. Many existing continuous directed evolution methods^14,34,35^ were similarly initially applied to POIs that control transcription, presumably because transcriptional control can be created in a multitude of ways: The well-established two-hybrid method converts protein-protein interactions into transcriptional output^160^. CRISPR-Cas systems were in many cases evolved by blunting the endonuclease activity of Cas proteins and coupling to transcriptional output.^25,26,161,162^ GPCRs were similarly screened by coupling downstream signaling to transcription.^163–165^ Thus, transcriptional control does not limit the POI to transcription factors. Furthermore, the crucial element of our method is control of Clb5 activity, which can in principle be coupled to POI activity in other ways.

Limitations on targeting POIs: Even if the cell cycle-coupled selection mechanism is genetically robust, genomic mutations can emerge that are in cellular pathways affecting POI function. An example in our study is *YOR1*, whose mutations allowed PhyB-Pif3 to respond to red light in the absence of PCB. We considered the mutations in *YOR1* to be sufficiently surprising and useful to report in their own right as an endpoint of that campaign, instead of seeking to suppress them.

Requirement of minimal computational functionalities: Optovolution is fundamentally not suited for proteins whose state switching cannot be controlled even by way of intermediates, or for proteins that cannot control Clb5 activity. Essentially, the POI must meet the minimum requirements for computational functionality: controllable inputs and an output that can be read out, specifically, by coupling to Clb5. For example, proteins that stochastically switch between states without input or that are not coupled to an activity that could ultimately control Clb5 are not evolvable with optovolution.

Extendibility to other organisms: Budding yeast provides many advantages for directed evolution of dynamic, multi-state, and computational functionalities, including a relatively short generation time, ease of genetic engineering, and a vast literature on genetics, growth, metabolism, and cell cycle control. As a eukaryote, budding yeast shares many homologous genes with mammals. While this makes budding yeast a well-appreciated platform for protein engineering and subsequent transfer to mammalian cells, proteins optimized for budding yeast may not immediately function as expected in mammalian cells. Further work is often required for adaptation to new hosts^166^. Our *yor1Δ* findings motivate a targeted screen of human ABC transporters and genes in the biliverdin metabolism pathway for enabling PhyB-Pif function in the absence of PCB^167^. To transfer our El222 mutants to mammalian cells, one should consider that LOV proteins are temperature sensitive, potentially requiring adaptation to elevated temperatures^54^. While all of our directed evolution campaigns were conducted at 30 °C, which maximized budding yeast growth, the organism is known to tolerate 37 °C as well, with only a mild (<10%) slow-down in the cell cycle^168^. Thus, optovolution can be used to evolve POIs closer to use in mammalian cells. Nevertheless, additional improvements of the POI may be needed beyond temperature adaptation.

### Green responsive El222 and *yor1Δ*-enabled PhyB-Pif3 function

Directed evolution methods rapidly assess an enormous number of mutations (determined by the mutation rate, spectrum, and number of cells) for a desired POI property, beyond what is currently feasible in standard site-directed mutagenesis and *in vitro* analyses. Optovolution substantially simplifies this powerful approach for dynamic, multi-state, and computational functionalities. We used it to uncover a number of practically useful and surprising mutations.

Rescuing Dajbog cells under pulses of green light, T83A and G80R El222 mutants, cloned into our reporter strains, showed responses to ≈500-520 nm green light. Comparing the 450±5 nm blue light response at different intensities to the 510±5 nm green light response (Fig. 6 J-M), we observed that the latter was lower but comparable when non-saturating blue light was applied. In contrast, El222 was not responsive to any of the light we administered except to the highest intensity of blue light used in this set of experiments (20%), preventing conclusions to be drawn regarding differential color sensitivity. The A157V mutation, which is far from the chromophore, produced a response to all blue light intensities in these experiments but, again, no detectable response to our highest-intensity green light, which was quite bright to the human eye. The *in vivo* data show that T83A and G80R gained green responsiveness that is not simply a consequence of a scaled overall light responsiveness.

Our biophysical characterization of T83A shows higher relative sensitivity in the 500-520 nm range compared to 450 nm than in wild type, as assessed by the absorption spectrum and by comparing the activation kinetics in blue and green light (Fig. 6 N-P). Changes to the relative sensitivity to green and blue light are novel and surprising. In contrast, we determined the purified G80R variant to be overall more light sensitive, which at least partially accounts for the greater green sensitivity *in vivo*. In any case, these results highlight the value of *in vivo* directed evolution driving protein engineering alongside biochemical and other approaches: Based on the *in vitro* alone, especially of G80R, the starkly different responsiveness to green light *in vivo* would have been difficult to predict. Irrespective of the underlying mechanism, this observation motivated the creation of a circuit that enables a green-only response (Fig. 6 R), which also demonstrates that the difference in responsiveness to green light suffices in practice. Green responsiveness is not readily found in other LOV domains, for example, it is absent in the phototropin *At*LOV2 module (Fig. 6 T).

For the second set of unexpected results, the findings that knock-out of *YOR1* obviates the need for supplemental PCB, we found a well-constrained mechanistic explanation. In particular, the genetic analysis that deletion of *HEM14* and *HMX1* abolish the ability of *yor1Δ* to support PhyB-Pif3 function, together with the established role of Yor1 as a plasma membrane transporter, suggest that biliverdin or a biliverdin derivative, naturally produced by budding yeast, serves as chromophore. Further work is needed, however, to pinpoint the identity of the chromophore precisely.

### Outlook for evolving computational functionalities

Given the appropriate selection pressure, directed evolution can solve challenging engineering problems and uncover unexpected solutions -- such as green-responsive El222 mutants or PCB independence conferred by *yor1Δ*. Here, we present the first method for continuous directed evolution of dynamic, multi-state, and computational protein functionalities. Such capabilities are fundamental to signal transduction -- and thus to life. While much of cellular biochemistry computes decisions, it had been unclear how continuous directed evolution could be adapted to tune these properties. Apart from interest in this as a fundamental problem, a solution is also needed for biological engineering since dynamic, multi-state, and computational properties are of intense current interest, especially, in protein design^169–173^.

More advanced computational capabilities involve more inputs for more complex input-output relationships. Using optovolution, different colors (UV^174,175^, blue^103^, green^176^, red^108,177^) and optogenetic systems allow administering different inputs to the POI and cycling through input combinations during directed evolution.^99–102,176,178,179^ The number of optogenetic effectors is increasing, enabling different kinds of input to the POIs^180^. Single-protein computers would have numerous advantages over multi-gene circuits, including simpler genetic introduction into cells and presumably faster kinetics. However, rational engineering of proteins often leads to poorly performing molecules, underscoring the need for directed evolution approaches. Our motivation for engineering and evolving an AND gate involving PEST-rtTA was to demonstrate that optovolution is well-suited for such systems, beyond light-sensitive proteins.

Optovolution is particularly timely for the revolution that is currently taking place in artificial intelligence-based protein design, in which there is substantial interest in dynamic, multi-state, and computational properties^29,181–183^. Prototypes that have been computationally designed can be screened and improved in a high-throughput manner with optovolution. Additionally, machine-learning guided methods will rely on experimentally validated training sets, which can be further enriched through directed evolution. We expect optovolution to provide solutions to previously intractable engineering problems and pave the way for more sophisticated proteins.

## Methods

### Media

We used synthetic complete media^184^ (SCD) with methionine (+Met) or without methionine (-Met) and supplemented with 2% (w/v) glucose. For preparing solid SC plates, we used 2% agar. For experiments with 5-FOA, we used a standard recipe with 0.1% 5-FOA^71,185^. For controlling and evolving the transcriptionally and doxycycline-regulated AND gate (PEST-rtTA), we used 30 µM doxycycline diluted from a 1000x stock in water (Sigma, USA). For characterization of the PEST-rtTA variants, we used 10 µM doxycycline. For experiments with PhyB-Pif3, we used 30 µM PCB, diluted from a 1000x DMSO stock (Sychem, Germany) when supplying exogenous PCB.

### Plasmid construction

Cloning was performed using restriction enzyme digestion and ligation with T4 ligase (New England Biolabs) or Gibson assembly (New England Biolabs). Plasmids were propagated in DH5alpha *E. coli* in LB medium supplemented with ampicillin. See Supplementary Table 1 for a list of the plasmids.

### Strain construction and preparation for directed evolution

Transformations of budding yeast were performed using the high-efficiency LiAc/DNA carrier/PEG (polyethylene glycol) protocol^186^. All strains are derivatives of W303 (*leu2-3,112 trp1-1 can1-100 ura3-1 ade2-1 his3-11,15*). A list of strains is provided in Supplementary Table 2.

The Dajbog1 (*cln1-3Δ clb3-6Δ*) and Dajbog2 (*cln1-3Δ clb3-6Δ sic1Δ*) strains proliferated in the absence of light pulses in SCD-Met medium plates due to the presence of the *URA3*-marked training wheel plasmid pRD-TW1 (*MET3pr-CLN2, CLB5pr-CLB5, CEN/ARS*), which was built using the pRS416 vector^187^. (Lack of methionine induces *MET3pr-CLN2*.) For the different directed evolution campaigns, we built additional strains starting with the Dajbog strains. To evolve *EL222* (first four campaigns), we transformed the strains with *LIP-CLB5*, *LIP-CLN2*, and *PGK1pr-EL222* constructs. For the evolution of the PhyB-Pif3 system (fifth campaign), we transformed them with *ADH1pr-PHYB-GALBD*, *ADH1pr-GAL4AD-PIF3*, *GAL1pr-CLN2*, and *GAL1pr-CLB5* constructs. Lastly, for evolving the AND gate (sixth campaign), we transformed the strains with *PGK1pr-EL222*, *LIP-PEST-rtTA*, *tetOpr-CLN2*, and *tetOpr-CLB5*. We confirmed transformations by PCR.

To prepare the strains for directed evolution experiments, we selected for the loss of the training wheel plasmid as follows: First, cells were spread on fresh SCD+Met plates and placed under pulsing light. Methionine shuts off *MET3pr*, reducing any advantage that the training wheel may confer and thus increasing the fraction of cells that lose the training wheel plasmid. Then, we transferred cells to SCD+Met+5-FOA plates under the same pulsing light regime, so only cells that had lost the training wheel plasmid could proliferate. We picked isolated colonies in which we confirmed the loss of the training wheel plasmid by the lack of growth in darkness on plates lacking methionine. From then on, the strains were propagated under pulsing light.

After directed evolution, we characterized the evolved POIs by cloning the mutated POI genes into a backbone for single-copy integration. A W303 strain was transformed with these plasmids as well as plasmids for single-copy integration of constructs in which the transcription of a fluorescent protein was driven by *LIP* (in case of El222) or *tetOpr* (PEST-rtTA). Integration of single copies was confirmed by PCR in all transformants. For measuring the transcriptional rate of PhyB-Pif3 as a split Gal4 system, we used the strain from ref.^110^

### Light conditions

Throughout, we handled cells sensitive to blue and green light under red light and red-responsive cells as well as PCB under green light. Plates were otherwise always wrapped in aluminum foil.

For administering blue light to induce El222, we used commercial blue light LED panels (model 884667106091218, HQRP, USA). The panels were placed inside incubators at a distance of 20 cm above the agar plates or a spinning wheel of liquid cell culture tubes. For evolving green-responsive El222 variants, we used LEDs with peaks around green (525 nm, model LTL3H3TGRADS), yellow-green (555 nm, model WP710A10PGC), yellow (575 nm, model 3SUGC-F), orange (615 nm, model TLCO5100), and red (660 nm, model TLDR5800) (Mouser, Switzerland). For characterizing the green-responsive El222 variants, we used band-pass filters FBH450-10, FBH500-10, FBH510-10, FBH520-10, FBH530-10, and FBH650-40 (Thorlabs) placed in the path of the microscope’s diascopic light (Supplementary Figure 1). For administering red and far-red light that switches PhyB-Pif3, we used red light panels (model TLDR5800, Mouser, Switzerland) with a peak around 650 nm and custom-made printed circuit boards with 750 nm far-red LEDs (model LED750-33AU, Roithner Lasertechnik, Austria).

The spectral characteristics of the light sources, measured using an ST-VIS-25 spectrometer (Ocean Insight, USA) are shown in Supplementary Figure 1.

To control the LEDs or lasers, we used an NI USB 6501 (Texas Instruments, USA) I/O device. The system was run using a custom Matlab script (**Code availability**). The regime of pulsing light could be adjusted with a precision of less than one second.

### Quantification of the transition from *cln1-3Δ* arrest with optogenetic cell-cycle control

To score the portion of cells that bud in response to a pulse of light (Fig. 2 B), we tested *cln1-3Δ MET3pr-CLN2* and *cln1-3Δ clb3-6Δ pRD-TW1* cells with additional *EL222* and LIP-controlled Cdk1 cyclins, as indicated. After loading cells into a microfluidic chip (CellASIC ONIX Y04C-02, Sigma-Aldrich Chemie GmbH, Germany) and growing them for 4-5 hrs, we supplied methionine, which shuts off *MET3pr*. Two hours after methionine had been added, cells were large and unbudded (G1 arrest). Different optogenetic constructs (*LIP-CLB5* and *LIP-CLN2*) were activated by using diascopic light from the microscope set to 80% of maximal intensity for 20 min or 40 min, depending on the experiment. Cells were imaged every 10 min.

### Optovolution experiments on agar plates

Prior to evolution, cells were propagated under the light pulsing regime that allowed proliferation on SCD agar plates until growth saturation. For the El222 campaigns, this was 5 times 25 sec on, 35 sec off every 100 min; for the PhyB-Pif3 campaign, 40 min on, 60 min off; for the PEST-rtTA campaign, 30 times 30 sec on, 30 sec off every 100 min. To increase the mutational diversity of the population, cells were then mutagenized by exposing them for 10 sec to UV light with 254 nm emission peak (model G30T8, Sankyo Denki, Japan). We scraped cells from one plate and resuspended them in 1 mL of liquid SCD medium prewarmed to 30°C. Then, we plated 100-200 µL of cell culture on fresh agar plates and chose the desired selection pressure by adjusting the light pulse regime. After two to three days, we picked isolated colonies and restreaked them on fresh SCD plates. After two days of growth of the streaks, we extracted genomic DNA and identified the mutations by Sanger sequencing the POI gene or whole-genome sequencing.

### Long-term continuous optovolution in liquid medium

As in the agar plate experiments, cells were propagated on SCD plates and UV-mutagenized. We scraped cells from one plate and resuspended them in 1 mL of liquid SCD medium. We added 200 µL from this suspension to 100 mL of liquid SCD medium. We used transparent 250 mL Erlenmeyer flasks (VWR, Switzerland). Cell cultures were placed in an incubator shaker (Infors HT, Switzerland) set to 250 rotations per minute and 30°C. For administering pulsing light, we used panels of blue LEDs mounted on a custom rack inside the incubator.

To prevent the cultures from reaching saturation, we diluted them in warm SCD medium to a final OD_660_ of approximately 0.04-0.1 every 12 hrs, unless the OD had not changed. We performed the experiment for a week, after which we proceeded to Sanger sequencing bulk culture genomic DNA for the POI gene.

### Mutation identification

To screen for mutations in the genes of interest, we performed Sanger sequencing (Microsynth AG, Switzerland) using primers that span the coding sequence of interest. The identified mutations were confirmed by sequencing the plasmids after cloning the POI gene.

For identifying mutations in the genome, we performed paired-end whole genome sequencing of the evolved and ancestral genomes. The sequencing was performed using 150 bp long reads on the DNBseq platform with a total data output of 1 Gb. The ancestral and the evolved reads were then mapped onto the W303 reference genome using Bowtie 2. For variant calling, we used BCFtools.

The mutations were called if they appeared in the evolved but not the ancestral genome, and if they were sequenced with a depth of at least 10.

### Variant characterization

In all cases, strains were grown to saturation for 18-20 h, diluted, and then allowed to reach log phase. Transcriptional activities were scaled to the maxGAL1 unit, where 1 maxGAL1 corresponds to the steady-state expression of *GAL1pr* in synthetic complete medium containing 3% raffinose (w/v) and 2% galactose (w/v)^88^.

For characterizing the light-sensitized and *in vivo* green-responsive El222 variants, we cloned the mutant *EL222* genes into plasmids that inserted as single copies into the genome of strains that contained a single-copy *LIP-ymScarletI* reporter. While *LIP-ymScarletI* was the reporter of choice for characterizing light-sensitized variants to avoid excitation light inducing the mutant El222, it has a comparatively lower sensitivity than a *yEVenus* reporter. Hence, for carefully characterizing the leakiness of wild-type El222 and low-leakiness variants, we introduced them into a strain containing a single-copy *LIP-yEVenus* reporter. All variants were induced in log phase in liquid culture for 6 h. The light-sensitized or low-leakiness variants were tested under two different light pulse regimes, high or low time-averaged LED intensity (Fig. 5). The low LED regime corresponds to the blue LEDs (Supplementary Figure 1) pulsed for 10 sec every 5 min. The high LED regime corresponds to 30 sec on, 30 sec off every minute. For characterizing the dynamics of the evolved variants with timelapse fluorescence microscopy, we used variants of the reporters in which the corresponding fluorescent proteins are fused to the PEST degron, reducing the half-life of the reporters. From the measured fluorescence values, we subtracted the fluorescence levels of cells containing only the corresponding *LIP*-based reporter, which showed elevated fluorescence levels compared to wild-type (no-reporter) cells (Supplementary Figure 3).

For characterizing the effect of *yor1Δ* in combination with the PhyB-Pif3 system, we used a strain from ref. ^110^ with a *GAL1pr-Venus* reporter. In case PCB was used, cells were incubated with it in darkness for 1 hr before exposure to red light. We used 30 sec on, 30 sec off red light with an emission peak around 650 nm delivered for 6 h to cells grown in SCD liquid medium tubes on a spinning wheel. After that, the level of expression was measured using fluorescence microscopy. Previous measurements of *GAL1pr* activity in glucose showed that it does not exhibit leakiness that is distinguishable from wild-type cells without a reporter^88^. Hence, from the measured values, we subtracted the background fluorescence of a wild-type strain. To convert the expression levels to maxGAL1 units, we measured the steady-state expression level of *GAL1pr-Venus* in the same strain by growing it in SC medium with 3% raffinose (w/v) and 2% galactose medium (w/v).

For quantifying the activity of PEST-rtTA mutants, we introduced them into strains with a *tetOpr-yEVenus-PEST* reporter. We grew the strains with or without doxycycline for 6 hrs and then imaged them by fluorescence microscopy. From the measured values, we subtracted the corresponding background fluorescence values of the strain containing only a *tetOpr-yEVenus-PEST* reporter in the corresponding medium.

### Microscopy and image analysis

Images were recorded using a Nikon Ti2-E (Nikon, Switzerland) microscope equipped with a 60x oil, N.A. 1.40 objective and a Hamamatsu Orca-Flash 4.0 camera (Hamamatsu, Japan). The microscope was controlled using NIS-Elements software and the Nikon Perfect Focus System. When performing timelapse microscopy to monitor the dynamic properties of El222, we induced with diascopic light, as described previously. For green-responsive mutants, we inserted bandpass filters into the light path of the microscope’s diascopic light. The characteristics of this light are presented in Supplementary Figure 1. The diascopic light was set to the intensities indicated in the figures. Image analysis was performed using YeaZ^137^, a deep neural network for analyzing images of yeast cells.

### Expression and purification of El222 variants

El222 variants were produced by heterologous expression in *E. coli* and chromatographic purification according to previous expression protocols for El222^139^ and other LOV receptors^188^. In brief, a pET29b(+) vector encoding the El222 gene with an N-terminal His_10_ tag and under the control of a T7-lacO promoter was used^139^. The mutations G80R and T83A were introduced into this vector by site-directed mutagenesis. *E. coli* CmpX13 bacteria^189^ harboring the requisite plasmid were grown at 37 °C and 225 rpm agitation in 2× 800 mL lysogeny broth (LB) medium supplemented with 50 µg mL^-1^ kanamycin and 50 µM riboflavin. At an optical density at 600 nm of around 0.6, 1 mM isopropyl β-D-1-thiogalactopyranoside was added, the temperature was lowered to 16 °C, and the incubation continued overnight. All subsequent steps were carried out under red safe light. Cells were harvested by centrifugation and resuspended in 20 mM Tris/HCl pH 7.5, 300 mM NaCl; to promote solubility, in the case of the T83A variant, 500 mM of the osmolyte arginine was included in all buffers. Following lysis by sonication and centrifugation to remove cell debris, the supernatant was applied to a Co^2+^ immobilized metal ion chromatography (IMAC) and eluted by imidazole. The eluted fractions were analyzed by denaturing polyacrylamide gel electrophoresis and pooled based on protein content and purity. To remove the His_6_ tag, TEV protease was added, followed by dialysis into buffer without imidazole. The sample was again applied to an IMAC column, dialyzed into storage buffer (20 mM Tris/HCl pH 7.5, 30 mM NaCl, 10% (v/v) glycerol) plus 500 mM arginine for T83A, and concentrated by spin filtration.

### Absorption-spectroscopic analyses of El222 variants

All spectroscopic measurements were carried out on an Agilent Cary 60 UV/vis spectrophotometer at a temperature of 25 °C maintained by a circulating water bath. To this end, the El222 samples were diluted into 20 mM Tris/HCl pH 7.5, 500 mM arginine, 30 mM NaCl, 10% (v/v) glycerol. Spectra recorded on the dark-adapted El222 variants served to determine the concentration of the holoprotein based on a molar extinction coefficient of 12,500 M^-1^ cm^-1^ at 450 nm for the absorbance of the flavin-nucleotide cofactor. To record spectra of the light-adapted state, the sample was exposed to an LED with peak emission at 470 and a light power of 36 mW cm^-2^ measured at the position of the protein sample using a Newport model 842-PE power meter and a 918D-UV-OD3 silicon photodiode.

Dark-recovery kinetics were recorded at 25 °C by monitoring the return from the light-adapted state at a wavelength of 450 nm, corrected by the absorbance at 600 nm to account for baseline shifts. The resultant data were fitted by single-exponential functions using the Fit-o-mat software to determine the rate constant *k*_-1_ of dark recovery^190^.

To evaluate the forward kinetics of photoactivation by blue and green light, respectively, dark-adapted El222 samples were irradiated through (450 ± 5) nm and (510 ± 5) nm filters (Thorlabs) using a fiber-optic lamp. Measured at the sample position, the area intensities of the 450-nm and 510-nm light amounted to 0.75 mW cm^-2^ and 1.5 mW cm^-2^, respectively. The gradual transition to the light-adapted state during illumination was tracked by absorbance measurements at 450 nm, corrected by the reading at 600 nm. Control experiments showed that the probe light of the spectrophotometer did not promote detectable photoactivation in itself. The resultant absorbance data were fitted to single-exponential functions to determine observable rate constants *k*_o_. As recovery to the dark-adapted state takes place during the experiment, the rate constant *k*_o_ is the sum of *k*_-1_ (see above) and the light-dependent forward rate constant *k*_1_ of photoactivation.

### Statistics and reproducibility

No statistical method was used to predetermine sample size. The values in the heatmaps in Figs. 4 and 6 are rounded based on the first significant digit of the standard error of the mean.

### Data and material availability

The characteristics of the systems measured in this study have been added to the Inducible Transcriptional Systems Database (https://promoter-benchmark.epfl.ch/)^88^. Any additional data that support the findings of this study are available from the corresponding author upon request.

### Code availability

The Matlab code for controlling the light pulses is available at: https://github.com/rahi-lab/optogenetic_controller_script/.

## Acknowledgments

We thank Roxane Dervey for help with strain construction and media preparation; Dr. Kyle Douglass, Dr. Hélène Lambert, Dr. Matthew Lycas, and Prof. Suliana Manley for sharing resources; Prof. Peter Quail, Prof. Hana El-Samad, and Dr. Brian Graziano for providing PhyB-Pif3 strains and constructs; Prof. Yuhei Goto for sharing iRFP budding yeast strains; Prof. Mathieu Coppey for sharing a PhyB-PIF6 mammalian cell line; Prof. Kazuhiro Aoki for sharing plasmids; Prof. Wilfried Weber and Prof. Gerald Radziwill for sharing plasmids for transcriptional PhyB-PIF6 control; Prof. José L. Avalos and Dr. Evan Zhao for providing the *EL222* plasmid; Prof. Andrew Murray for providing the *pol3-L523D* strain; and Dr. Enrico Tenaglia for help with molecular cloning. We thank the mechanical and electronics workshops of the Institute of Physics in the Cubotron building at EPFL for help building the hardware for optogenetic stimulation. We acknowledge support for VG, ML, LS, and SJR provided by SNSF grants CRSK-3_190526, 310030_204938, and CRSK-3_221036 awarded to SJR. LS is also supported by the EPFLglobaLeaders doctoral fellowship. A.M. acknowledges support through grant DFG MO2192/10-1 by the Deutsche Forschungsgemeinschaft.

## Author contributions

VG, ML, LS, MB, AM, NM, and SJR designed the experiments and analyzed the data. VG constructed the strains used for evolution. VG performed the directed evolution experiments. VG and ML characterized the evolved variants, including reporter strain construction. LS constructed and tested the multiplexing system, aided by VG. VG, MB, and AM purified proteins and performed *in vitro* biophysical characterization. VG, AM, and SJR wrote the manuscript. NM supervised the experiments in mammalian cells. AM supervised the *in vitro* experiments. SJR supervised the overall project and acquired funding.

## Competing interests

VG and SJR have filed a provisional patent application for the method reported in this manuscript through EPFL.

## Supplementary Material

### Supplementary Note 1: List of previous multiple-goal continuous directed evolution campaigns

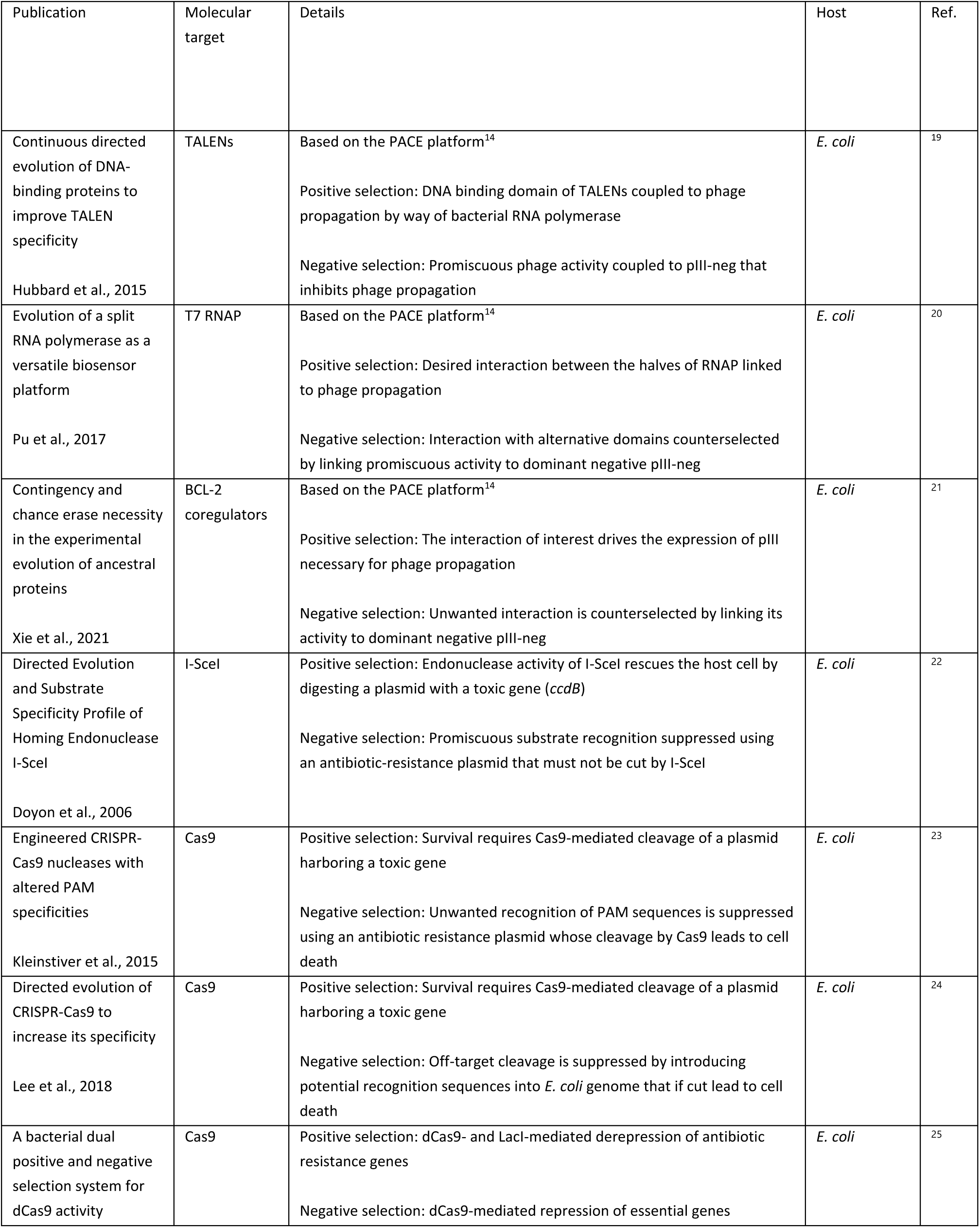
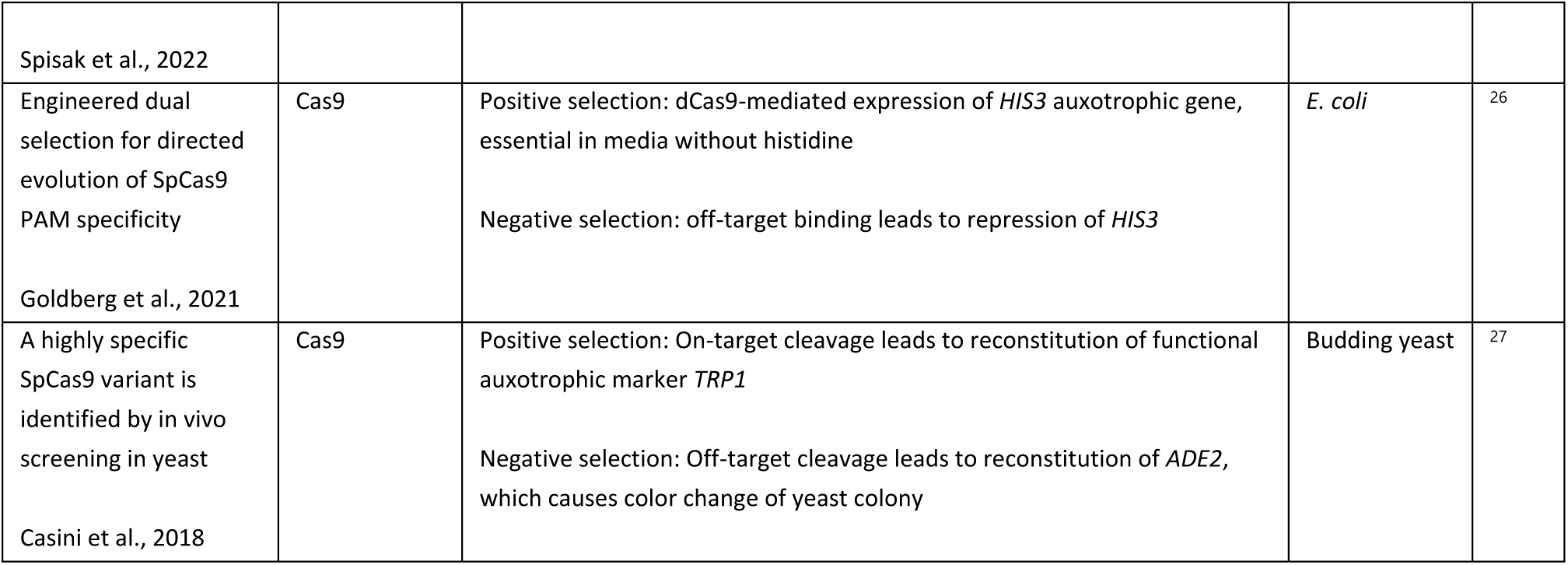

### Supplementary Note 2: List of previous selection/counterselection directed evolution campaigns

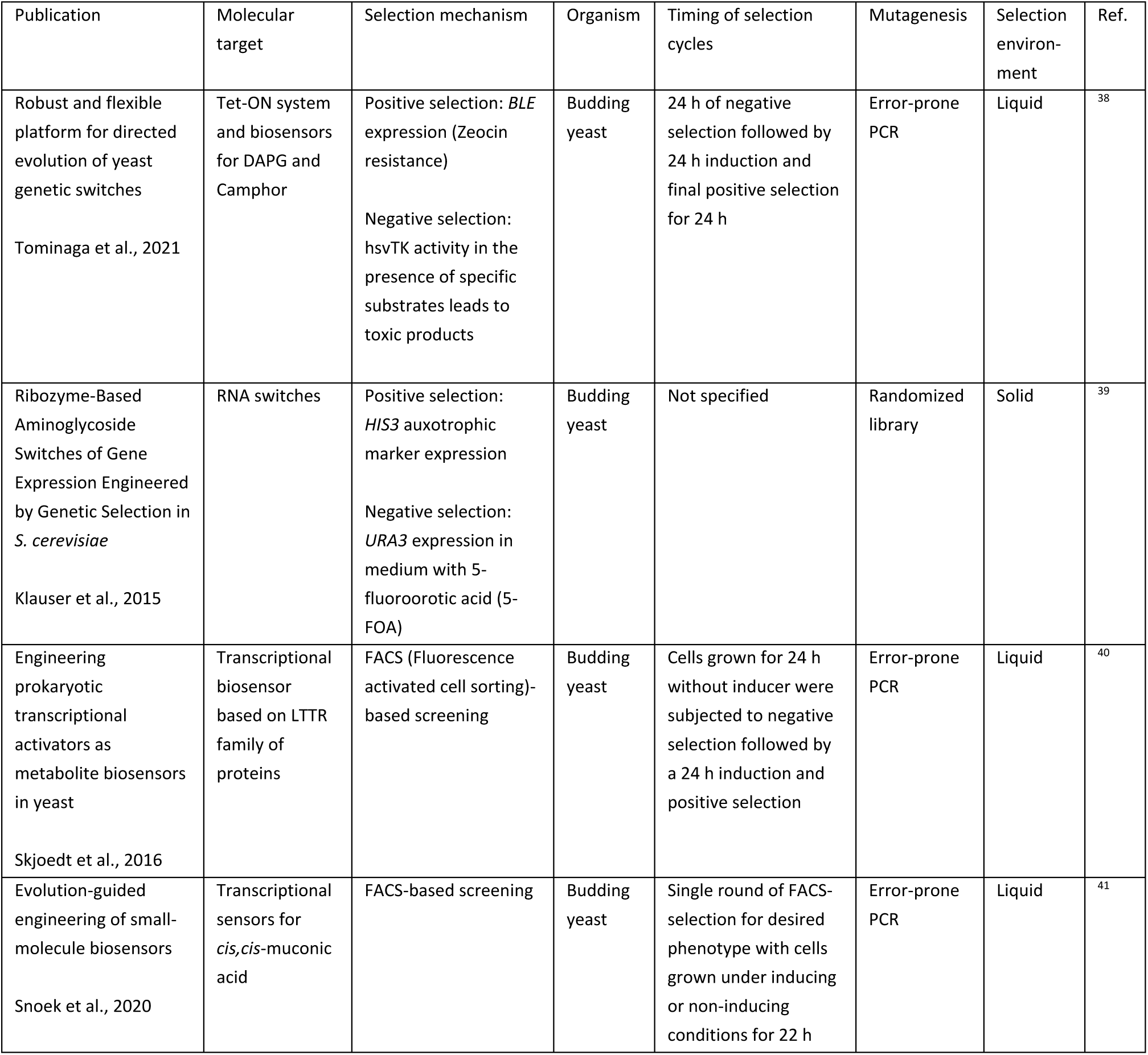
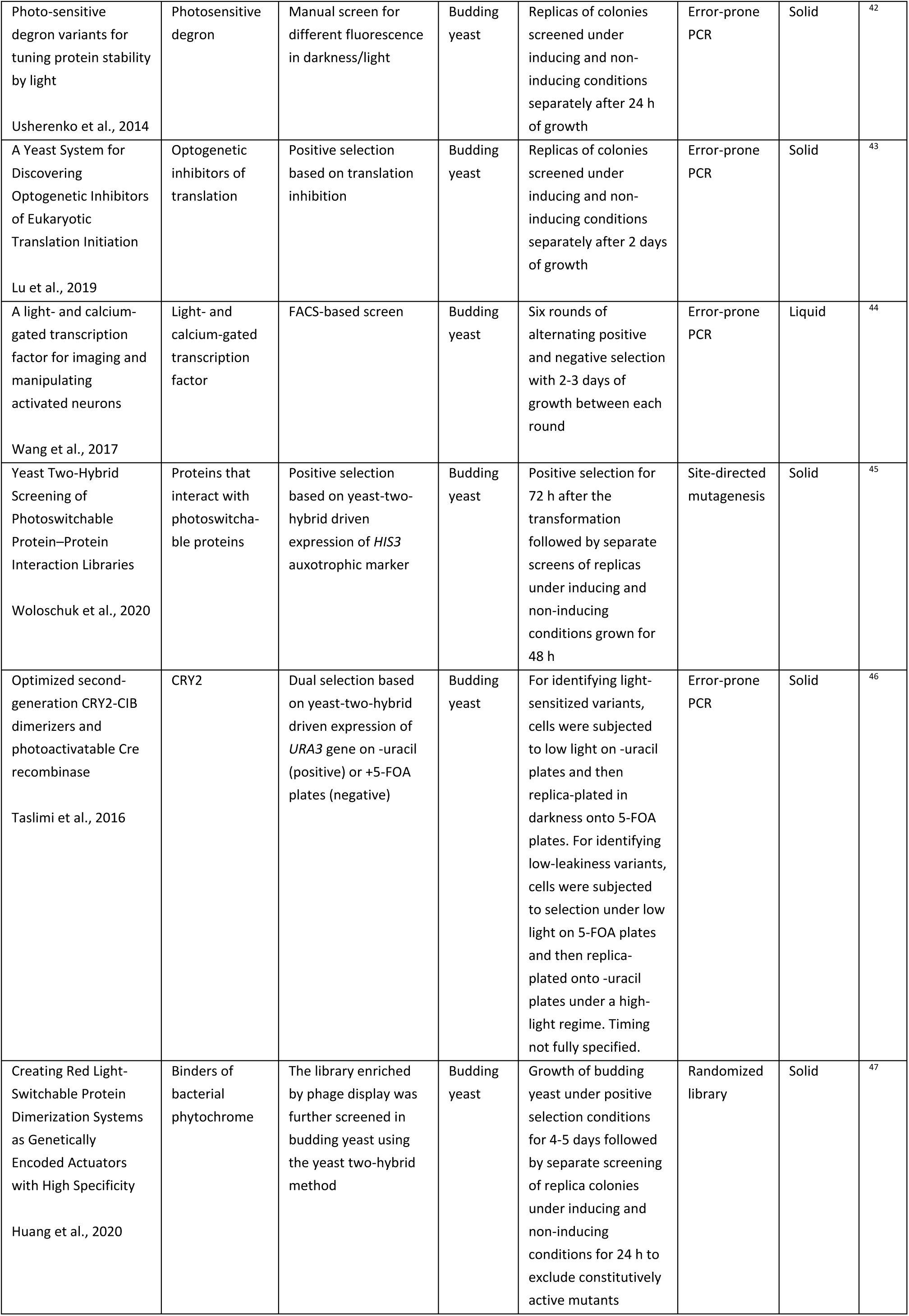
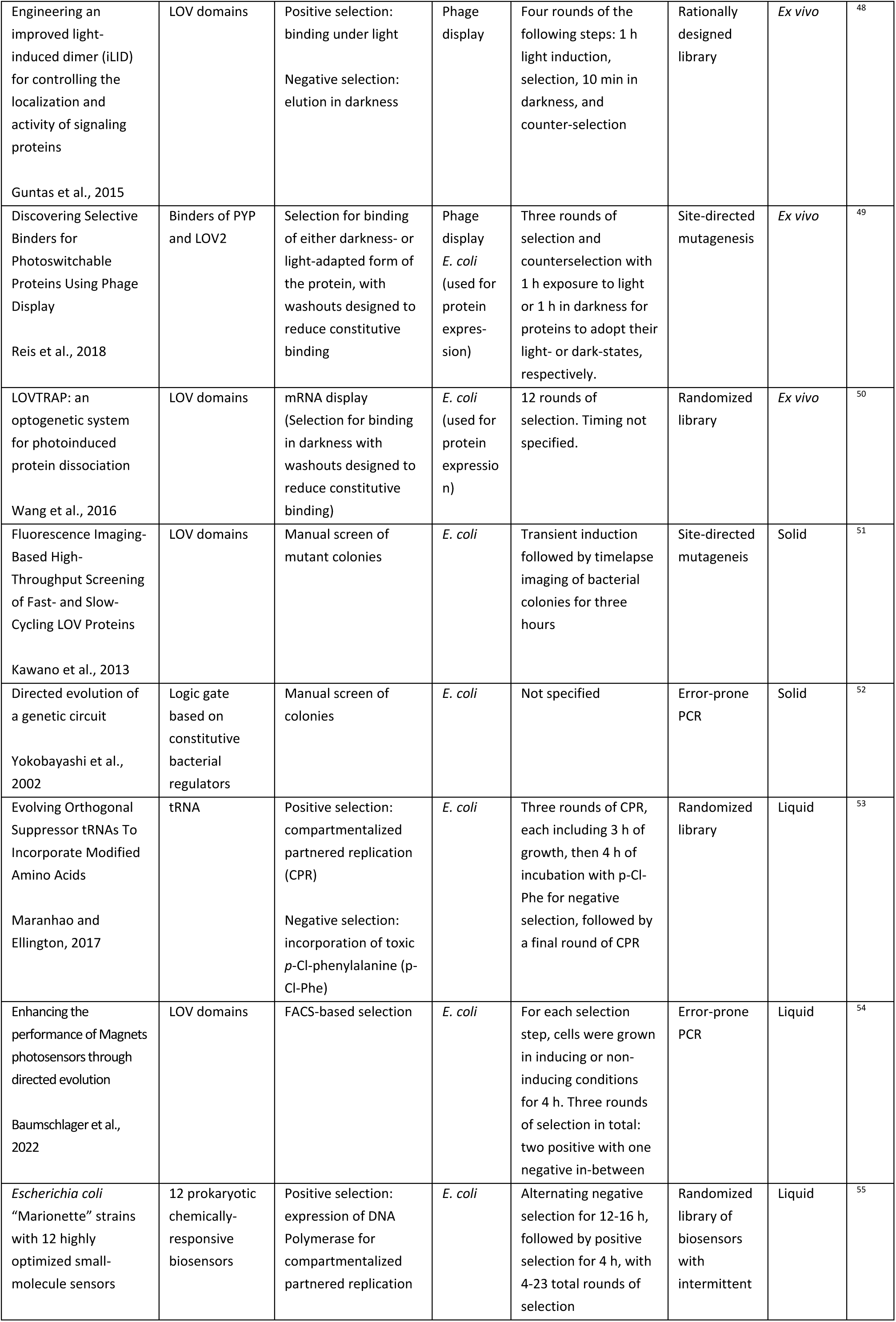
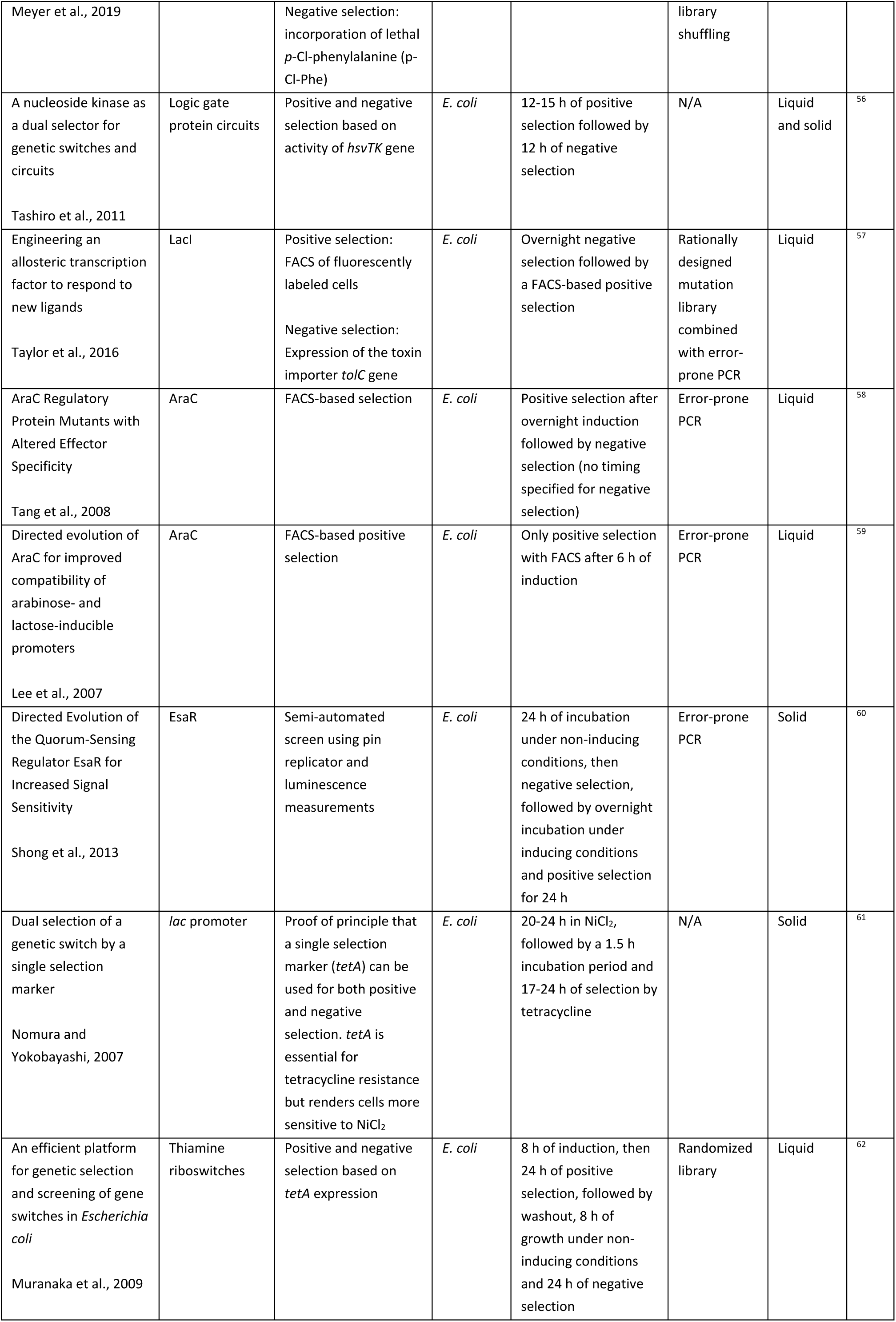
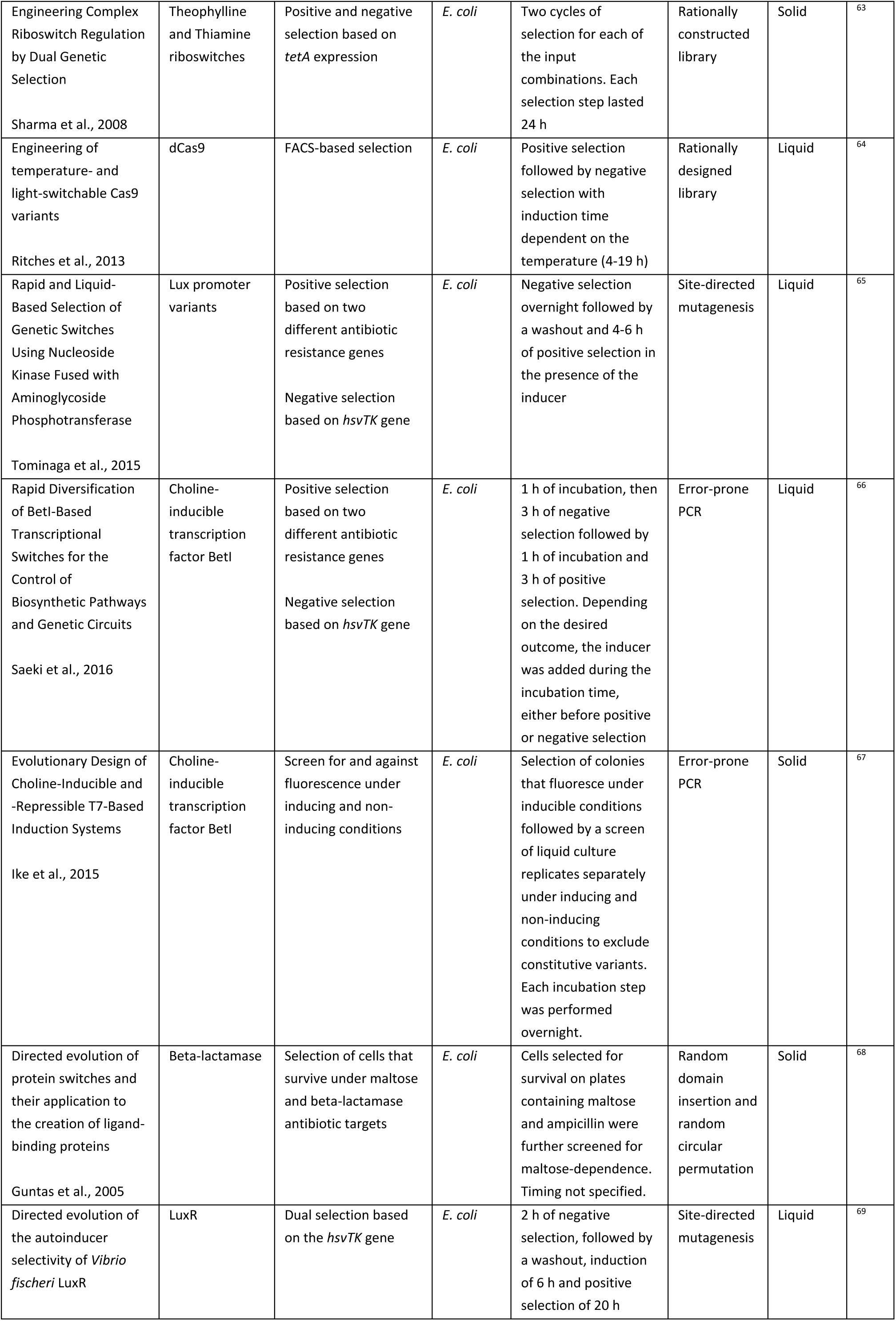
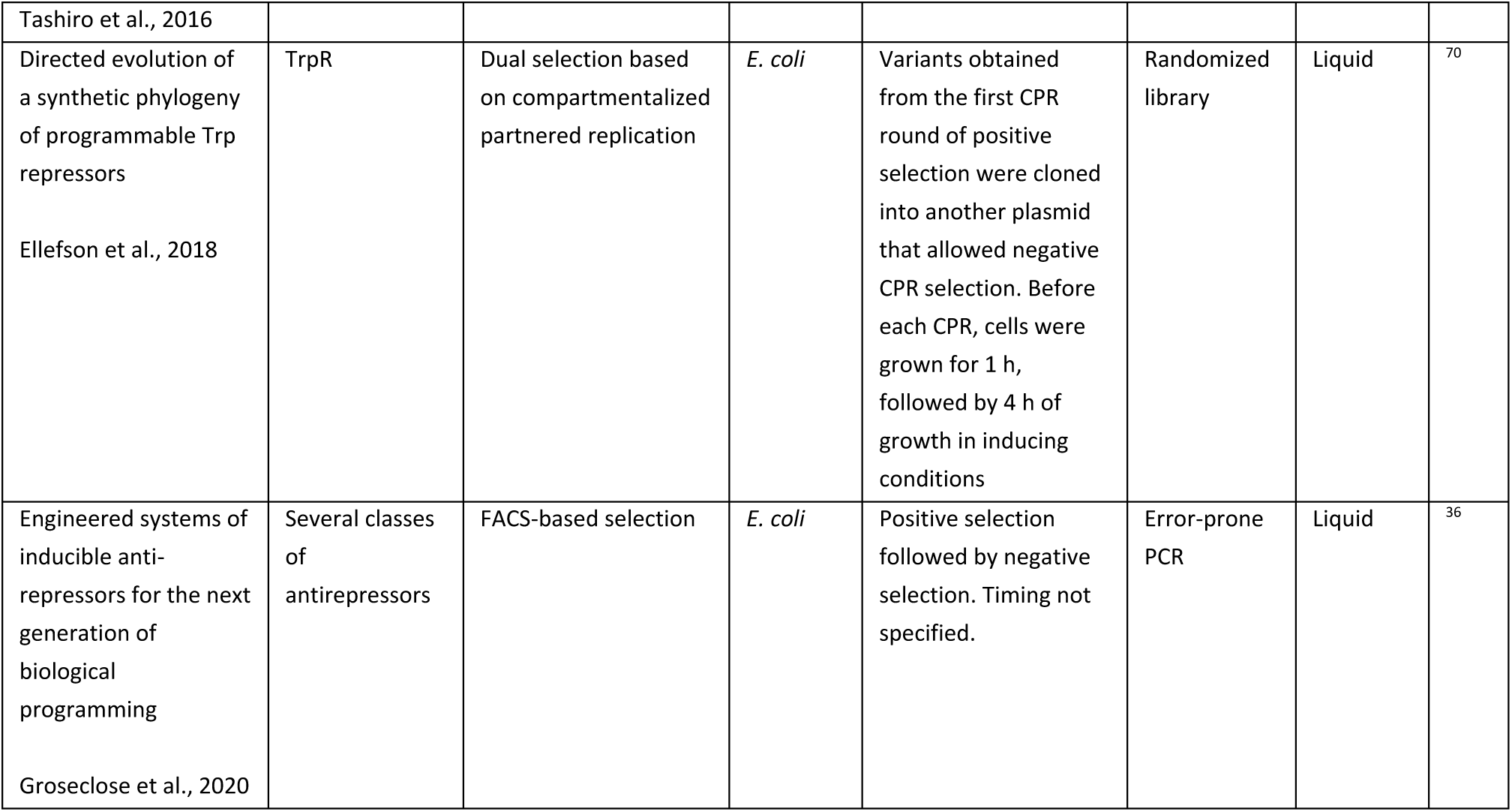

### Supplementary Note 3: El222 optogenetic system

El222 is a transcription factor from marine bacterium *Erythrobacter litoralis*.^140^ El222 has been adapted for use in eukaryotes by way of a tandem fusion of the prokaryotic El222 transcription factor, a nuclear localization signal, and the VP16 transcriptional activation domain. In darkness, the interaction of the helix-turn-helix (HTH) DNA binding domain with DNA is sterically hindered by the light-sensing LOV-domain^139^. Upon blue light illumination, a covalent bond is formed between the Cys-75 residue and the flavin chromophore^191^. (All references to mutant residue numbers in El222 are given with respect to bacterial El222 to match the numbering used in previous work^88,139,140^.) This leads to a conformational change, which dislocates the Jα helix of the LOV domain. Subsequently, the protein forms a homodimer and binds to the DNA substrate using its exposed HTH domain. In the absence of blue light, the protein reverts to its ground state. For brevity, we refer to the promoter consisting of 5 El222 binding sites^192^ upstream of a minimal eukaryotic promoter as *LIP*.

### Supplementary Note 4: PhyB-Pif3 optogenetic system

Phytochromes are photosensors in plants, bacteria, and fungi^193^. Phytochrome B and its interaction partner Pif3 are used by plants to adapt gene expression to environmental light conditions^194^.

The light-inducible PhyB-Pif3 interaction has been extensively used in organisms other than plants^106–109^. By fusing one protein to PhyB and another to Pif3, one can induce protein-protein interactions in a temporarily and spatially precise manner, for example, in order to sequester proteins to specific cellular compartments^107^. In the yeast two-hybrid version of the system, it is used for light-inducible transcriptional control in budding yeast.^108,195^ Typically, PhyB and Pif3 are fused to the Gal4 DNA binding domain and the Gal4 transcriptional activation domain, respectively.^108^ When bound to its chromophore and activated by red light (≈650 nm), PhyB binds Pif3, thereby, bringing the transcriptional activation and DNA binding domains close together, which leads to the expression of the *GAL* family of genes. In the presence of far-red light (≈ 740 nm) or prolonged darkness, PhyB changes conformation again and dissociates from Pif3.

The photoreceptive core of the PhyB protein contains a linear tetrapyrrole chromophore.^196^ In plants, this chromophore is phytochromobilin. However, phytochromobilin is not produced in many model organisms. Thus, to use PhyB-Pif3, one adds phycocyanobilin (PCB), which is the chromophore in algal phytochromes, to the incubation medium prior to optogenetic experiments. It has been shown that bacterial and fungal phytochromes can carry biliverdin, another tetrapyrrole molecule, as a chromophore.^196^

### Supplementary Note 5: Tetracycline-regulated gene expression systems

Tetracycline-responsive systems, originally identified in bacteria as part of a feedback loop that mediates resistance to the antibiotic tetracycline, are widely used for controlling transcription. The system is made up of the tTA regulator, which binds the *tetO* DNA sequence. In its original form, tTA is part of the Tet-Off system that is inhibited by tetracycline or the closely related molecule doxycycline^197^. Mutations that reverse tTA activity with respect to the inducer have been used to construct rtTA.^115^ To function as a transcriptional activator in eukaryotes, rtTA is typically fused to the VP16 transcriptional activation domain. To construct an inducible eukaryotic promoter, the binding sites for rtTA, *tetO*, are placed next to a minimal promoter. However, this system, called the Tet-On system, exhibits high basal activity in the absence of the inducer. To circumvent this, we fused the PEST degron to rtTA, which leads to increased degradation of rtTA.

### Supplementary Note 6: Summary of cell cycle biology relevant to optovolution

#### Wild-type cell cycle control

In this note, we briefly summarize the wild-type model of cell cycle control in budding yeast.^130^ In the next section, we focus on how the cell cycle runs in the Dajbog strains.

At the biochemical level, the rhythm of the cell-cycle oscillator is predominantly generated by a negative feedback loop^126^. In this loop, cyclin-dependent kinase (CDK) and the anaphase-promoting complex/cyclosome (APC/C) alternate in activity and drive cell cycle progression.

In budding yeast, there is a single cyclin-dependent kinase, Cdk1 (also known as Cdc28). Processes that Cdk1 controls include the transcription of hundreds of cell cycle-periodic genes, DNA replication, spindle pole body duplication, and spindle formation. Activity and specificity of Cdk1 towards different substrates is mainly regulated through its interaction with regulatory proteins called cyclins. Different cyclins generally promote distinct cell-cycle events and are expressed at different stages of the cell cycle. In *S. cerevisiae*, cyclins are commonly classified into four groups: G1, G1/S, S, and M cyclins. Each cyclin-Cdk complex promotes the activation of the next group of cyclins.

The G1 cyclin Cln3 activates G1/S cyclins Cln1 and Cln2 and has a role in cell-cycle entry in response to growth. Cln1-3 are responsible for the initiation of formation of the daughter cell and expression of the 100s of genes of the Start regulon. At the onset of S phase, levels of S cyclins Clb5 and Clb6 rise. Their association with Cdk1 is primarily responsible for the initiation of DNA replication by triggering the activation of origins of replication. Approaching mitosis, the concentrations of the remaining B-type cyclins increase. In budding yeast, there are four such cyclins, Clb1, Clb2, Clb3, and Clb4. Their association with Cdk1 enables assembly of the mitotic spindle and attachment of chromosomes to the spindle.

In order to enable mitotic exit and reset the state of the cell for new cell cycles, cyclins have to be inactivated. While the activity G1/S cyclins is brought down by degradation and inhibition of the transcription factor SBF, S and M phase cyclins are not degraded until anaphase. At least two different mechanisms help repress Clb cyclin activity. The first one is through the activation of APC/C by the proteins Cdc20 and Cdh1. APC/C is a ubiquitin ligase complex that marks cyclins for degradation by the proteasome. The second mechanism is through the protein Sic1, which stoichiometrically inhibits Cdk1-Clb complexes, thereby effectively reducing their activity.

#### Rewired cell cycle control in the Dajbog strains

Although cyclins are specialized for certain phases of the cell cycle, the functions of some cyclins overlap. For example, when overexpressed, S phase cyclin Clb5 can initiate budding in a *cln1-3Δ* block^127^. Similarly, the M-phase cyclins Clb1^198^or Clb2^123^can assume the roles of all other Clb cyclins in a *clb1-6Δ* background. Endogenous *CLB3-6* genes also suffice for cell cycle start when the Clb-cyclin inhibitor *SIC1* is deleted, rendering *cln1-3Δ sic1Δ* cells viable^133^.

While G1/S cyclin and S cyclin expression initiates at about the same time under the control of transcription factors Sbf and Mbf^130^, respectively, S cyclins are inactive until G1/S cyclins have sufficiently degraded Sic1 and Cdh1. Thus, in wild-type cells, although *CLN2* and *CLB5* are simultaneously transcriptionally activated, there is a delay between G1/S-Cdk and S-Cdk activity^130,199^. We expect this delay to persist in Dajbog1 cells. Deleting *SIC1* in Dajbog2 cells is expected to cause an increase in chromosomal rearrangements due to premature S-Cdk activity^200^ but this did not noticeably interfere with our directed evolution experiments.

Clb5-Cdk1 leads to the firing of origins of replication in S phase. Overexpression of destruction box deficient Clb5db in galactose medium arrests the cell cycle by inhibiting the formation of pre-replicative origins, a process known as origin of replication relicensing^201–203^. This represents the expected mechanism for the lethality of *CLB5* overexpression in glucose medium. This observation led us to further sensitize Dajbog1 cells to continuous Clb5 activity by deleting *SIC1*, resulting in Dajbog2 strains.

## Supplementary Figures

**Supplementary Figure 1.**
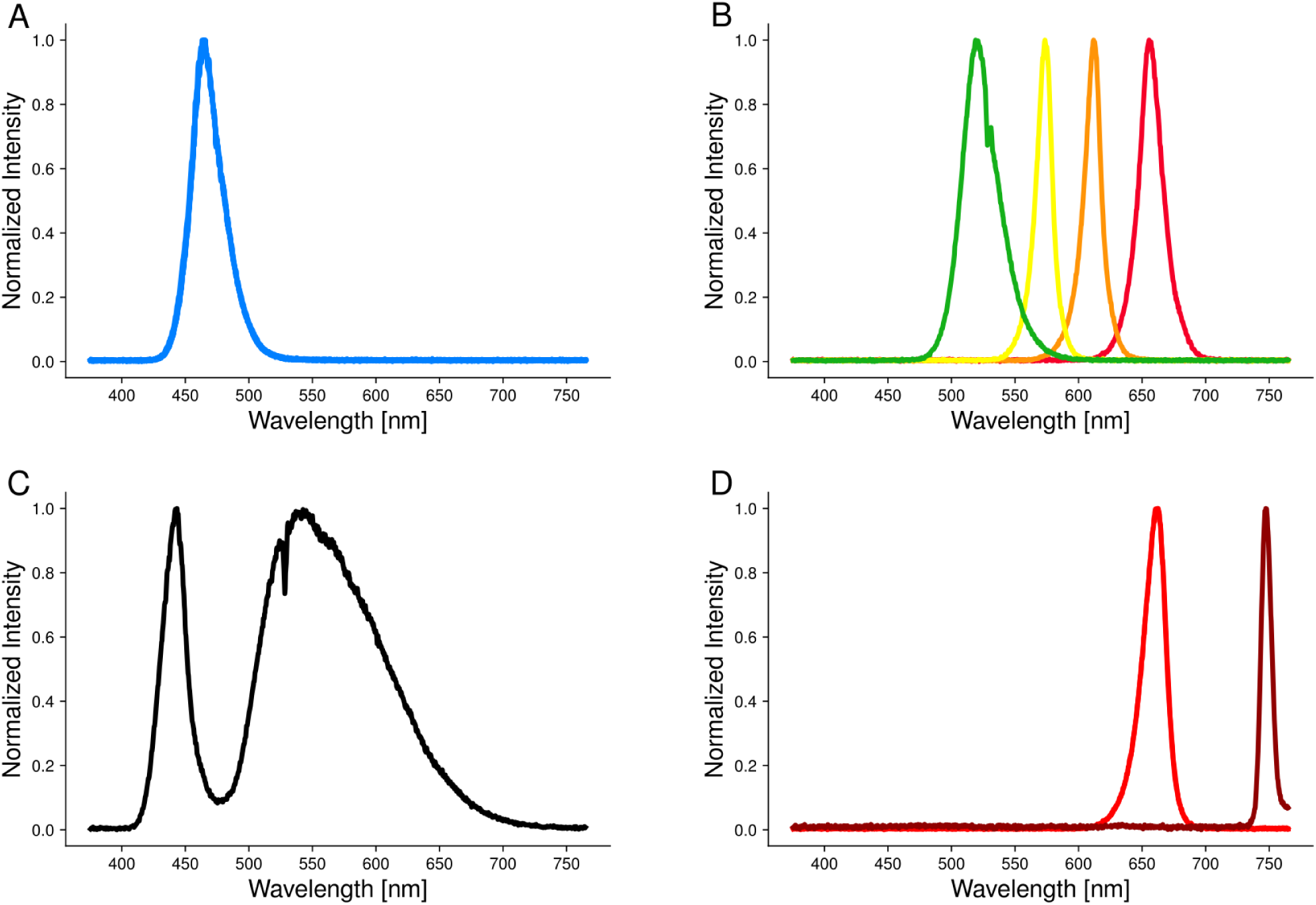
Emission spectra of light sources used. A. Blue LED panel used for evolving and inducing El222 and its variants (light intensity 8.3 mW/cm^2^). B. LEDs used to evolve *in vivo* green responsive El222 variants from left to right: 24 mW/cm^2^, 0.12 mW/cm^2^, 2.6 mW/cm^2^, 0.46 mW/cm^2^. C. Diascopic LED light used for inducing El222 and its variants for microscopy. The intensity of the light is 8.4 mW/cm^2^ when set to 20% of the maximal power and 97 mW/cm^2^ when set to 80% of maximal power. D. Red and far-red LEDs used for switching the PhyB-Pif3 system. Light intensity of the red LEDs is 36 mW/cm^2^, and the light intensity of the far-red LEDs is 26 mW/cm^2^. The reported intensities in panels A, B, and D are total intensities and are for the location of the agar plates.

**Supplementary Figure 2.**
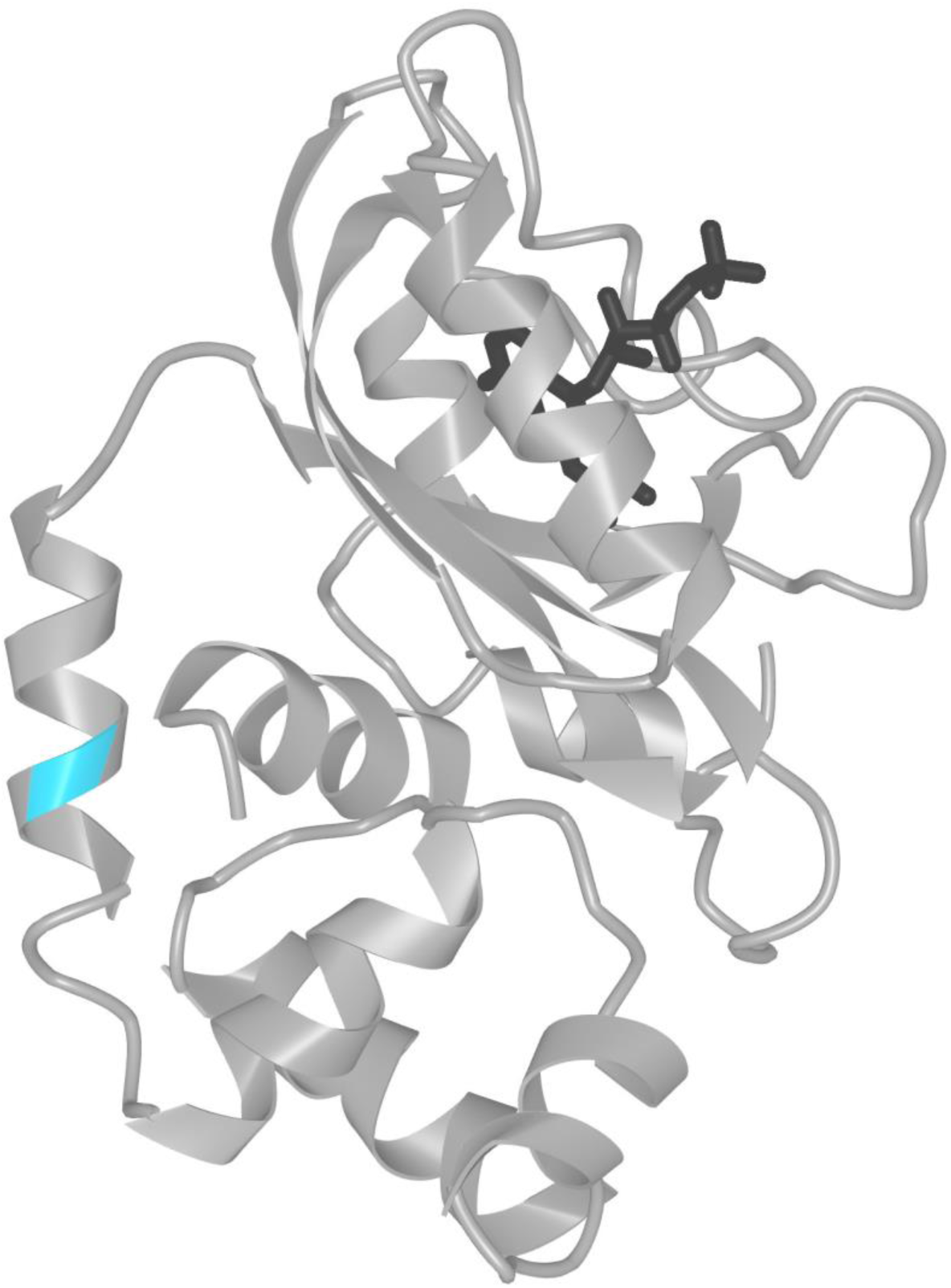
Location of A157 residue marked in teal in the dark state El222 structure (PDB: 3P7N). The chromophore is in black.

**Supplementary Figure 3.**
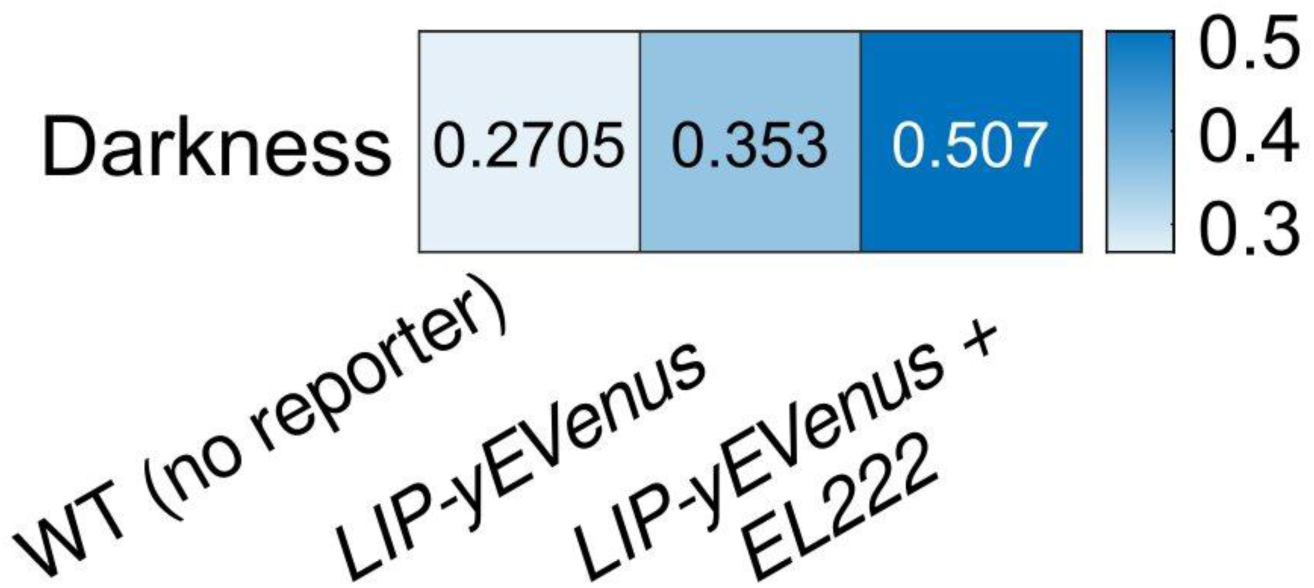
Baseline leakiness of the reporter sets the minimum leakiness attainable in optovolution. Comparison of fluorescence in no-reporter (WT), reporter-only (*LIP-yEVenus*), and reporter plus El222 strains (*LIP-yEVenus + EL222*) in darkness. Cells that contain a transcriptional reporter for El222 (*LIP-yEVenus*) but not the transcription factor (e.g., *PGK1pr-EL222*) showed elevated fluorescence compared to wild-type budding yeast cells. This suggests that other transcription factors might weakly drive the transcription from *LIP* and set the lower bound for leakiness of *LIP*. The leakiness reported for all El222 variants is the difference between the reporter-only and the reporter plus variant strains. Numbers of analyzed cells are given in the Supplementary Table 12.

**Supplementary Figure 4.**
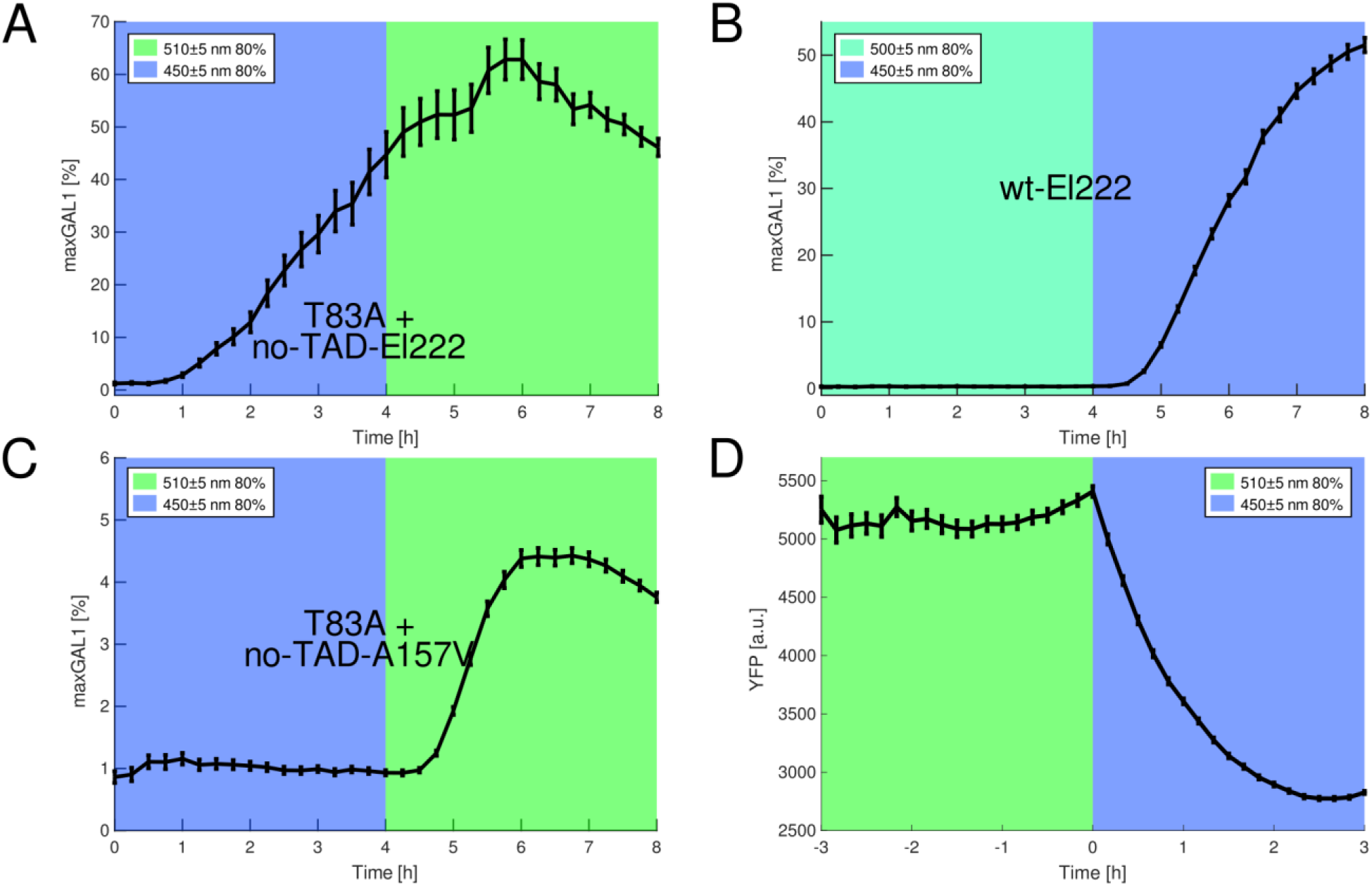
Related to Figure 6. A. no-TAD-El222 does not suffice to suppress El222(T83A). In the same reporter cells, *GAL1*-promoter-driven no-TAD-El222 is expressed in raffinose plus galactose medium but does not clearly inhibit the activity of *PGK1* promoter-driven El222(T83A) in blue 450±5 nm light. B. Wild-type El222 remains inducible in raffinose plus galactose medium in 450±5 nm light only. C. T83A response in the presence of no-TAD-A157V in 510±5 nm and 450±5 nm light. The combination of T83A and no-TAD-A157V produces a smaller response in 510±5 nm light than in 500±5 nm (Fig. 6 R). D. psd does not activate in 510±5 nm light but does activate in 450±5 nm. All error bars indicate the standard error of the mean (SEM).

**Supplementary Figure 5.**
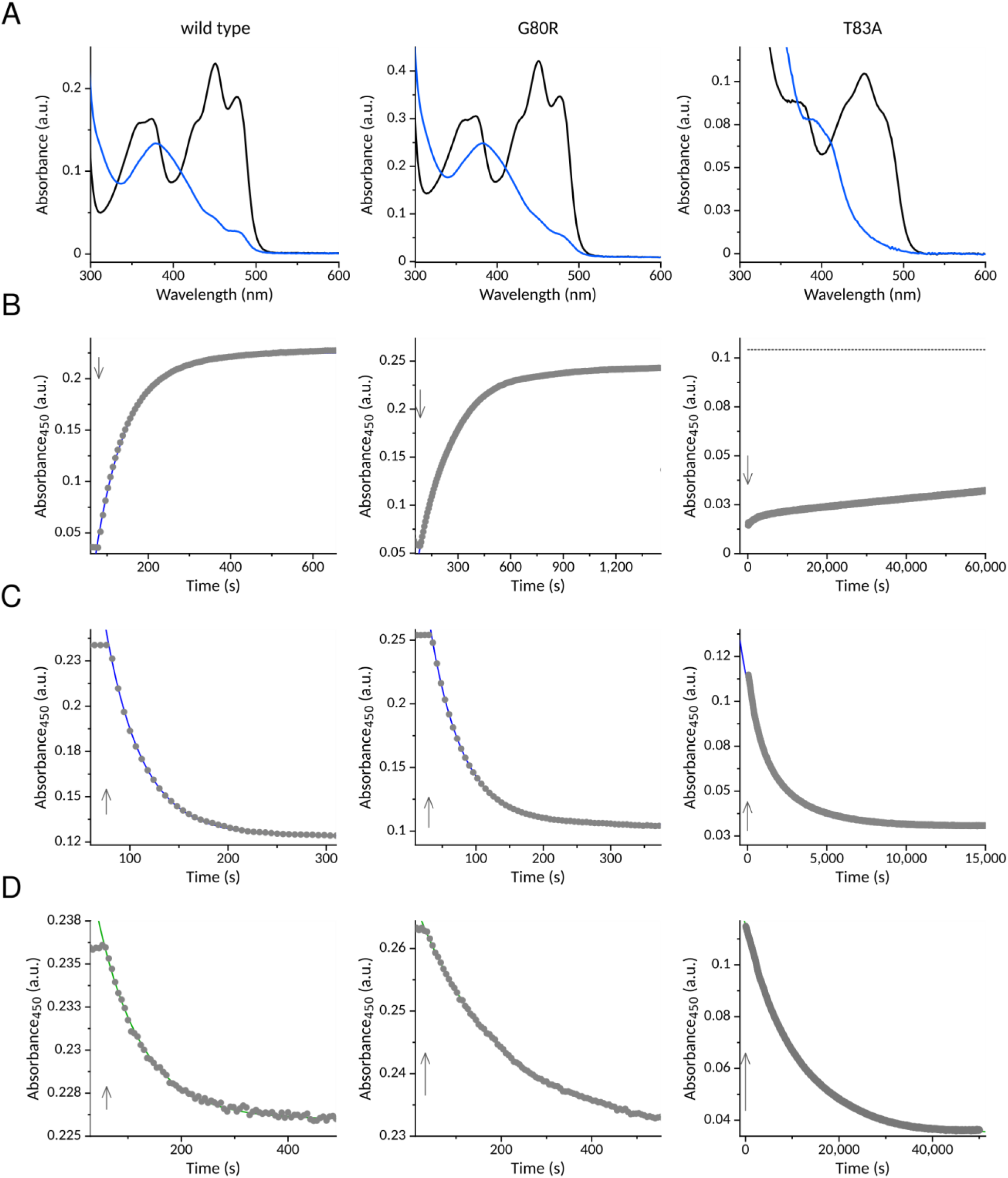
Absorption-spectroscopic characterization of wild-type El222 and the G80R and T83A variants. A. UV/vis absorption spectra of El222 receptors in their dark-adapted states (black lines) and upon exposure to blue light (450 nm, 36 mW cm^-2^) (blue lines). B. The El222 receptors were exposed to saturating 450-nm light to populate the light-adapted state. Upon cessation of illumination (marked by arrow), the thermal recovery to the dark-adapted state was monitored by tracking the absorbance at 450 nm. C. The El222 receptors were irradiated through an interference filter at (450 ± 5) nm at a power of 0.75 mW cm^-2^ while monitoring the photoconversion to the light-adapted state by the absorbance at 450 nm. D. As in panel C but the samples were irradiated through an interference filter at (510 ± 5) nm at a power of 1.5 mW cm^-2^.

**Supplementary Figure 6.**
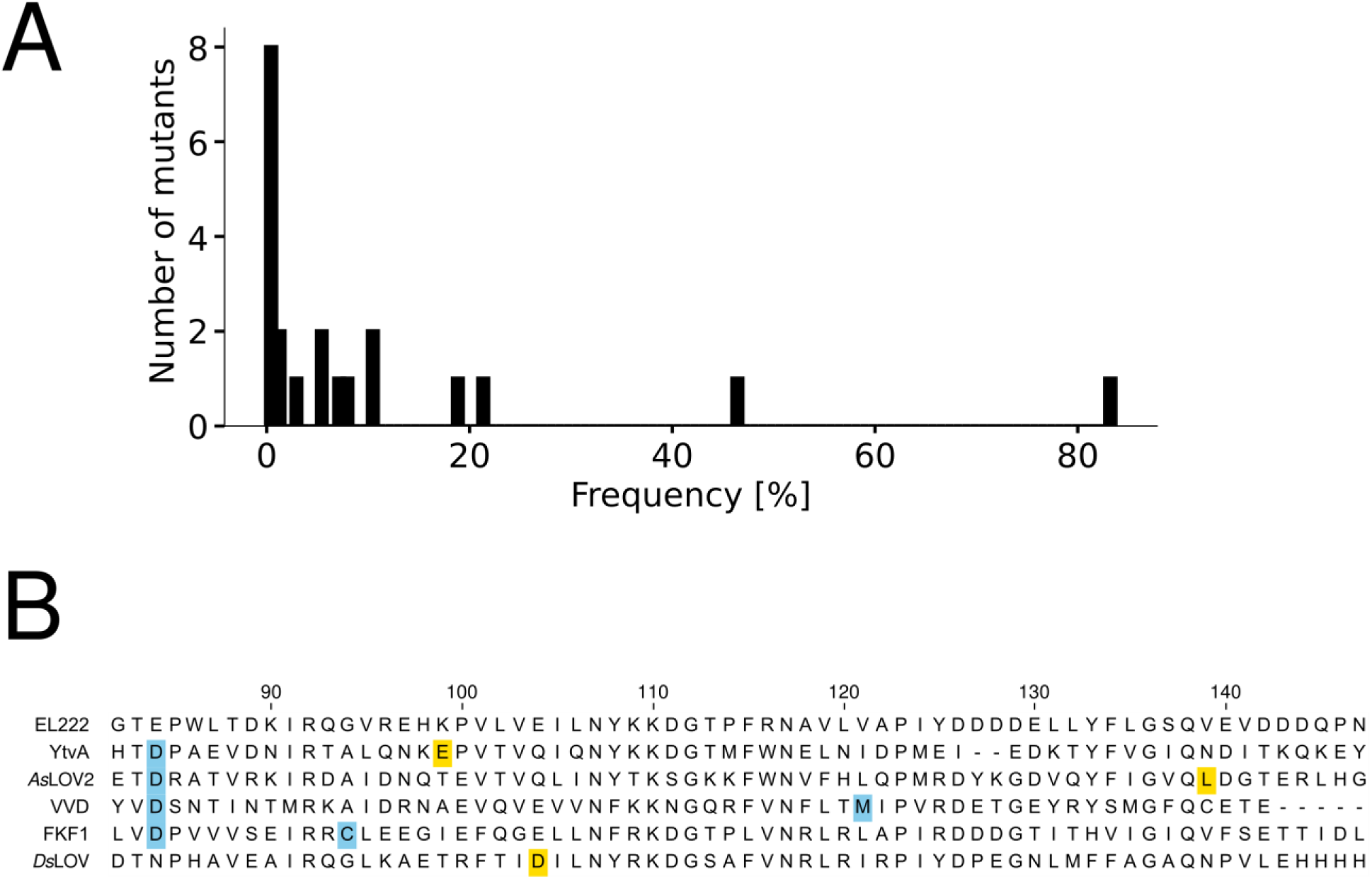
Analysis of El222 mutations found by optovolution in existing LOV domains. A. Histogram of the frequency with which the mutations from our three campaigns are found in putative LOV domains identified in ref.^143^. B. Searching for the mutations from our three campaigns in LOV domains with characterized dark-reversion kinetics^89^ (FKF1 from *Arabidopsis thaliana*, VVD from *Neurospora crassa*, YtvA from *Bacillus subtilis*, *Ds*LOV from *Dinoroseobacter shibae*, and the LOV2 domain of phot1 from *Avena sativa*) identifies the light-sensitizing (blue squares) and low-leakiness mutations (yellow squares) but not the *in vivo* green-responsive mutations.

## Supplementary Tables

**Supplementary Table 1.**
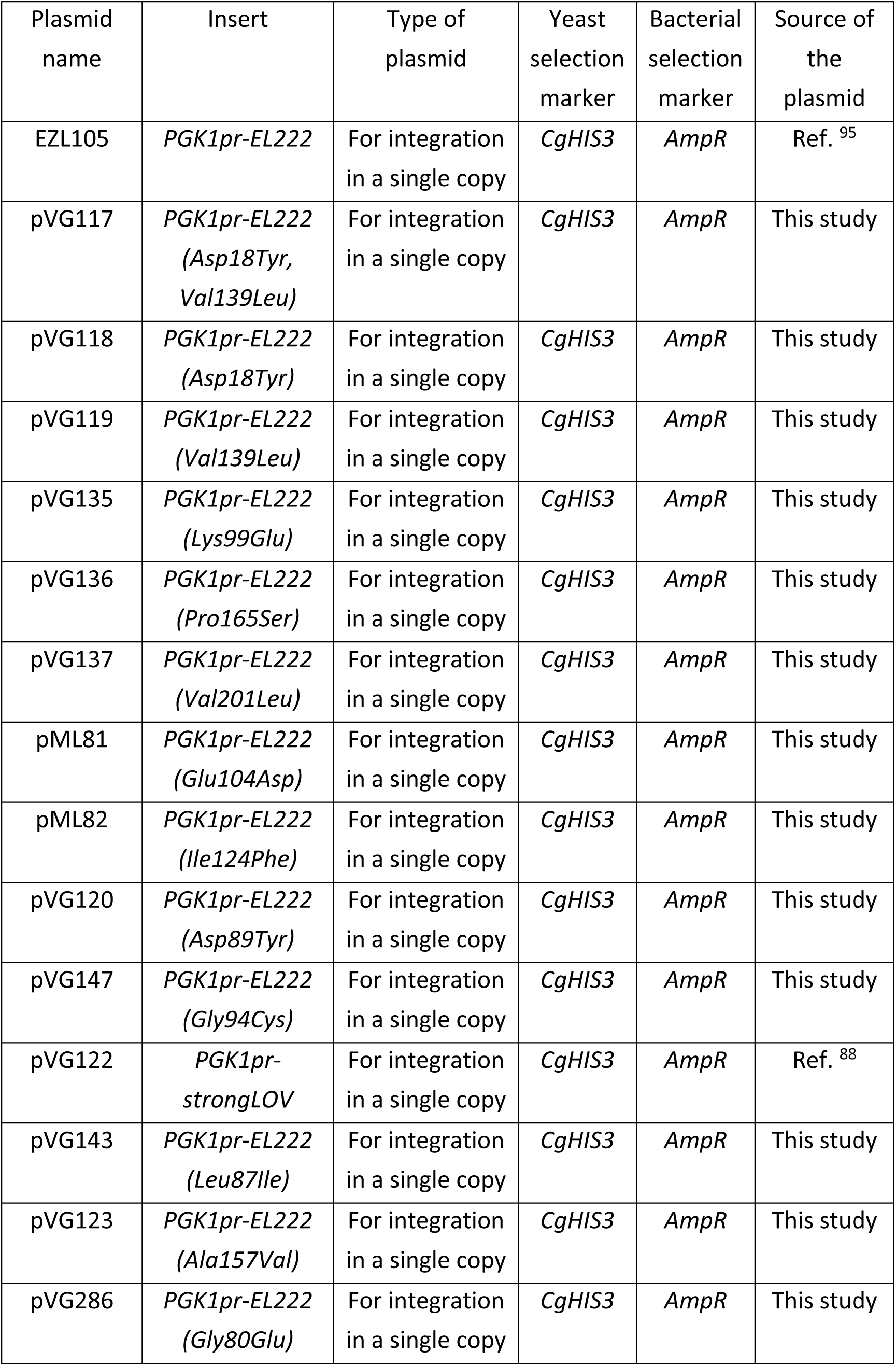
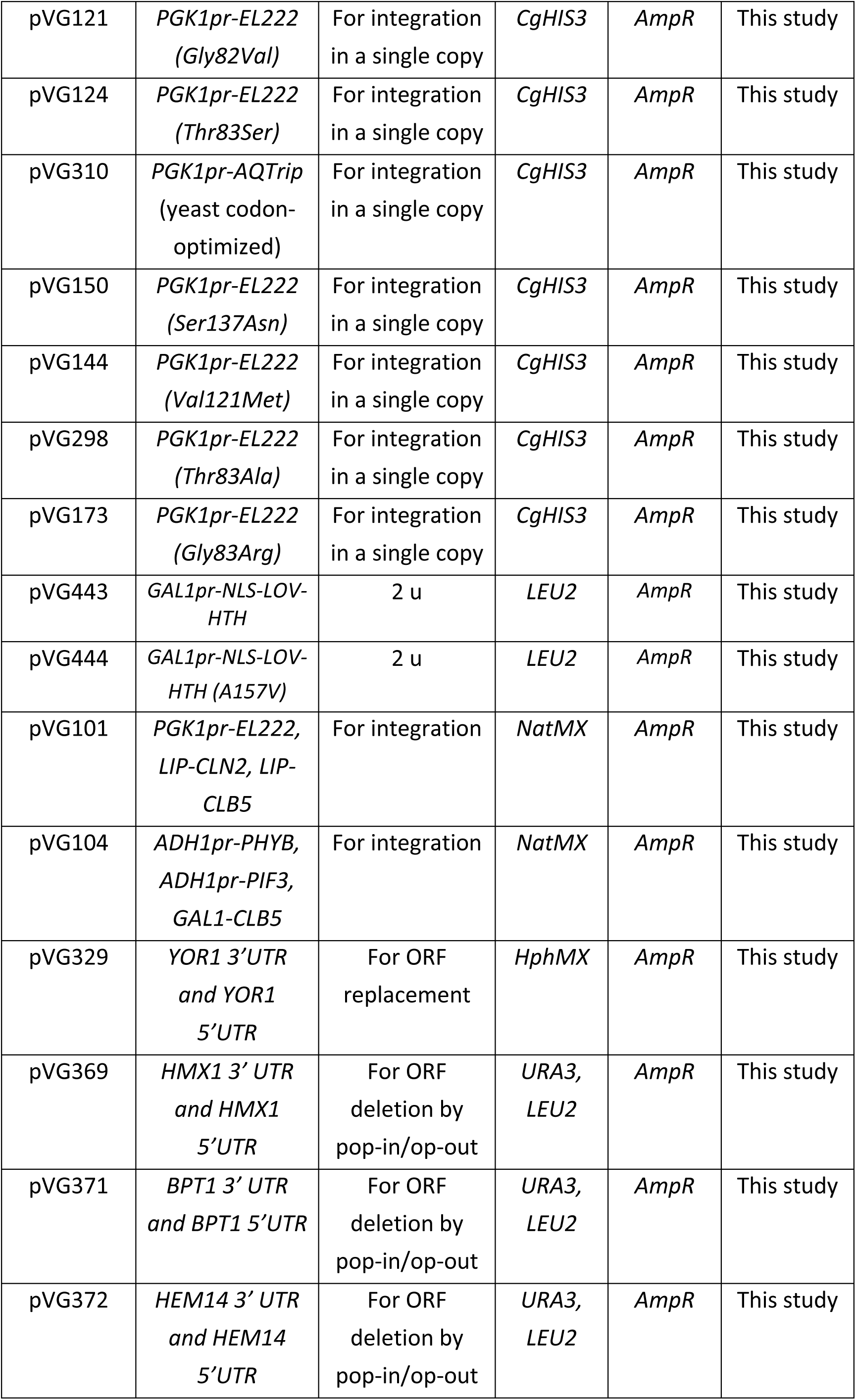
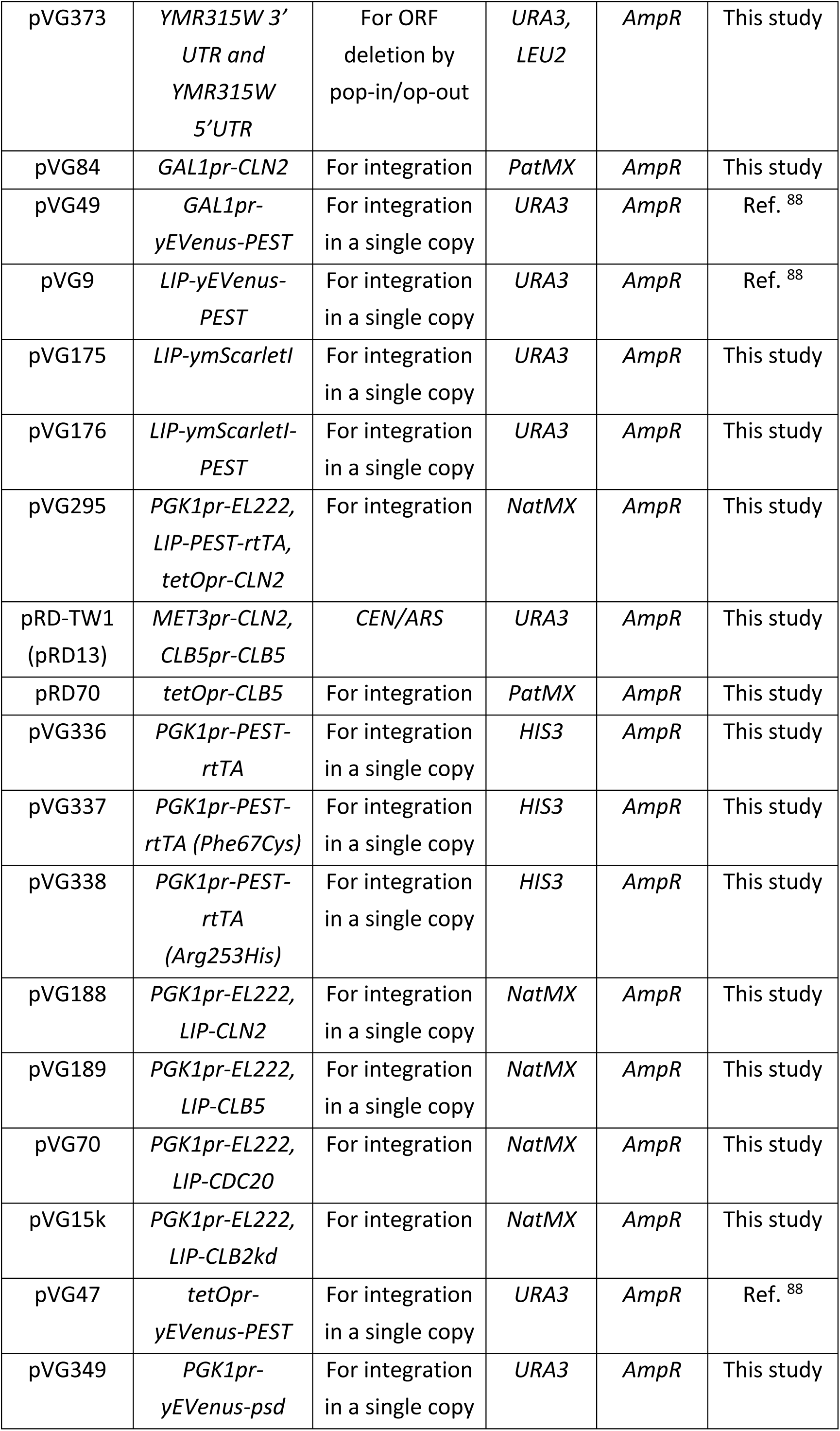
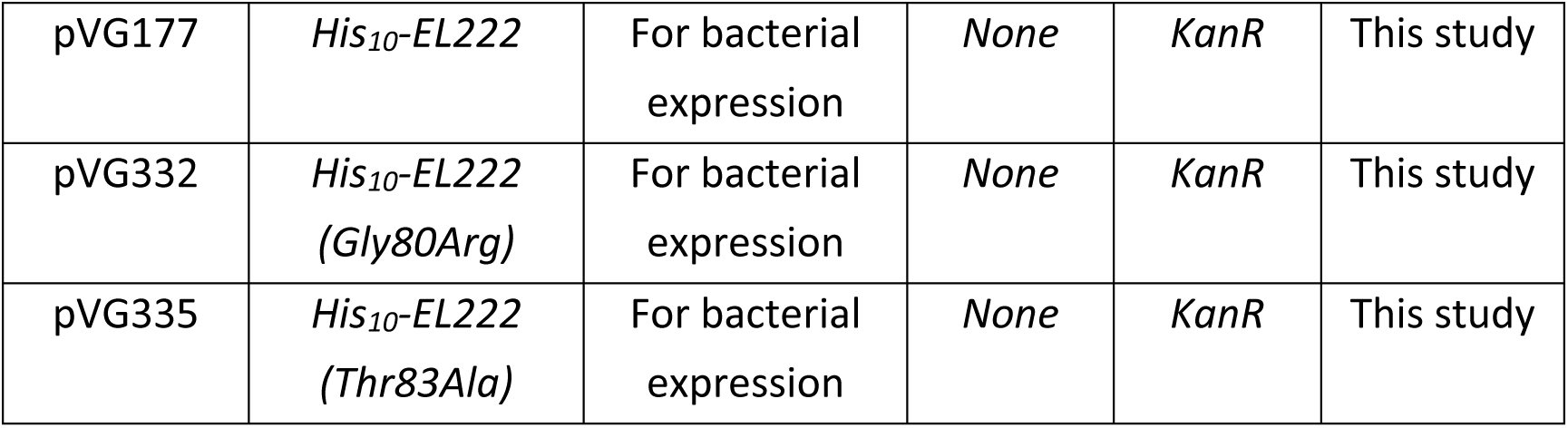
Plasmids used in the study.

**Supplementary Table 2.**
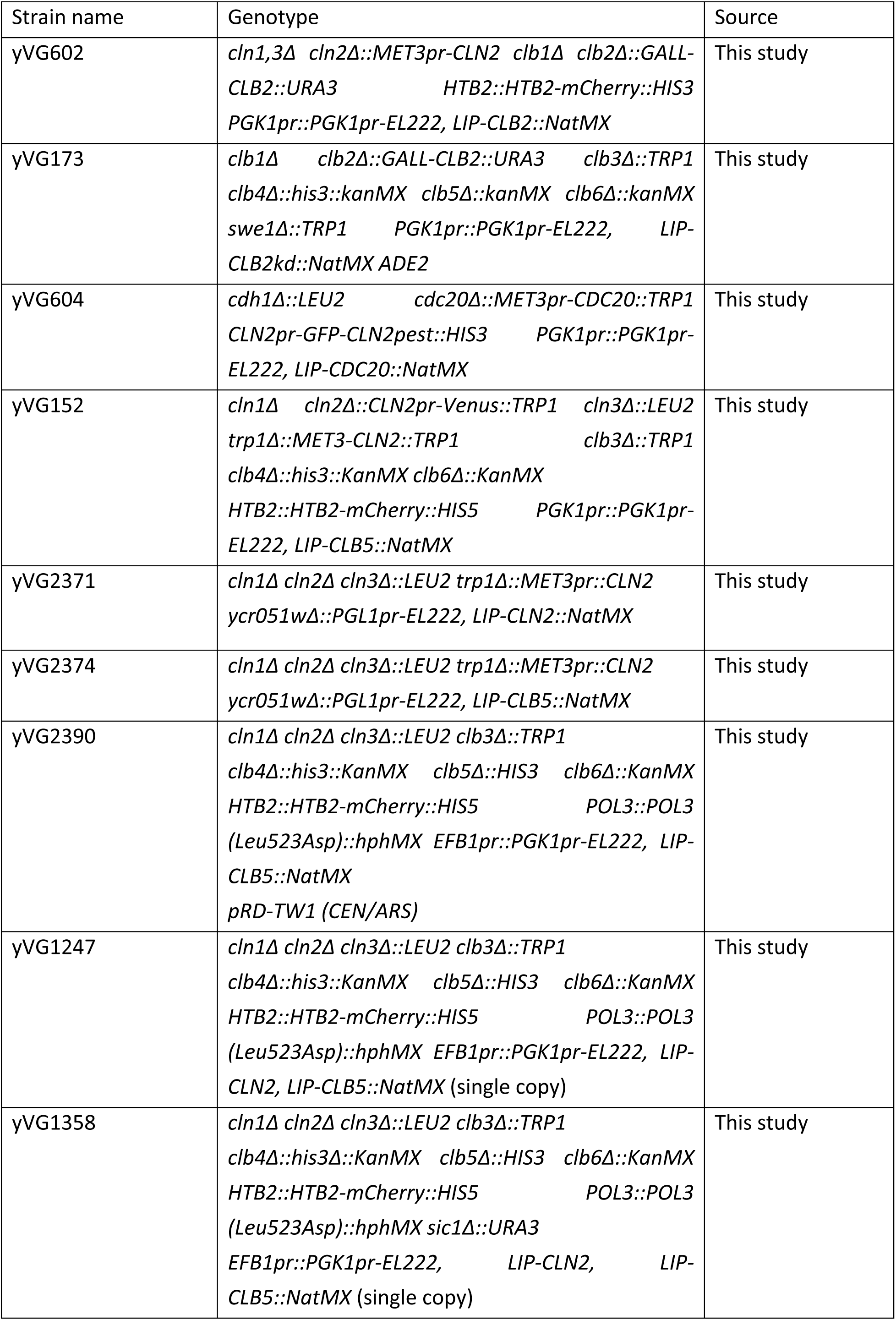
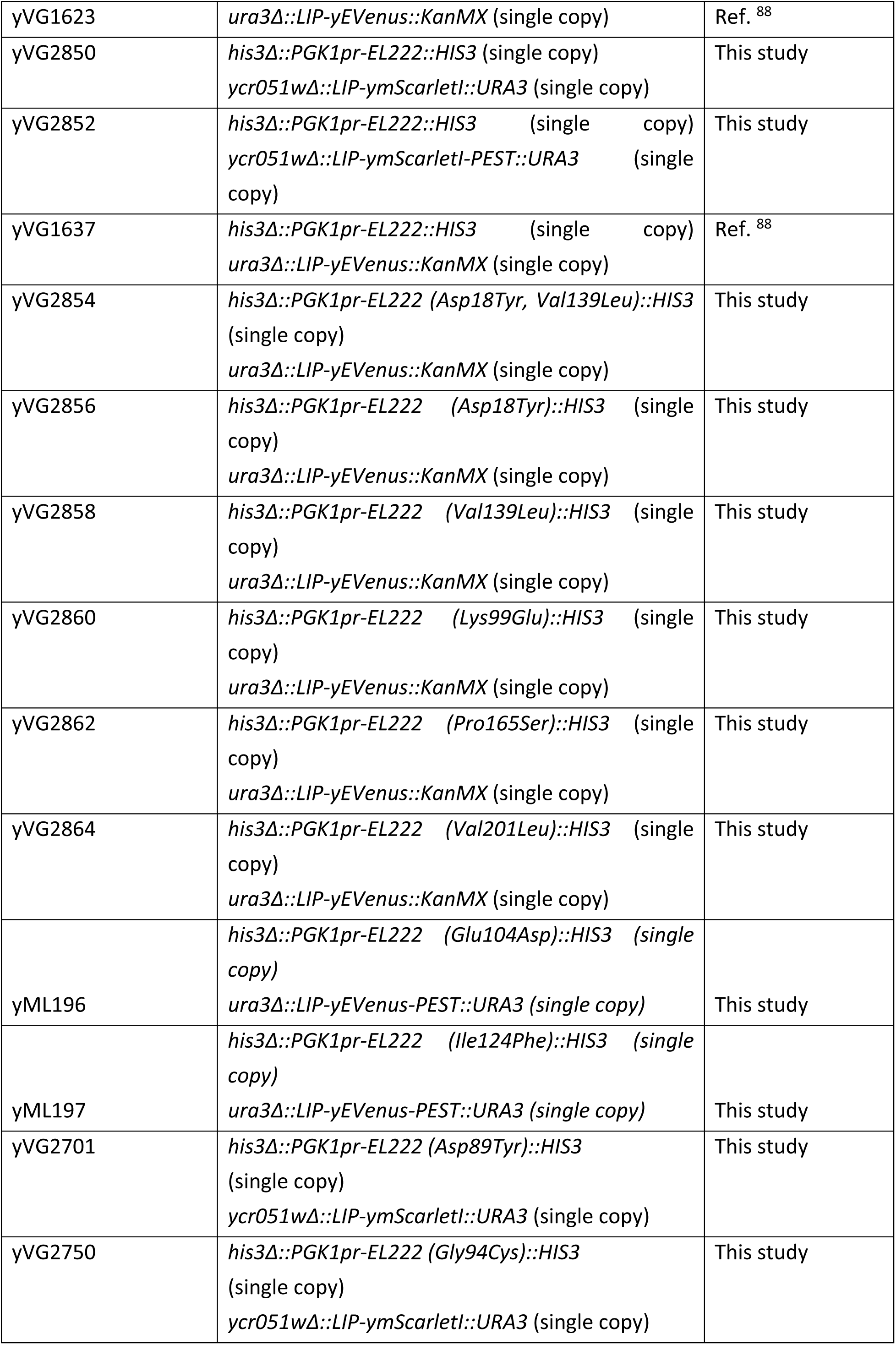
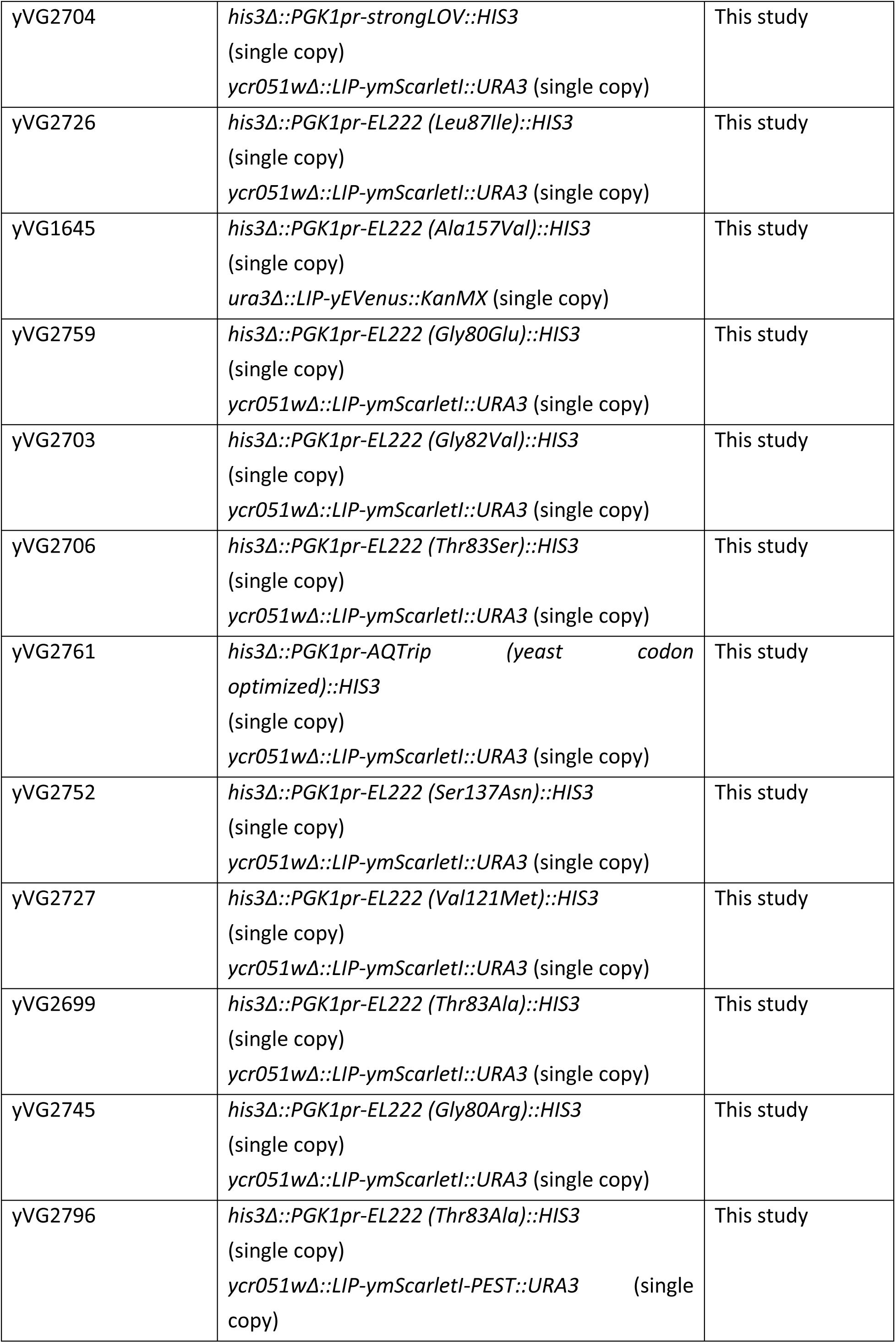
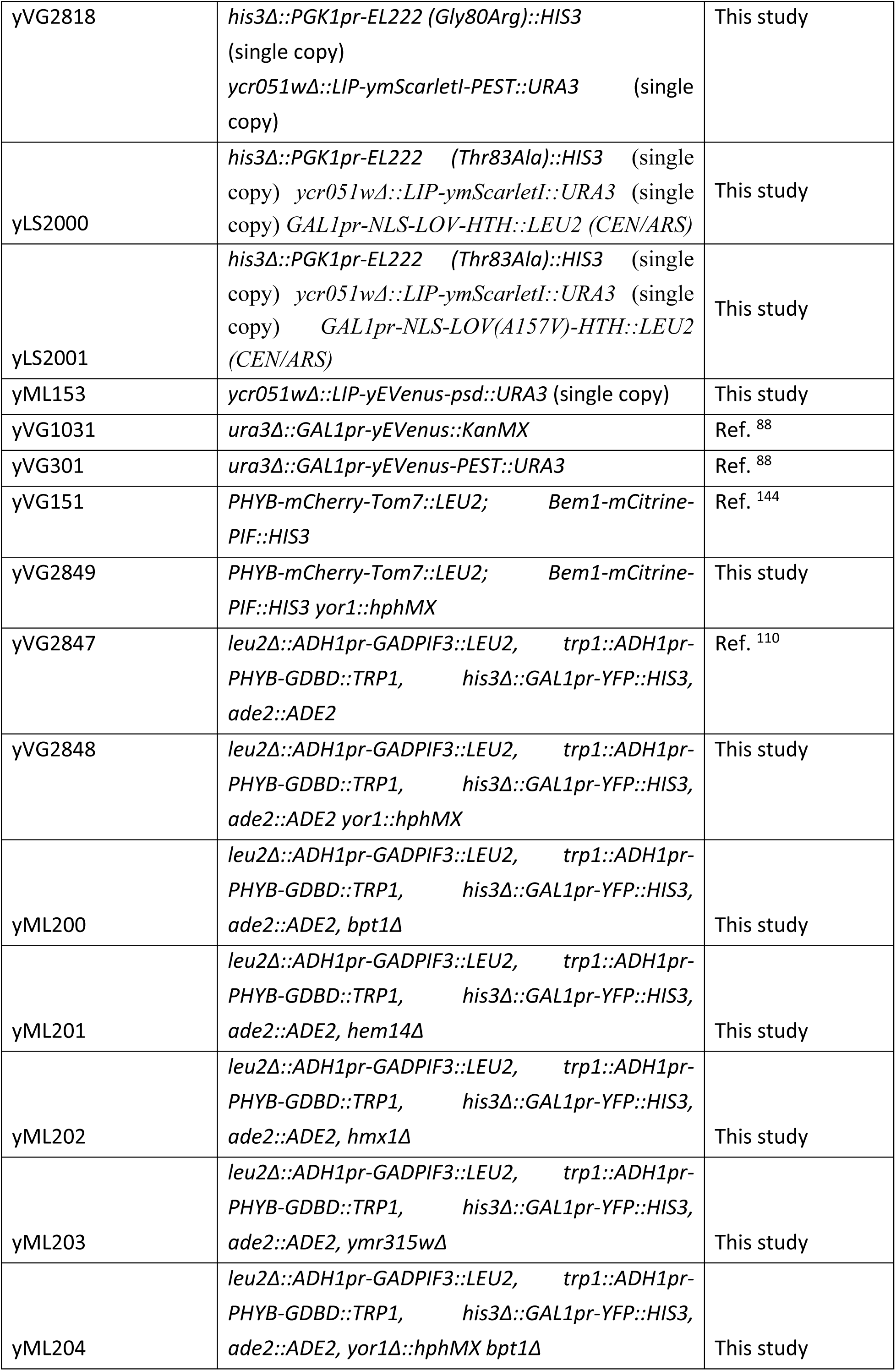
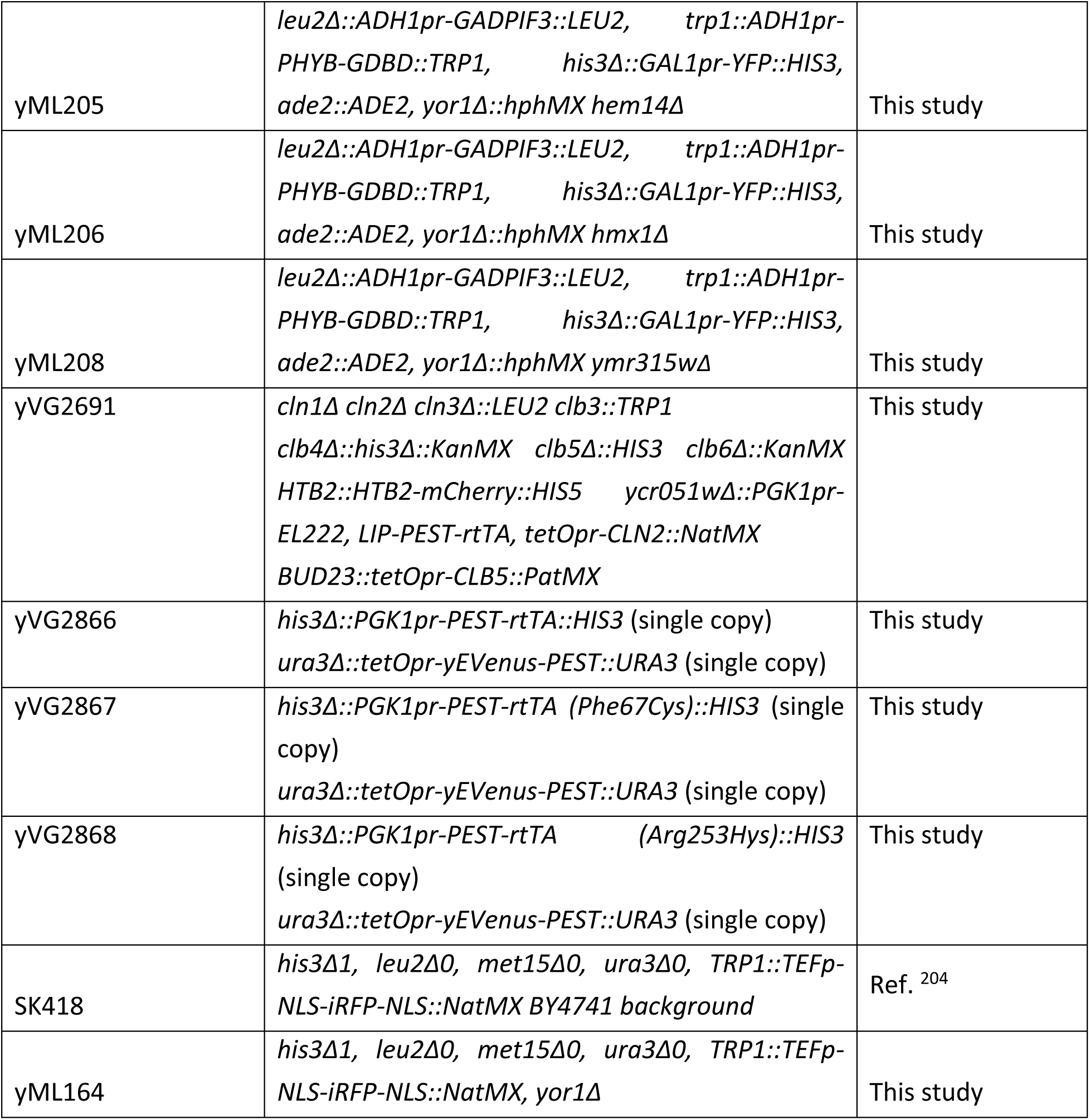
Strains used in the study.

**Supplementary Table 3.**
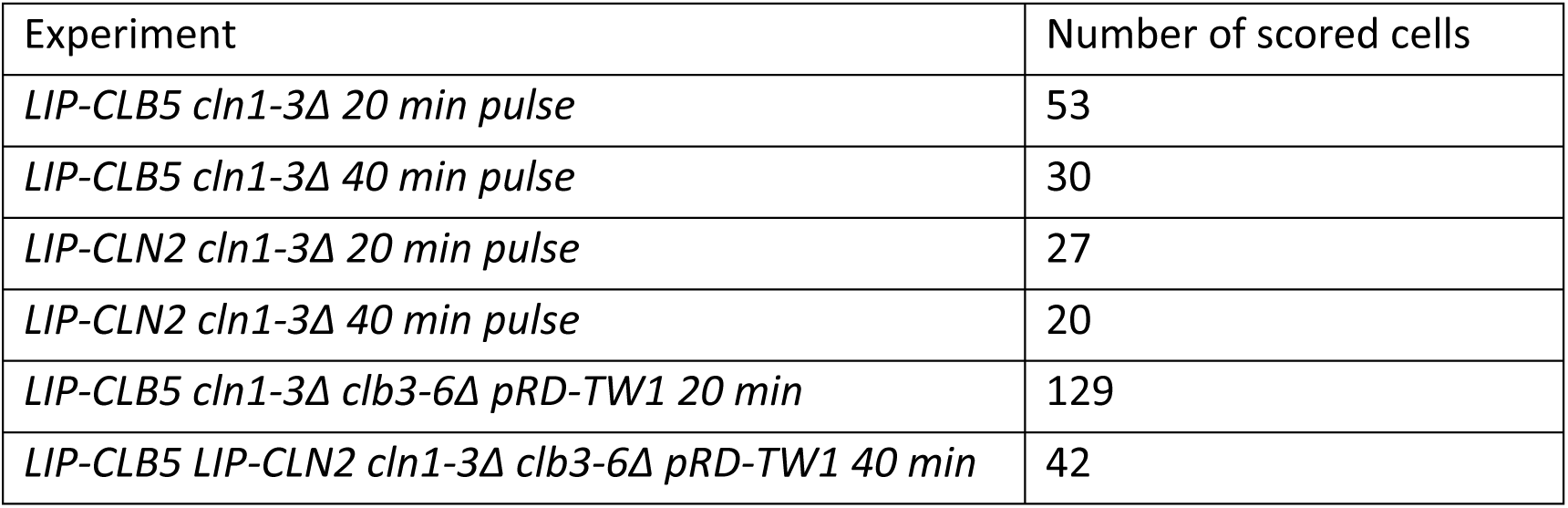
Number of cells analyzed in Fig. 2 B.

**Supplementary Table 4.**
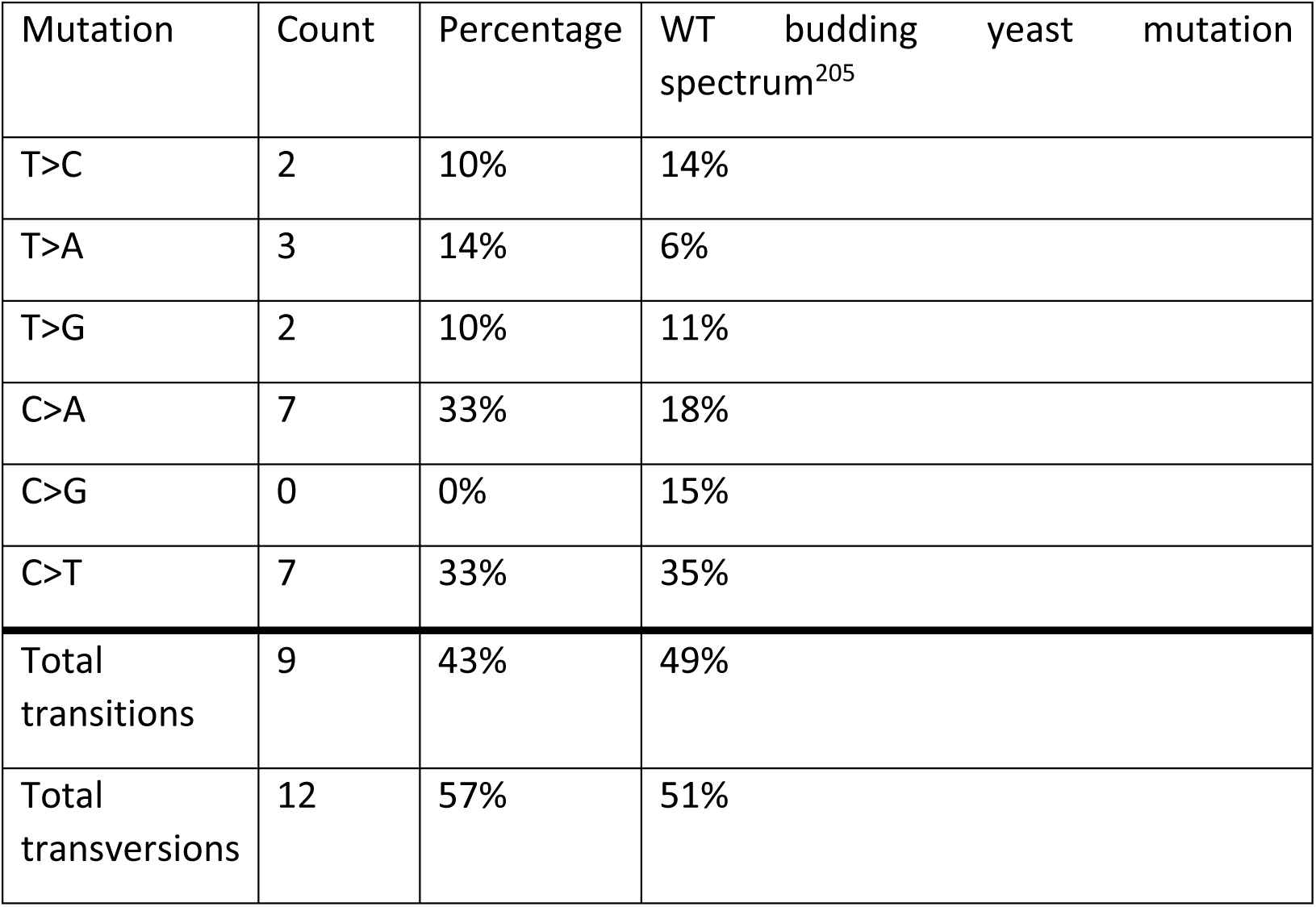
Distribution of mutation types across campaigns 1-3 for *EL222* compared to the WT budding yeast mutation spectrum from ref. ^205^

**Supplementary Table 5.**
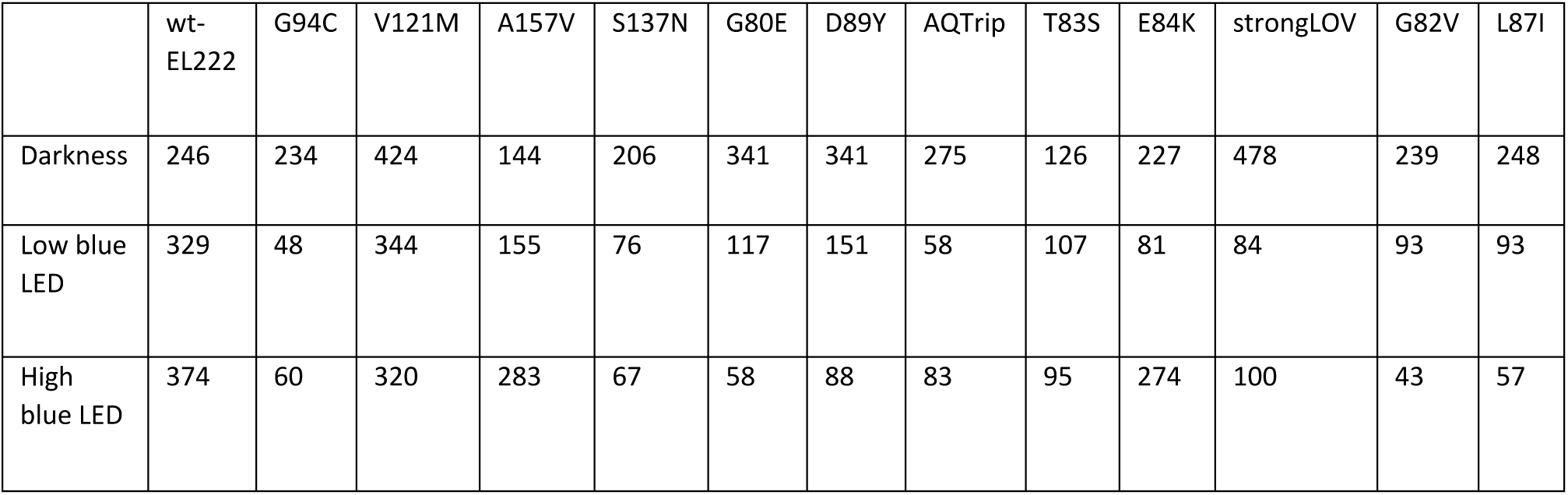
Number of cells analyzed in Fig. 5 B.

**Supplementary Table 6.**
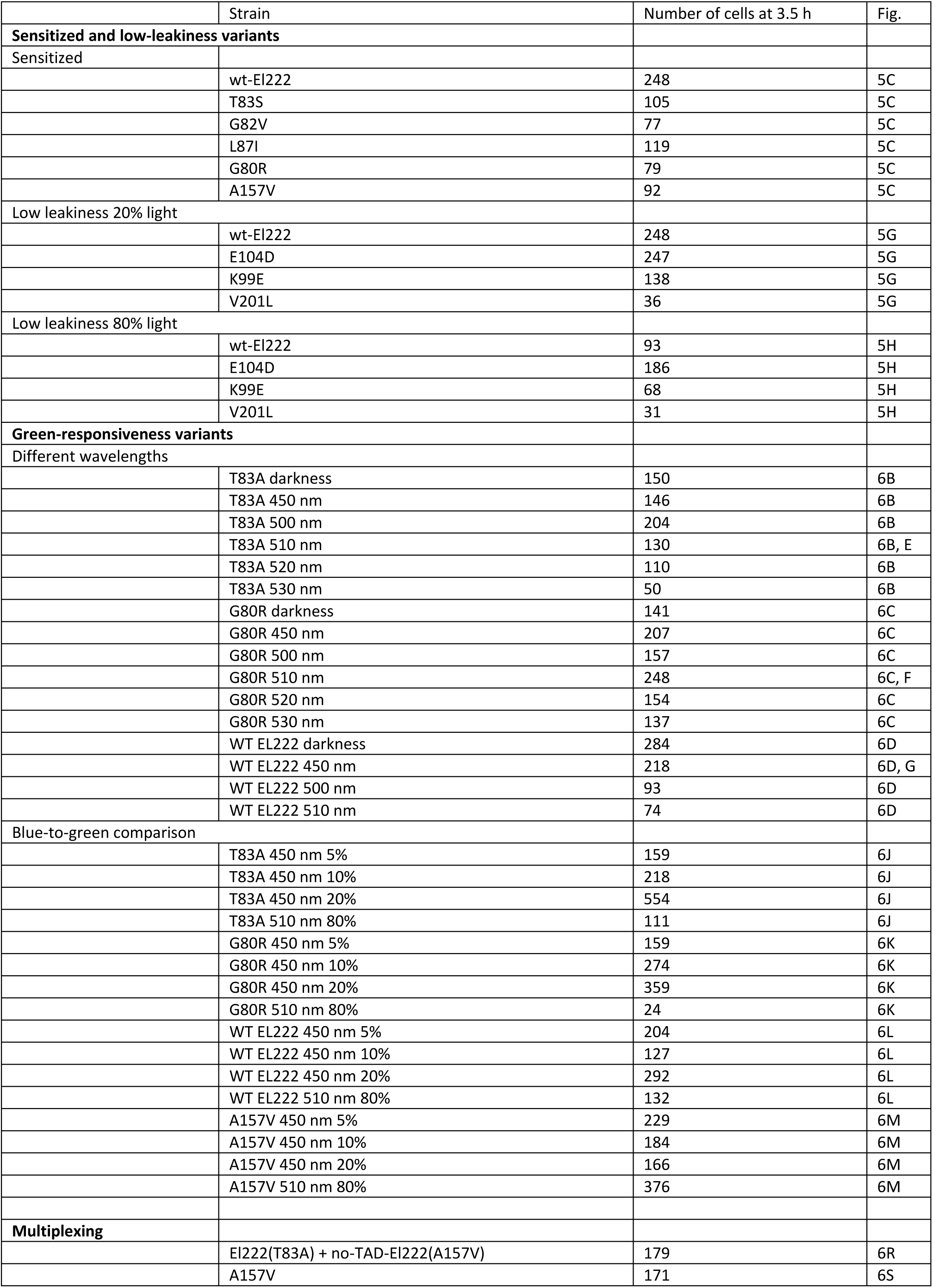
Number of cells analyzed in Fig. 5 C, G, H and Fig. 6 B-G, J-M, R-T.

**Supplementary Table 7.**
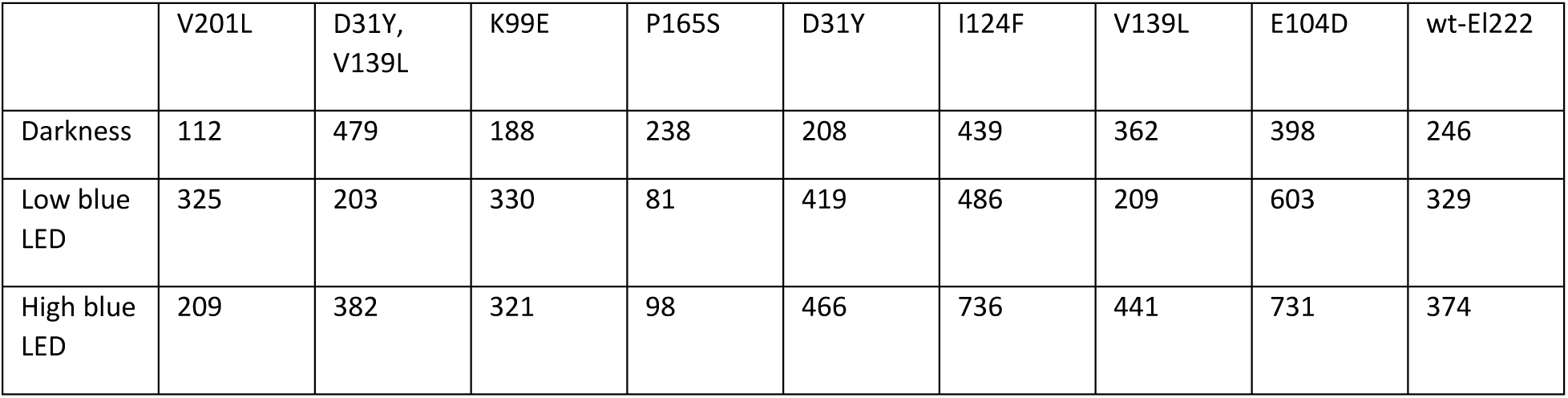
Number of cells analyzed in Fig. 5 F.

**Supplementary Table 8.**
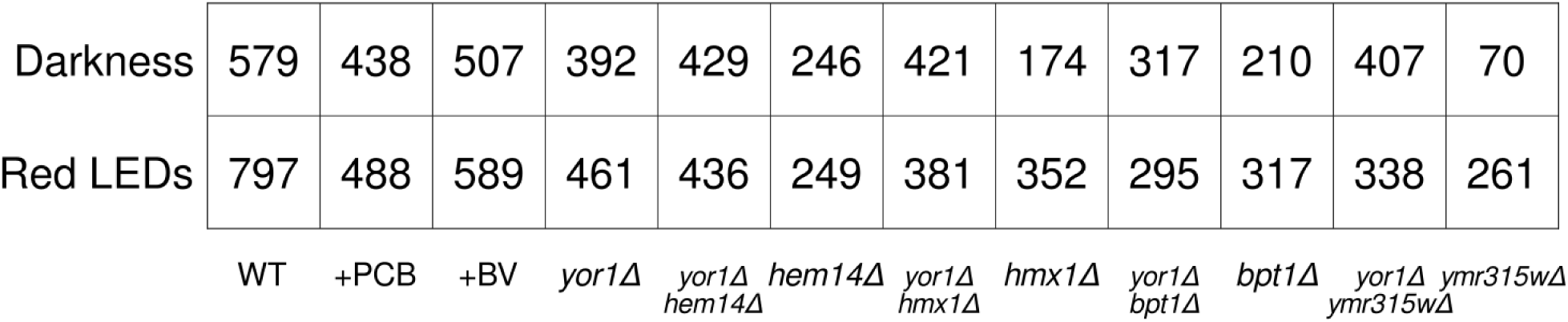
Number of cells analyzed in Fig. 7 C.

**Supplementary Table 9.**
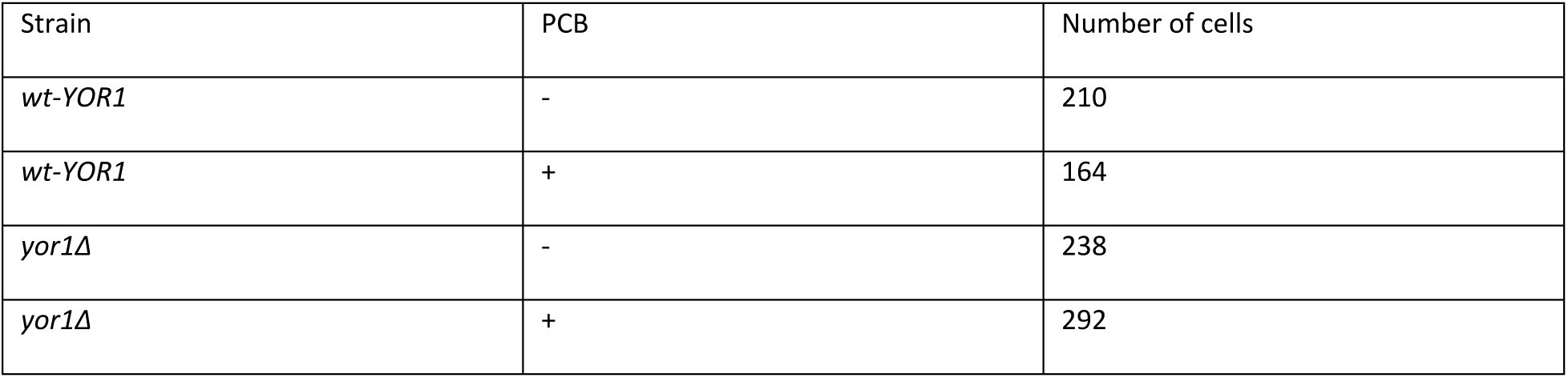
Number of cells analyzed in Fig. 7 F, G.

**Supplementary Table 10.**
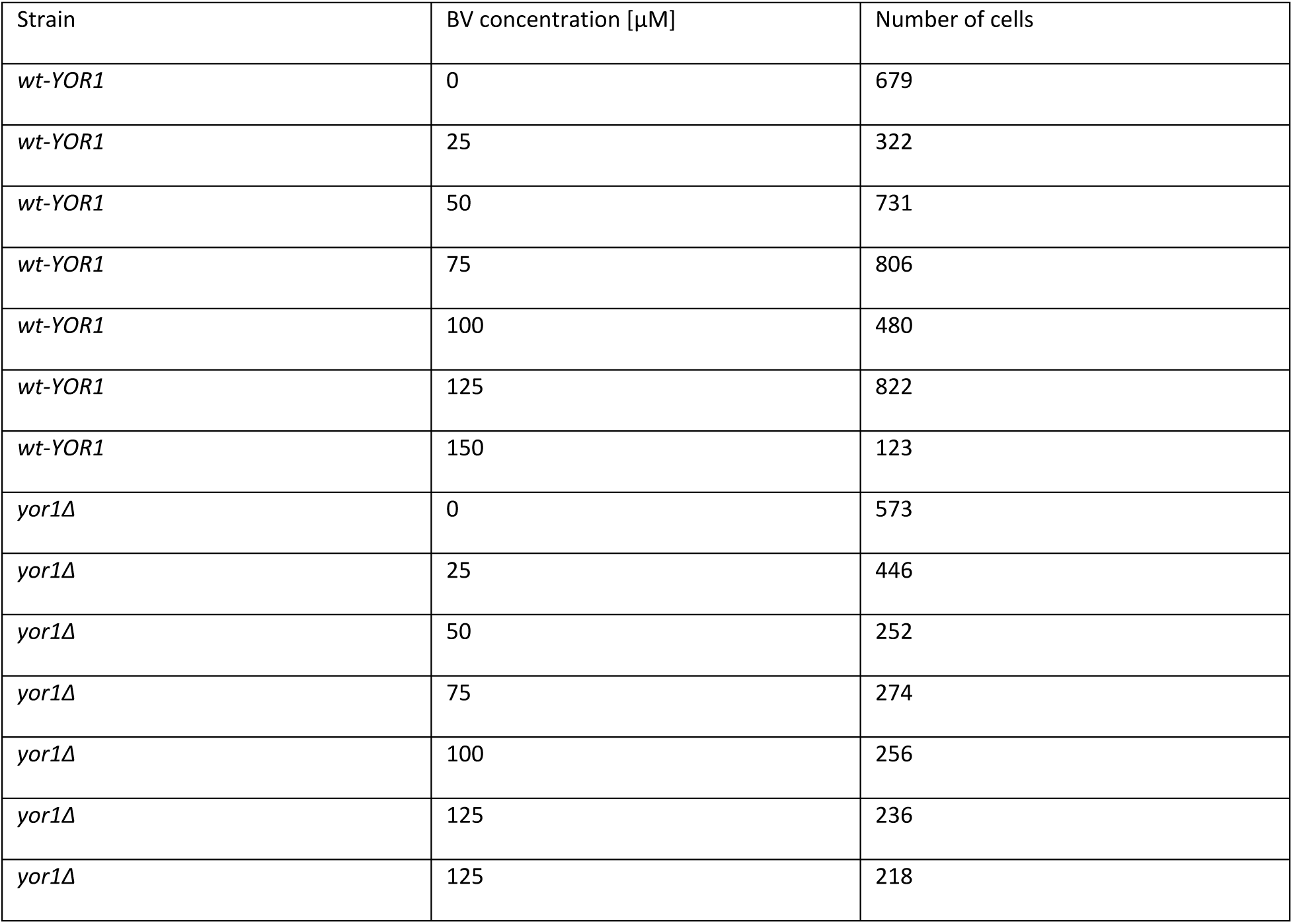
Number of cells analyzed in Fig. 7 H.

**Supplementary Table 11.**
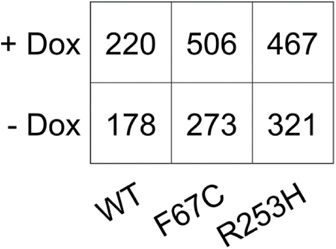
Number of cells analyzed in Fig. 8 D.

**Supplementary Table 12.**
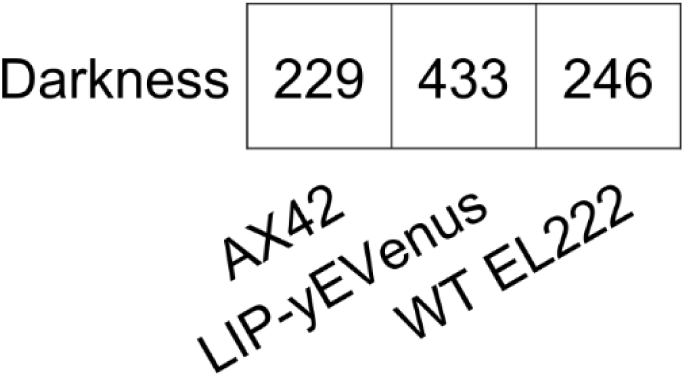
Number of cells analyzed in Supplementary Figure 3.

## Notes

### Summary of Updates

Several additions regarding El222 and PhyB-Pif3 campaigns and characterization of the results.

